# Fine tuning of CpG spatial distribution with DNA origami for improved therapeutic cancer vaccination

**DOI:** 10.1101/2022.06.08.495340

**Authors:** Yang C. Zeng, Olivia J. Young, Christopher M. Wintersinger, Frances M. Anastassacos, James I. MacDonald, Giorgia Isinelli, Maxence O. Dellacherie, Miguel Sobral, Haiqing Bai, Amanda R. Graveline, Andyna Vernet, Melinda Sanchez, Kathleen Mulligan, Youngjin Choi, Thomas C. Ferrante, Derin B. Keskin, Geoffrey G. Fell, Donna Neuberg, Catherine J. Wu, David J. Mooney, Ick Chan Kwon, Ju Hee Ryu, William M. Shih

## Abstract

Multivalent presentation of ligands often enhances receptor activation and downstream signaling. DNA origami offers precise nanoscale spacing of ligands, a potentially useful feature for therapeutic nanoparticles. Here we introduce a “square block” DNA origami platform to explore the importance of spacing of CpG oligonucleotides, which engage Toll-like receptors and thereby act as danger signals for dendritic cells. Through *in vitro* cell-culture studies and *in vivo* tumor-treatment models, we demonstrate that square blocks induce Th1 immune polarization when CpG is spaced at 3.5 nm. We observe that this DNA origami vaccine enhances DC activation, antigen cross-presentation, CD8 T cell activation, Th1-polarized CD4 activation and NK cell activation. The vaccine also synergizes effectively with anti-PD-L1 for improved cancer immunotherapy in melanoma and lymphoma models and induces long-term T cell memory. Our results suggest that DNA origami may serve as an advanced vaccine platform for controlling adjuvant spacing and co-delivering antigens.

**One Sentence Summary:** This study developed a DNA origami-based cancer vaccine (DoriVac) that co-delivers antigen and CpG immune adjuvant with an optimal nanospacing for Th1 immune polarization.

## Introduction

Therapeutic personalized cancer vaccine technologies have made tremendous strides in recent years ^1^. Successful vaccination requires immune activation via engagement of pattern-recognition receptors such as Toll-like receptors (TLRs) expressed by antigen-presenting cells (APCs). One class of TLR ligands is CpG oligodeoxynucleotides (termed CpGs here) containing unmethylated cytosine-phosphate-guanosine present in bacteria and viruses ^2, 3^. Recognition of CpGs by TLR9 in APCs can be exploited to boost the antigen-specific immune response ^3^. Nanoparticles that display multiple copies of CpGs represent promising candidates for cancer vaccine adjuvants ^4, 5^. However, there is limited knowledge of how specific spatial arrangements of CpGs presented on nanoparticles could be optimized to influence the magnitude, duration, and polarization of immune responses. Such information could prove to be critical for the design of future vaccines targeting cancer or infectious diseases.

The nanoscale distribution of ligands on therapeutic nanoparticles is believed to impact which signaling pathways are activated in targeted cells, as subtle differences in ligand spacing can be translated into diverse cellular responses ^6–8^. The adjuvant CpG is well known to provoke differential Th1 or Th2 immune responses, depending on the context of CpG presentation ^9^. In particular, the spacing between multiple CpGs presented by therapeutic nanoparticles may determine immune polarization. The crystal structure of CpG-bound TLR9 revealed that two CpG molecules bind to dimeric TLR9 such that they are spaced 2.9 nm apart ^10^. In one study, poly (lactic-co-glycolic acid) (PLGA) nanoparticles decorated with a relatively high density of CpGs were found to induce a Th1-polarized immune response, while nanoparticles decorated with a lower density provoked a Th2-polarized immune response ^11^. Another study suggested that CpGs at an optimum spacing of 3.5 nm can interlock with multiple TLR9s like a zipper, resulting in amplified immune responses ^12, 13^; however, these CpGs were presented in a double-stranded DNA context, whereas TLR9 is thought to bind to single-stranded DNA^10^. Most recently, CpG dimers were fabricated on DNA origami with 7 nm and 38 nm spacing ^14^, where only the former strongly activated RAW264.7 cells. Another recent study also found that both CpG copy number and spatial organization could contribute to the magnitude of TLR9 signaling when CpG was fabricated on a wireframe DNA origami structure ^15^. Overall, these studies indicate that the spacing of CpG at the nanoscale level can impact receptor activation and subsequent immune polarization. For cancer therapy, a Th1-polarized APC response would be beneficial as it would lead to Th1-polarized CD4 and cytotoxic CD8 T lymphocyte activation and secretion of antitumor cytokines ^16^. In addition, nanoparticles with optimally spaced CpGs could enable significant Th1 immune polarization with a minimal adjuvant dose, potentially providing the means for reduced adjuvant-associated side effects ^17^. However, most of the currently investigated nanomaterials (e.g., those not based on DNA origami) cannot provide uniform CpG spacing, and instead only control the average spacing, which may lead to mixed induction of Th1 and Th2 responses.

DNA origami has been explored for cancer therapy due to its properties of rigid compact structure, controlled drug release, and precise stoichiometric loading with multiple cargos ^18–21^. DNA origami can be used to display precise nanoscale arrangements of ligands ^14, 15, 22, 23^ such as CpGs and co-deliver tumor antigens. Using Watson-Crick complementary base-pairing, a long single-stranded DNA (ssDNA) scaffold and hundreds of short staple strands can self-assemble into custom three-dimensional (3D) nanoscale shapes ^24–28^. CpGs can be incorporated into specific locations on the surface of DNA origami via conjugation to specific staple strands. Antigens can be co-delivered on the DNA origami for improved presentation and cross-presentation ^29–31^. Methods have been developed for stabilization of DNA nanostructures against low-salt denaturation and nuclease degradation *in vitro* or *in vivo* via electrostatic formulation with PEGylated oligolysine ^32, 33^. Furthermore, DNA origami itself is well tolerated when administered systemically ^34, 35^. In this study, we introduce a square block DNA origami (SQB; see below) to investigate how CpGs with finer-tuned nanospacings (2.5 to 7 nm) than what has been previously explored on DNA origami can be exploited to induce a Th1-polarized immune response (Fig. 1A, B) for enhanced vaccine efficacy. We refer to our DNA origami vaccine — a combination of SQB with CpG and antigen attachments — as DoriVac. We fabricated DoriVac using either model antigen or neoantigens and investigated the efficacy of DoriVac monotherapy and combination therapy on several murine tumor models.

**Figure 1.**
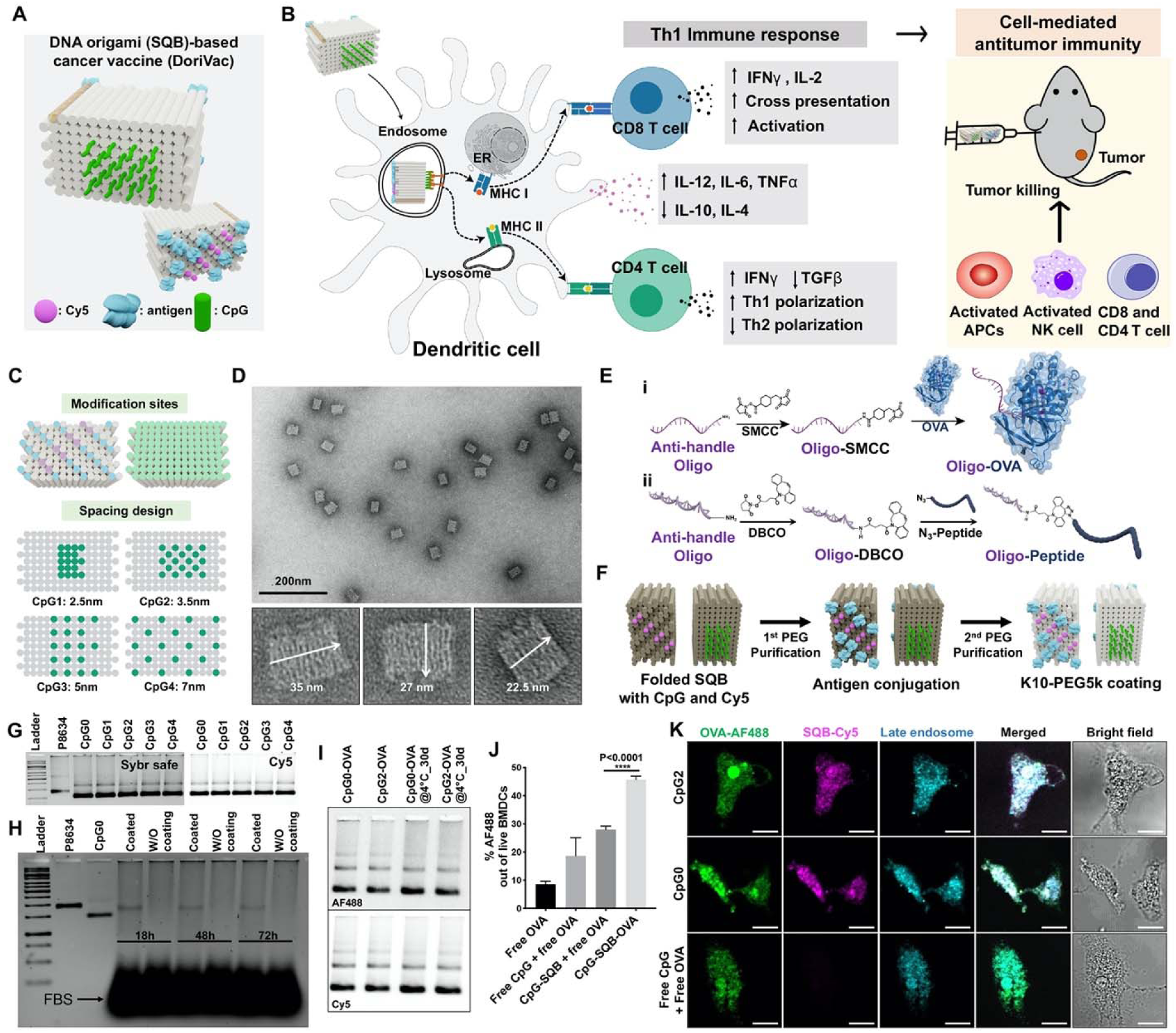
DNA origami square blocks (SQBs)-based vaccines (DoriVac) were fabricated with different spacings of CpG adjuvant. (**A**) Proposed schematic of the DNA origami-based cancer vaccine (DoriVac) containing CpG (green) as adjuvant, OVA protein (blue) as model antigen and Cy5 dye (pink) as tracer. (**B**) Schematic figure showing DoriVac co-delivering antigen and adjuvant at optimal spacing induces improved Th1-polarized immune response. First, DoriVac is internalized into dendritic cells (DCs) and the CpGs on the vaccine nanoparticles interact with TLRs at endosome membrane. Second, DCs are differentially activated depending on CpG spacing, in the optimal case secreting more Th1-polarizing cytokines such as IL-12, IL-6, and TNFα, but less Th2-polarizing cytokines IL-10 and IL-4. Finally, DoriVac with optimized CpG configuration is administered to tumor-bearing mice and demonstrates enhanced anti-tumoral cellular immune responses. (**C**) The modification sites and CpG array on DNA origami SQB. Green dots: CpG modification sites. Pink dots: Cy5 modification sites. Blue dots: antigen (e.g. OVA, neoantigen) modification sites. Different CpG (green dots indicate the positions for a single CpG strand) spacings were designed to arrange CpGs on specific helix ends of the SQB. More information is shown in Fig. S4. CpG1, 2, 3, and 4 refer to four versions of DoriVac with different CpG spacings. (**D**) Representative TEM images of the 3D SQB nanostructure. White arrows indicate dimensions of SQB. (**E**) Schematic figure showing (i) conjugation strategy between OVA and anti-handle via an SMCC linker. The maleimide group of the SMCC-modified anti-handle (purple wavy line) can be conjugated to four cysteine residues of OVA. (ii) Conjugation strategy between neoantigen (short peptide) and anti-handle via azide-DBCO click chemistry. (**F**) Schematic of the DoriVac composition, purification with polyethylene glycol (PEG) and coating with PEGylated oligolysine (K10- PEG5k). Two steps of PEG purification enable removal of excess staple DNA strands and excess anti-handle conjugated OVA. K10-PEG5k coating promotes DNA origami resistance against low-salt denaturation and nuclease degradation. (**G**) Agarose gel results showing the SQB folded with different CpG spacing imaged for SYBR Safe (left) and Cy5 (right) (no antigen included). (**H**) Agarose gel results of SQB with and without K10-PEG5k coating after incubation with 10% FBS culture medium for 18, 48 and 72 hours. (**I**) Agarose gel results of DoriVac (CpG0-OVA, CpG2-OVA) freshly prepared or stored at low temperature (4 °C) for 30 days. Model antigen OVA is conjugated with Alexa Fluor 488 (AF488) and SQB is attached with Cy5. (**J**) Quantification on uptake of DoriVac by bone marrow-derived DCs (BMDCs). Model antigen OVA is conjugated with Alexa Fluor 488 (AF488) and SQB is attached with Cy5. DoriVac is treated at 1 nM SQB or equivalent amounts of free CpG and/or OVA as loaded on the SQB for the controls. n=5. (**K**) Confocal images of BMDCs treated with DoriVac for 1 day. The scale bar represents 10 µm.

## Results

### 1. DNA origami square blocks (SQBs)-based vaccines (DoriVac) were fabricated with different spacings of CpG adjuvant

SQB DNA origami has a rigid structure and a high density of potential modification sites, allowing the ligands to be spaced in increments as fine as 2.5 nm. Compared to other nanoparticles conjugated with CpGs, CpG-functionalized DNA origami can be additionally modified with other cargos (i.e., antigen for co-delivery) without changing the CpG spatial distribution. Meanwhile, by optimizing the spacing and stoichiometry of the adjuvant, the dose can be tailored to maximize efficacy while minimizing adjuvant toxicity ^17^. We designed the folding of SQB with CaDNAno software by coupling an M13-based ssDNA scaffold (8634 nucleotides; Table S1) with hundreds of complementary staple strands (Table S2) ^36^ (Fig. S1–3). There are 126 double helices in total, allowing facile modification on both ends of the helices, where ssDNA docking handles can be programmed for precise, addressable spacing of cargos. One face of the SQB is flat, while the other face has extruding helices (Fig. 1C). These structures are monodispersed with dimensions of 35 × 22.5 × 27 nm, and exhibit minimal aggregation as evidenced by TEM imaging (Fig. 1D, Fig. S3).

We explored various spatial patterns of CpGs on the flat face of the SQB (Fig. 1C, Fig. S4). Staple strands directly functionalized with CpGs were first incorporated into SQB during initial folding; folded particles were purified with polyethylene glycol (PEG; Fig. 1E, F) precipitation. Eighteen phosphorothioate-modified CpG strands (20 mer) were introduced into SQB with precisely controlled distances of 2.5, 3.5, 5 and 7 nm between CpGs, corresponding to CpG1, CpG2, CpG3, and CpG4 (Table S3, Fig. S5-10). All CpGs on SQB were placed in the same orientation with the 5’ end extruding from the SQB (Fig. 1G, Fig.S4B), except that to achieve 2.5 nm spacing in CpG1, a mixture of ten 5’ and eight 3’ extensions were used, or else a 3’-3’ linkage was created via DBCO-azide click chemistry to preserve the 5’ orientation for all CpG extensions (see Methods, Fig. S5). We studied the CpG conjugation efficiency across all structures, consistently observing levels of over 90% incorporation for DoriVac designed with 18 CpGs and 80-90% incorporation for DoriVac design with over 18 CpGs (Fig. S10). CpG0, with no conjugated CpG strands on SQB, served as an origami control. To investigate the impact of CpG spacing in the context of complete cancer vaccine, the model antigen protein or neoantigen peptides can be modified onto the DNA origami through SMCC linkage or DBCO-Azide click chemistry linking to a hybridizing anti-handle oligo strand (Fig. 1E, Fig. S6, 7). The DoriVac fabrication underwent two steps of PEG purification and coated with PEGylated oligolysine (K10-PEG5k) before cell culture or animal studies to secure its stability against nuclease and low salt ^32, 33^. The SQB coated with K10-PEG5k could survive in 10% FBS culture medium for at least 72 hours (Fig. 1H). DoriVac remained monodispersed with minimal multimerization which was maintained stable and intact after storage at 4°C for one month (Fig. 1I).

To investigate the impact of CpG spacing on Th1-polarized immune responses, we set up sequential immune cell co-culture experiments and performed analysis by flow cytometry and enzyme-linked immunosorbent assay (ELISA; Fig. S11). In a leading cell culture study, we verified that SQBs (CpG0 – CpG4) could be taken up by HeLa cells, 293T cells, and mouse bone marrow-derived dendritic cells (BMDCs) (Fig. S12). We observed that BMDCs underwent greater apoptosis after exposure to free CpG alone compared to CpG-SQBs, indicating that free CpG is more cytotoxic in a non-antigen specific fashion than the same molar amount of CpG in SQBs (Fig. S13A–E). Interestingly, co-delivery of antigen and adjuvant by SQB significantly improved antigen uptake (Fig. 1J). DoriVac was observed to be readily taken up into BMDCs and to be colocalized with late endosomes in confocal imaging (Fig. 1K, Fig. S14A, Supplementary Video 1).

### 2. CpG, delivered at a spacing of 3.5 nm on DNA origami SQB, provides enhanced dendritic cell (DC) activation for Th1-polarized immune response

Next, we evaluated the CpG spacing effects on several DC cell types including mouse BMDC, human plasmacytoid DCs (pDCs), monocyte derived DCs (moDCs), and mouse RAW264.7 macrophages. Mouse mature BMDCs (indicated by CD86 and MHC LJ double-positive population) were significantly increased in the CpG2 group having 3.5 nm CpG spacing (Fig. 2A, Fig. S13G–J). CD40 can bind to the ligands on T helper cells as a costimulatory signal for Th1- polarized immune response ^37^. Moreover, DEC205 has been reported as a receptor of CpG on the cell surface and also a marker of activated DCs involved in antigen uptake ^38^. We found that the population of CD40^+^DEC205^+^ cells was greatly increased in all the DoriVac-treated groups, especially in the CpG2 group (Fig. 2B, Fig. S14B). We also observed the highest increase of SIINFEKL MHC I^+^ and TLR9^+^MyD88^+^ populations in the CpG2 group (Fig. S14C, D). By ELISA assay, we found that the DoriVac stimulated BMDCs to generate more Th1-polarizing cytokines including IL-12, TNFα and IL-6 (Fig. S14E, G, J, K), but fewer Th2-polarizing cytokines including IL-10 and IL-4 (Fig. S14F, H, L) compared to free CpG and free OVA administration (termed bolus vaccine). The marked increase in IL-12/IL-10 ratio indicated that CpG delivered at a spacing of 3.5 nm (CpG2) induced greater Th1 polarization compared to delivery through other spatial configurations (Fig. 2C, Fig. S13K–M, Fig. S14I) ^39–41^. Of note, the SQB alone (CpG0) generated only minimal IL-12 and IL-10 secretion, indicating the low immunogenicity of DNA origami (Fig. S14E-H). CpG2 consistently showed the strongest Th1 polarization on human pDCs and moDCs (Fig. 2D, E, Fig. S15). We note that moDCs showed more vigorous response to DoriVac stimulation compared to pDCs which corresponds to a recent study showing the CD11b^+^ myeloid cells preferentially internalize DNA origami (Fig. S15) ^34^. The spacing effects were further verified on the mouse RAW264.7 macrophages, which showed a significant increase of CD11c^+^, SIINFEKL MHC I^+^, and CD40^+^DEC205^+^ populations (Fig. 2F, G, Fig. S16A). In mouse RAW264.7 cells, stimulation by DoriVac was compared with liposome nanoparticles, which are widely used for vaccine formulation. Flow cytometry results showed that DoriVac significantly stimulated more CD11c, CD40, CD11b, PD-L1, and CD103 expression (Fig. 2H, I, Fig. S16B, C), and strikingly twelve times more of SINFEKL MHC I^+^ expression (Fig. 2J, Fig. S16D) on the RAW264.7 cells compared to liposomes carrying the same number of OVA and CpG.

**Figure 2.**
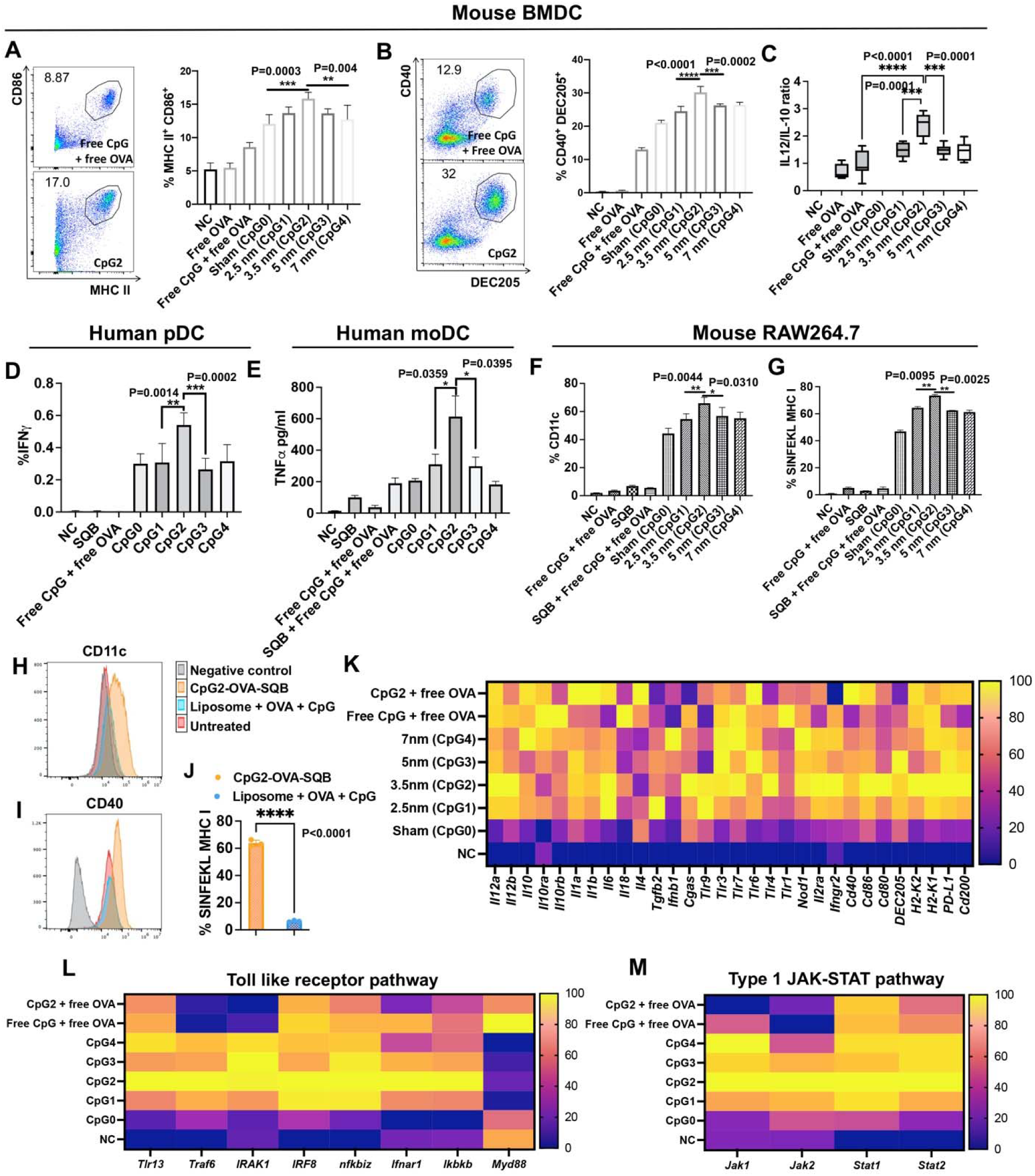
CpG, delivered at a spacing of 3.5 nm on DNA origami SQB, provides enhanced dendritic cell (DC) activation for Th1-polarized immune response. **(A-B)** Representative scatter plots (left) and percentages (right) of double-positive CD86^+^MHCII^+^ and CD40^+^DEC205^+^ populations detected in mouse BMDCs by flow cytometry. Numbers adjacent to outlined areas in the scatter plots represent percentage of cells in each subpopulation. (**C**) The ratio of IL-12 to IL- 10 detected by ELISA after one-day stimulation with various vaccine groups treated to mouse BMDCs. Significantly increased IL-12/IL-10 ratio observed for CpG2 treated BMDCs is consistent with Th1 polarization. (**D**) IFNγ expression in human plasmacytoid DCs (pDCs) treated with various vaccine groups as determined by flow cytometry. (**E**) TNFα secretion in human monocyte-derived DCs (moDCs) treated with various vaccine groups as determined by Luminex. (**F, G**) Percentages of CD11c^+^ and SINFEKL MHC I^+^ population detected in mouse RAW264.7 cell lines by flow cytometry, indicating similar CpG spacing effects on mouse RAW264.7 cell line as BMDCs. (**H, I**) CD11c and CD40 expression on RAW264.7 cells treated with DoriVac or liposome carrying the same amount of CpG and OVA. NC is unstained flow control. (**J**) Percentages of SINFEKL MHC I^+^ population detected in mouse RAW264.7 cells treated with DoriVac or liposome carrying the same amount of CpG and OVA by flow cytometry. (**K**) Heatmap of the expression patterns of BMDC genes across all the treated groups. Each row of the heatmap indicates a gene and each column indicates a sample. RNA sequencing revealed increased expression of Th1-polarization-related genes in the vaccine group when CpGs were spaced at 3.5 nm. In particular, Th1-polarizing cytokines *IL-12*, *IL-1*, and *IL-6* were notably increased in CpG2 group, compared to other spacings; Th2-polarizing cytokines *IL-10*, *IL-18*, and *IL-4* were downregulated compared to bolus vaccine. Toll-like receptors (*TLR9*, *TLR7*, and *TLR4*), DC maturation makers (*CD40*, *CD80*, *CD86*, and *DEC205*) and MHC I molecules (*H2- K1* and *H2-K2*) were most upregulated in CpG2 group, compared to all other treatment groups. The scale bar represents the normalized expression intensity. (**L, M**) Heatmap of the expression patterns of BMDC genes related to potential signaling pathways: Toll like receptor pathway and Type1 JAK-STAT pathway. NC represents negative control without treatment. Sham (CpG0) sham (CpG0) refers to SQB conjugated with OVA antigen but without CpG. n=5 for graphs A-G, n=3 for graph J. Various DNA origami vaccine constructs (based on the concentration of 1 nM SQB), free OVA (4 nM for A-C, K-M or 10 nM for D-J) or free CpG (18 nM) was treated.

To understand the gene profiling of immune polarization, we performed mRNA sequencing (RNAseq) analysis on the BMDC samples, which corroborated (Fig. 2K, Fig. S17) the ELISA results and demonstrated upregulation of *IL-12*, *IL-1*, and *IL-6* gene expression (Th1-polarizing cytokines) and downregulation of *IL-10*, *IL-4*, and *IL-18* gene expression (Th2-polarizing cytokines) in the context of CpG2 with 3.5 nm spacing compared to bolus vaccine and other spacing configurations. DC maturation markers (*CD40*, *CD80*, *CD86*, and *DEC205*), and MHC I molecules (*H2-K1* and *H2-K2*) were upregulated in the CpG2 group compared to others. Furthermore, we found that *TLR13* is significantly increased in DoriVac groups in the TLR signaling pathway (Fig. 2L. Fig. S17G). A previous study showed that chicken TLR21 is an innate CpG receptor distinct from human TLR9 ^42^, and showed similarity to mouse TLR13. We suspect that TLR13 might also be a receptor for CpGs delivered by SQB in our study, although a different study showed mouse TLR13 recognizes RNA ^43^. Based on the gene sequencing results, this pathway might not rely on MyD88 activation (Fig. 2L). It coincides with the previous study that TLR9 was activated by plasmid DNA even in MyD88-deficient mice ^44^. Some other pathways that may participate in DC activation include but are not limited to: Type1 JAK-STAT pathway (Fig. 2M, Fig. S17D), BMP-SMAD pathway (Fig. S17E), and STING pathway (Fig. S17F), based on the gene sequencing analysis. Of note, CpG2 showed the highest immune stimulation. Altogether, these data demonstrate that the spatial distribution of CpG has potent biological effects to direct Th1 polarization, and that adjuvant spacing can impact vaccine effectiveness.

### 3. T cell activation by DCs instructed by DoriVac bearing varying CpG spatial patterns and densities revealed distinctive anti-tumoral effects

We next queried whether antigen-specific CD4 and CD8 T cell responses could be triggered by DoriVac-pulsed DCs co-cultured with OVA-specific OT-I and OT-II T cells. Flow cytometry results revealed that both CD8 and CD4 T cell activation were increased by CpG2 (3.5 nm spacing) pulsed DCs (Fig. S18A–D). Interferon (IFN)γ-expressing Th1-polarized CD4 T cells were significantly increased in the CpG2 group when the antigen was co-delivered (Fig. 3A). IL- 2 secretion by CD4 T cells showed negligible differences among different spacings (Fig. S18E). TGFβ secretion by CD4 T cells was downregulated in all the DoriVac groups (Fig. S18F). IFNγ and IL-2 secreted by OT-I CD8 T cells were markedly increased in the CpG2 group (Fig. 3B, C), corresponding to increased proliferation of CD8 cells (Fig. 3D, E), while CpG-SQBs without co- delivery of antigen showed a similar trend but limited secretion of IFNγ and IL-2. In addition, CpG2 led to increased tumor cell killing by activated CD8 T cells compared to other spacings (Fig. 3F, Fig. S19). These data indicate that (1) spacing of CpG at 3.5 nm enhances the Th1- polarized immune response; and (2) simultaneous presentation of antigen and adjuvant through DoriVac strongly stimulates DCs and preferentially increases cross-presentation via MHC I for improved CD8 T cell activation.

**Figure 3.**
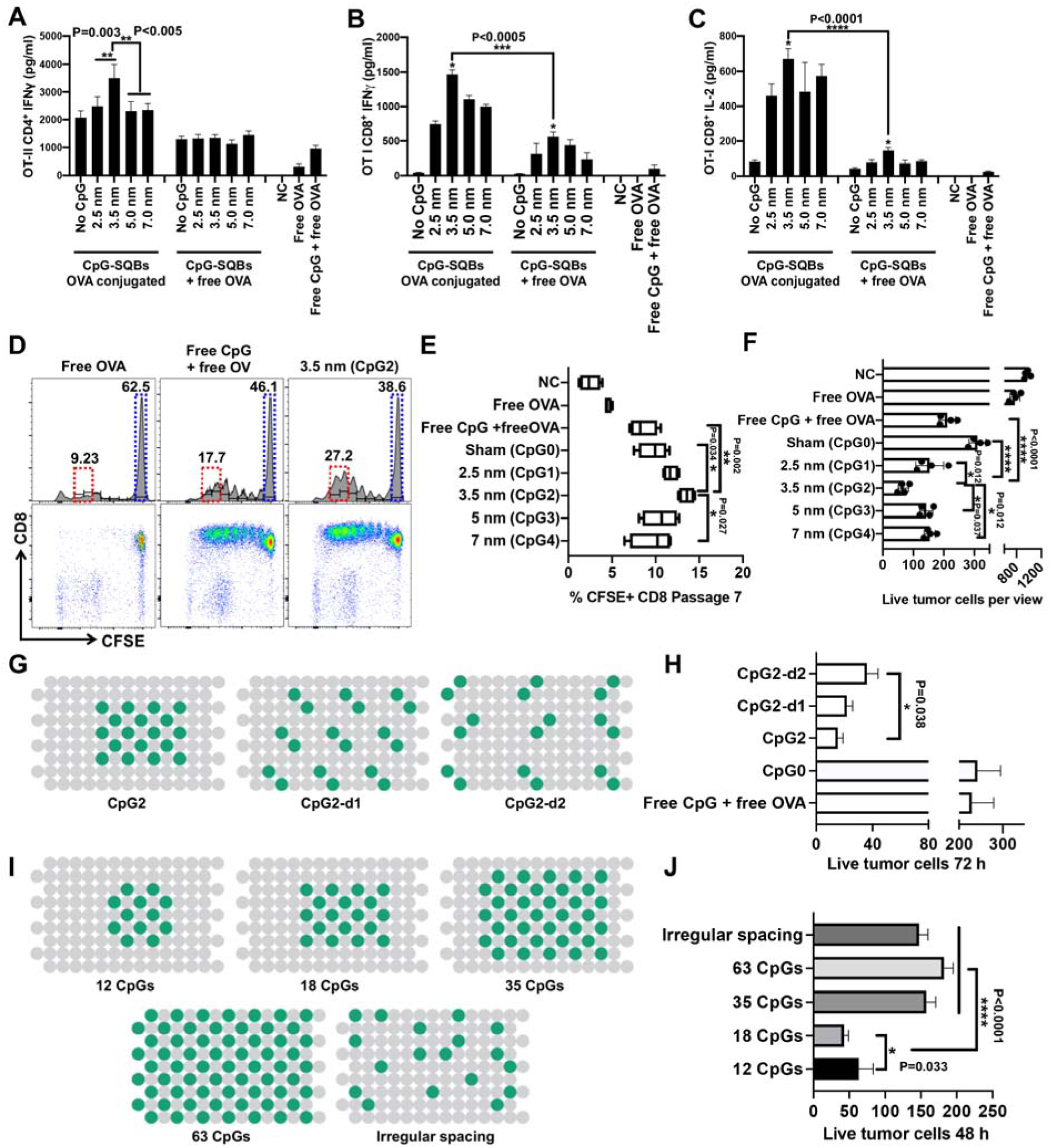
T cell activation by DCs instructed by DoriVac bearing varying CpG spatial patterns and densities revealed distinctive anti-tumoral effects. (**A, B**) Quantification of IFNγ in the co-culture supernatant of OT-II CD4 and OT-I CD8 T cells by ELISA. Co-delivery of antigens (left panels; CpG-SQBs OVA conjugated) could greatly induce cytokine secretion compared to freely delivered antigens (right panel; CpG-SQB + free OVA). SQB with 3.5 nm spacing of CpG and conjugated antigen (CpG2) enhanced co-delivery to DCs and greatly induced IFNγ secretion compared to all other groups. (**C**) Quantification of IL-2 in the co-culture supernatant of OT-I CD8 T cells by ELISA, indicating improved cross-presentation and CD8 T cell proliferation. (**D**) Representative scatter plots of CFSE fluorescent dye labeled OT I CD8 T cells in three groups. The proliferation of CD8 T cells was greatly enhanced with CpG spacing of 3.5 nm as indicated by increased populations of passage 6 and 7 cells (red frame). Non-proliferating cells (blue frame) were correspondingly decreased. (**E**) The frequency of OT I CD8 T cells at passage 7. (**F**) Quantification of live tumor cells after 48 hours co-culture of OT I CD8 T cells activated with various DoriVac constructs and B16-OVA melanoma cells. Activated OT I CD8 T cells were co-cultured with tumor cells in a 10:1 ratio. (**G**) Various design with different spacings of CpG dimer on SQB. Two CpG oligonucleotide strands were placed as a dimer unit and distributed with three different spacings (CpG2, CpG2-d1, CpG2-d2). The gray dots represent the double helices of the DNA origami, and the green dots indicate the positions for a single CpG strand. (**H**) Quantification of live tumor cells after 72 hours co-culture of OT I CD8 T cells activated with various DoriVac constructs and B16-OVA melanoma cells. Activated OT I CD8 T cells were co-cultured with tumor cells in a 10:1 ratio. (**I**) Various design with different CpG densities on SQB. CpGs were spaced at 3.5 nm, but the number of CpGs varied from 12 to 63. A design with 18 irregularly spaced CpGs was tested to verify the effects of precise spacing. (**J**) Quantification of live tumor cells after 48 hours co-culture of OT I CD8 T cells activated with various DoriVac constructs and B16-OVA melanoma cells. Activated OT I CD8 T cells were co-cultured with tumor cells in a 10:1 ratio. The tumor killing was greatly increased in 18 CpGs group, corresponding with the distinct CD8 T activation. NC represents negative control without treatment. n=5 for graphs A-C, E, H, and J, n=4 for graph F. Various DNA origami vaccine constructs (based on the concentration of 1 nM SQB), free OVA (4 nM for A-F, 6 nM for G-J) or free CpG (18 nM) was treated.

A previous study showed that when two CpGs were conjugated together as a dimer at the 3’ end, the DC stimulation could be enhanced ^45^. This study motivated us to explore whether a spatial configuration of CpG dimers might affect receptor multimerization. To this end, we conceptualized adjacent CpGs (with 3.5 nm spacing between the two monomers) as a dimer unit and spaced them apart on the SQB with decreasing densities (CpG2-d1 and CpG2-d2 in Fig. 3G). Interestingly, we found that both IL-12 and IL-10 were elevated as spacing of the dimer increased even though we did not observe an associated change in the IL-12/IL-10 ratio (Fig. S20A–C). However, CD8 and CD4 T cell activation (indicated by IFNγ secretion) decreased as the dimer spacing increased (Fig. S20D, E), and the tumor-killing effects showed a corresponding decrease (Fig. 3H, Fig. S20K–M). These data suggest that spatial configuration of CpGs could impact receptor activation and subsequent immune polarization through receptor multimerization. Furthermore, DNA origami may provide an informative tool to further study the role of multimerization in TLR activation and other receptor activation.

Another study previously demonstrated that increasing the molar ratio of CpG on a DNA duplex induced increasing secretion of IL-12 ^46^. To investigate if the number of CpGs influences the immune polarization, we kept constant the CpG 3.5 nm spacing, antigens per DNA origami particle, and number of particles, but only increased the number of CpGs per particle from 12 to 63 (Fig. 3I). Irregularly spaced CpGs (CpGi) were also designed as a control with 18 CpGs. IL- 12 and IL-10 secretion both increased when there were more CpGs (Fig. S20F, G). However, IL- 10 secretion was greatly induced with 35 and 63 CpGs (Fig. S20G). The IL-12/IL-10 ratio was higher with 12 or 18 CpGs than with 35, 63, or irregularly spaced CpGs (Fig. S20H). We found that 12 to 18 CpGs showed no difference in terms of IFNγ secretion from CD4 OT-II T cells, but 12 CpGs induced less IFNγ secretion from CD8 OT-I T cells (Fig. S20I, J). However, 35, 63, and irregularly spaced CpGs induced less IFNγ expression in both CD4 OT-II and CD8 OT-I T cells, consistent with elevated IL-10 leading to inhibition of CD8 T cell function ^47^. The tumor-killing effects were most notable in the 18 CpGs group (Fig. 3J, Fig. S20N). These data suggest that both optimal CpG stoichiometry and uniform CpG spacing may be beneficial for Th1 immune response.

### 4. DoriVac distribution, *in vivo* immune-cell stimulation and prophylactic-vaccination effects

To explore the potential of DoriVac as a vaccine, its biodistribution was first evaluated through *in vivo* fluorescence imaging. DoriVac primarily accumulated in the nearest draining lymph nodes (LNs) from the injection site with minimal increase in other LNs (Fig. 4A, Fig. S21). The DoriVac persisted in the draining LN for at least 48 hours and the antigen maintained high intensity even after 48 hours as evidenced by AF488 conjugated on OVA (Fig. 4A, B). The SQB was cleared by both the liver and kidney within two days, as the Cy5 fluorescent signal was barely detectable at this time point (Fig. 4A-C). Lung and spleen only showed minimal Cy5 signal increase at two and four hours (Fig. 4C). Administration of the DoriVac in naïve mice verified that optimized spatial configurations (CpG2; 3.5 nm spacing and 18 CpGs) could greatly increase DC activation and improve the CD8^+^ tetramer^+^ population without increasing the immunosuppressive regulatory T cells (Tregs) (Fig. 4D–H, Fig. S22). Encouraged by the preferential Th1 immune response, two doses of the DoriVac treatment were subcutaneously injected on days 0 and 7 as a prophylactic treatment, and then the mice were inoculated with B16-OVA tumor cells on day 14 (Fig. 4I). On day 28, measurable tumors were observed in control groups and in 40% of the bolus vaccine group (Fig. 4J). No tumors were observed in DoriVac-treated group until day 42. While all mice in the untreated group and 60% of mice in the bolus vaccine group died by the conclusion of the study, only one mouse in the CpG2 group died but with a much longer life span compared to other groups (Fig. 4K). These results verified the prophylactic efficacy of the DoriVac in an aggressive murine melanoma model.

**Figure 4.**
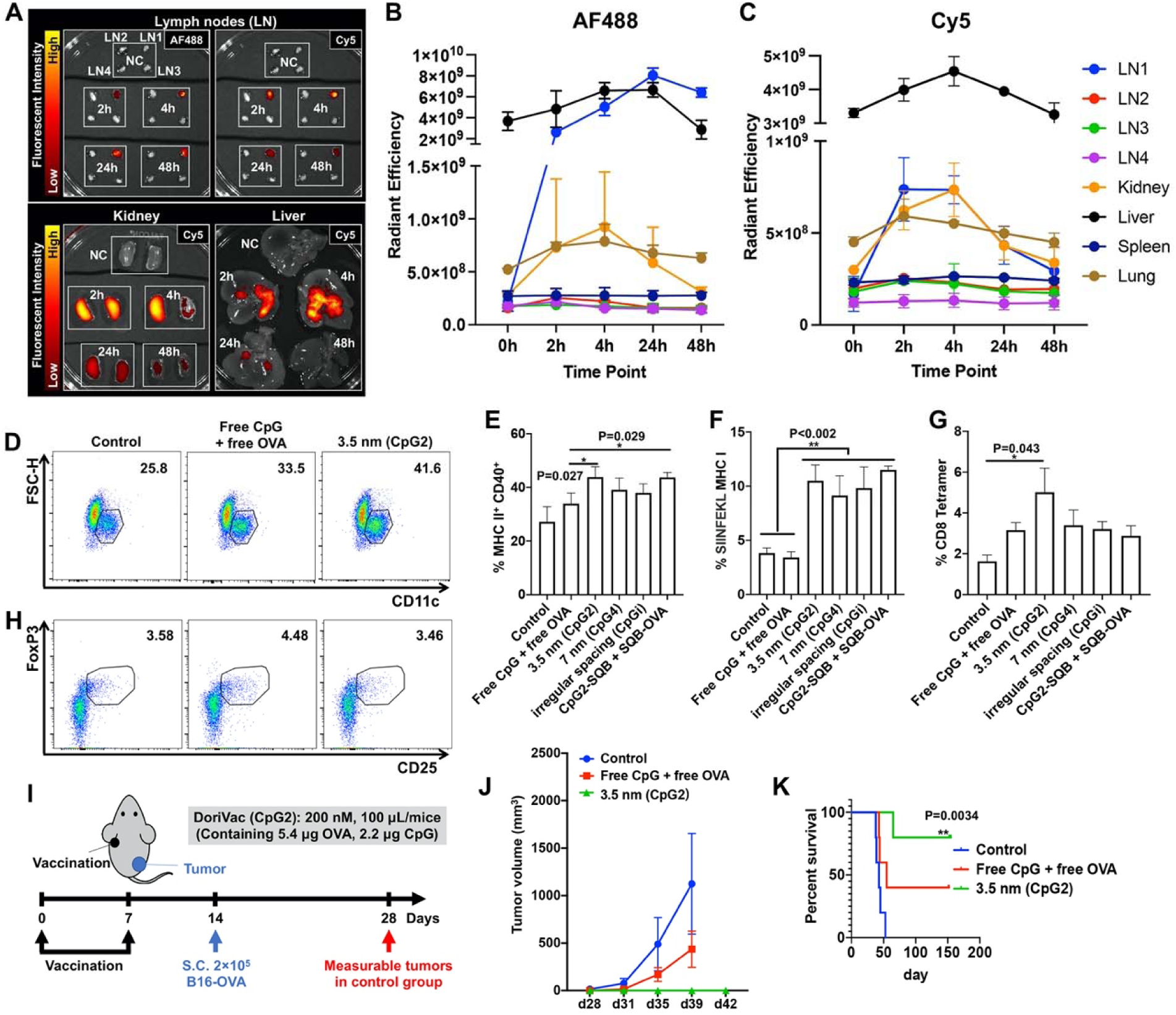
DoriVac distribution, *in vivo* immune-cell stimulation and prophylactic- vaccination effects. (**A**) Representative *ex vivo* images showing the organ distribution of the DoriVac within 48 hours after subcutaneous injection on left shoulder analyzed by IVIS machine detecting the fluorescence carried on the vaccine nanoparticles. The images showed the AF488 (OVA) or Cy5 (SQB) in different organs. The nanoparticles primarily accumulated in the draining lymph node and were cleared by both liver and kidney. LN1: left brachial; LN2: right branchial; LN3: left inguinal; LN4: right inguinal. (**B, C**) Radiant efficiency of AF488 and Cy5 in different organs corresponding to *in vivo* accumulation, noting that different organs may have different baseline fluorescent readings. (**D-H**) Before applying the vaccine in a tumor model, we vaccinated naïve C57BL/6 mice and detected DC activation and T cell response in the draining lymph node on day 1 and day 8 post vaccination. n=5 **(D)** Representative scatter plots showing CD11c^+^ DC cells in the lymph nodes on day 1 post vaccination. Increased DC activation is observed in CpG2 applied group. Numbers adjacent to outlined areas in the scatter plots represent percentage of cells in each subpopulation. (**E**) Flow results showing MHC II^+^ CD40^+^ population on DCs. (**F**) Flow results showing SIINFEKL MHC I expression on DCs. (**G**) Flow results showing CD8 T cell population bound with SIINFEKL MHC I tetramer on day 8 in the lymph node. Only CpG2 greatly induced tetramer^+^ CD8 T cell population on day 8. When CpG and OVA was delivered separately by SQBs (CpG2-SQB + SQB-OVA), the DC maturation markers and cross-presentation markers showed a demonstrated increase. However, the tetramer^+^ CD8 T cell population was not increased when CpG and OVA were delivered separately. (**H**) Representative scatter plots showing CD4^+^CD25^+^FoxP3^+^ Treg cells in the lymph nodes on day 1 after vaccination. The DNA origami vaccine did not increase the population of Tregs, in contrast to the bolus vaccine (free CpG and free OVA) which increased the Treg population. (**I**) Schematic showing prophylactic vaccination treatment plan and tumor challenge protocol in C57BL/6 mice. Mice were vaccinated on day 0 and 7 and then inoculated with 2 × 10^5^ B16-OVA tumor cells on day 14. (**J**) Tumor growth graph. Note that only one mouse in the DoriVac-treated group demonstrated tumor growth, after a remarkably delayed tumor onset (4 weeks after tumor inoculation). (**K**) Mouse survival curve (n=5). Control refers to untreated group. Two out of five mice survived from the bolus vaccine (free CpG + free OVA) group, whereas four out of five mice survived from the origami vaccine (CpG2) group. ** refers to comparison with the control.

### 5. Immune cell profiling revealed a Th1-polarized immune response after therapeutic DoriVac treatment in mouse melanoma models

We next evaluated therapeutic antitumor effect of DoriVac. A melanoma mouse model was established by subcutaneous injection of B16-OVA cells in the flank of mice, after which we treated the tumor-bearing mice three times, each with 10, 20, 40, or 80 pmoles of DoriVac with optimized spatial configurations of CpG (CpG2). The results showed that 10 or 20 pmoles DoriVac inhibited tumor growth and increased survival time more efficiently (Fig. S23A, B). Increasing the doses did not result in further improvement in efficacy. Of note, 20 pmoles of DoriVac contain only 5.4 µg OVA and 2.2 µg CpG, significantly less than other pre-clinical CpG studies ^48^. When 20 pmoles of DoriVac with different spacings or the corresponding bolus controls were applied, CpG2 with optimal CpG configuration (i.e. 3.5 nm spacing) significantly delayed tumor growth compared to other spacing configurations (Fig. 5A, B, Table S5). The median survival of CpG2 DoriVac (26 days) was higher compared to other groups with median survival between 18 and 21 days (Fig. 5C, Fig. S23C, D). Anti-CD8 antibody treatment abolished the therapeutic effects observed in the context of CpG2, suggesting that the DoriVac strongly relies on CD8 activation (Fig. 5B, C). Next, immune cells from the LNs and tumor tissues were analyzed after three treatments (Fig. S24A–C). We observed significantly increased DC accumulation in the LN, and improved DC maturation and cross-presentation, decreased myeloid-derived suppressor cells but increased monocytes in CpG2 group (Fig. 5D, Fig. S24D– J). Th1-polarized CD4 and CD8 T cells were activated more in CpG2 group compared to bolus vaccine (Fig. S25). We observed that 3.5 nm spacing induced significantly increased CD8 activation compared to other spacings (7 nm or irregular spacing) (Fig. S25A, C). In the tumor tissue, infiltrated CD3 T cells accumulated notably in the CpG2 vaccine-applied group (Fig. 5E, Fig. S26A–D). CpG2 group showed increased effector memory cells and effector cells in both CD4 and CD8 T cell populations (Fig. S26E–H). A large portion of the infiltrated CD8 T cells were OVA-specific in CpG2-treated group, significantly more than the CpG4 group where CpG spacing was at 7 nm (Fig. 5F). More IL-2^+^ CD8^+^ and CD4^+^ T cells were found in CpG2 group as well (Fig. 5G). The Tregs, myeloid-derived suppressor cells, and M2 macrophages in the tumor tissue were not increased in the DoriVac-treated groups with a further decreased trend in CpG2 group (Fig. 5H, I, Fig. S26I-M). Furthermore, 35 CpGs (3.5 nm spacing) did not show enhanced therapeutic effect compared to 18 CpGs (CpG2, 3.5 nm spacing), most likely due to the decreased CD8 activation (Fig. S27. Fig. S20F–J). These results indicate that DoriVac CpG2, with optimal CpG configurations (3.5 nm spacing and 18 CpGs) directed a Th1-polarized immune response in the treated tumor through a cohort of immune-cell regulation.

**Figure 5.**
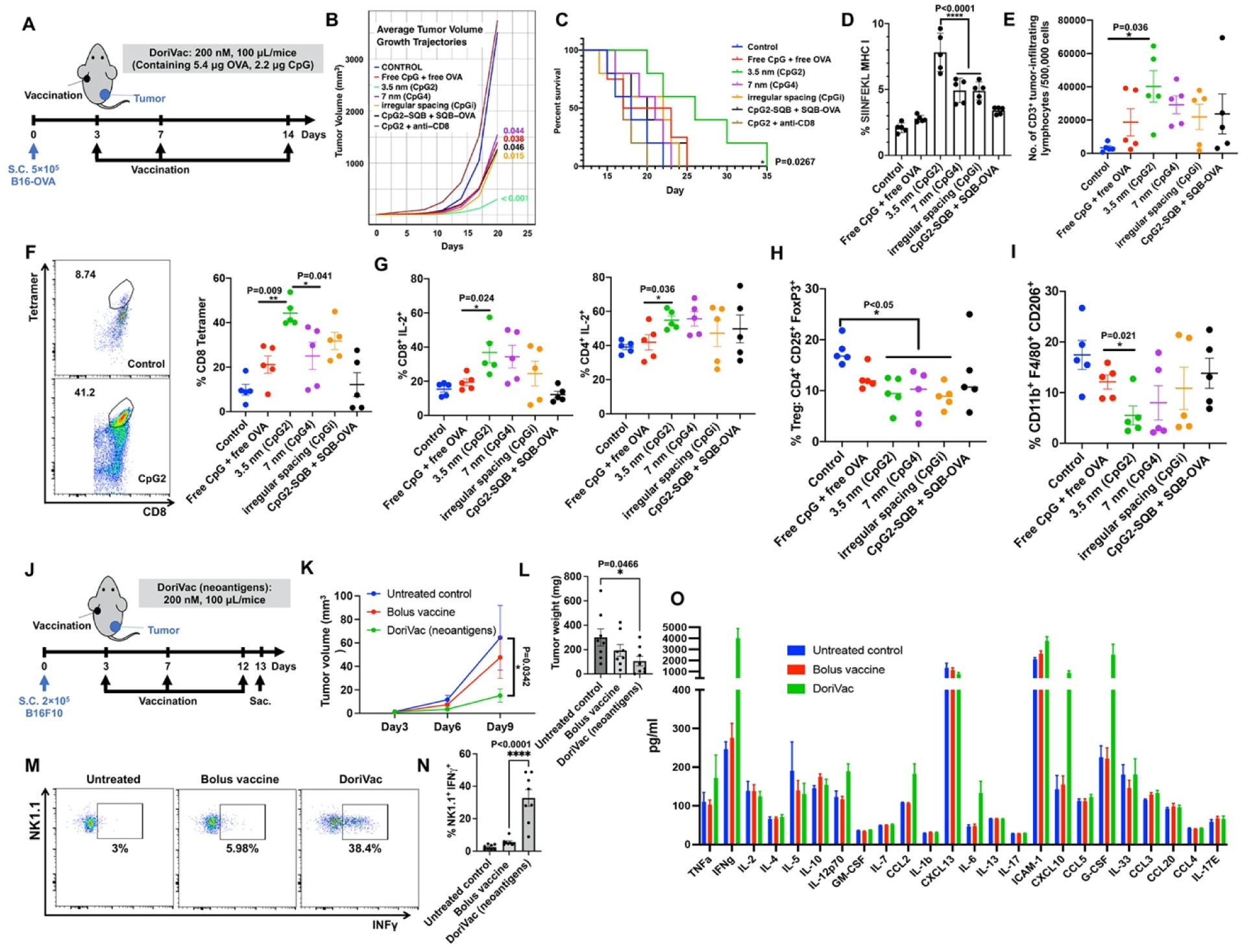
Immune cell profiling revealed a Th1-polarized immune response after therapeutic DoriVac treatment in mouse melanoma models. (**A**) Schematic delineating mouse tumor model setup and therapeutic vaccination treatment plan in C57BL/6 mice. Mice were inoculated with 5 × 10^5^ B16-OVA tumor cells on day 0 and treatment was administered on day 3, 7 and 14. (**B**) Average tumor volume growth trajectories (n=5). (**C**) Mouse survival curve (n=5). Median survival of CpG2, CpG4, CpGi, free CpG+ free OVA, CpG-SQB+SQB-OVA, and untreated control was 26, 21, 22, 20, 20, and 18 days, respectively. CD8 antibody was administered intraperitoneally (i.p.) every other day for three times, starting the day before vaccination. * refers to comparison with the control. To analyze immune cell profiling, B16- OVA tumor-bearing C57BL/6 mice were treated with various vaccines on days 3, 7 and 11, and then sacrificed on day 15. The draining lymph nodes (D) and tumor tissue (E-I) were processed to single cells and analyzed by flow cytometry. (**D**) Percentages of SIINFEKL MHC I^+^ cells in the CD11c^+^ population in draining lymph node. CpG2 with 3.5 nm spacing showed the most significant increase in CD11c^+^ DCs presenting SIINFEKL MHC I. (**E**) Number of intratumoral CD3^+^ tumor-infiltrating lymphocytes (TILs) out of 500,000 cells in various treatment groups. (**F**) Representative scatter plots (left) and percentages (right) of SIINFEKL MHC I tetramer^+^ cells in intratumoral CD8^+^ T cells. CpG2 group had significantly more tetramer^+^ cells compared to other CpG spacings. (**G**) Representative IL-2^+^ scatter plots on CD8^+^ cells (left) and percentages of IL- 2^+^ cells in CD8^+^ and CD4^+^ T cells (right). The difference of IL-2 expression between the CpG2 and bolus group was more notable in CD8 T cells. (**H, I**) Quantification of CD25^+^FoxP3^+^CD4^+^ Treg cells and CD11b^+^F4/80^+^CD206^+^ immunosuppressive type 2 macrophages population in tumor tissue. All the origami vaccine groups induced downregulation of Treg cells. The immunosuppressive type 2 macrophages population were also downregulated in the CpG2 group. (**J**) Schematic delineating mouse tumor model setup and therapeutic vaccination treatment plan with DoriVac bearing neoantigen in C57BL/6 mice. Mice were inoculated with 2 × 10^5^ B16F10 melanoma cells on day 0 and treatment with three vaccine groups was administered on day 3, 7 and 12. Untreated control (naïve mice), Bolus vaccine (Free neoantigen peptides + Free CpG), DoriVac (neoantigen). The mice were sacrificed (sac.) on day 13 for immune cell profiling. (**K**) B16F10 tumor growth graph (n=8). (**L**) B16F10 tumor weight on day 13. (**M, N**) Representative scatter plots and percentage of double-positive IFNγ^+^NK1.1^+^ populations detected in NK1.1^+^CD3^-^ population by flow cytometry. **(O)** Cytokine expression profile of the plasma samples analyzed by multiplexed Luminex after sacrificing the mice on day 13. Numbers adjacent to outlined areas in the flow plots represent the percent frequency of cells in each subpopulation.

The therapeutic anti-tumor effect of DoriVac was further assessed against neoantigens (Fig. 5J). Neoantigens are considered ideal vaccine targets as they are not expressed in normal tissues. However, neoantigen peptides often demonstrate poor immunogenicity, necessitating an adjuvant-bearing vaccine system. For targeting neoantigens in the murine B16F10 melanoma model, DoriVac was fabricated with B16F10 neoantigen peptides, M27 and M33 (CD8^+^ responsive) and M30 and M47 (CD4^+^ responsive)^48, 49^ (Fig. S28). DoriVac bearing neoantigen induced significant tumor suppression compared to untreated control with one mouse in the DoriVac group showing no tumor after three treatments on day 3, 7 and 12. (Fig. 5K, L, Fig. S29A). Additionally, DoriVac could significantly increase NK cell activation detected by IFNγ expression (indicated by NK1.1^+^IFNγ^+^) without obvious increase in the NK cell number in the LN (Fig. 5M, N, S29B-E). Additional immune cell profiling on DC and T cells demonstrated similar activation phenotypes as shown in Fig. 4 and Fig. S24 and 25 (data not shown). The plasma was analyzed by multiplex Luminex for cytokine profiling after sacrificing the mice on day 13. We observed massive increase of Th1-polarized cytokines in the blood of DoriVac-treated groups such as TNFα, IFNγ, IL-12, CCL-2, IL-6, CXCL10, and G-CSF, but no increase of Th2-polarized cytokines in the DoriVac-treated groups was detected compared to bolus vaccine or untreated groups (Fig. 5O). The cytokine level returned to baseline in one week after the final treatment as tested in another DoriVac experiment (data not shown). We also evaluated the anti-dsDNA antibodies and anti-PEG antibodies. Our results disclosed that the increase of anti-dsDNA antibodies is associated with effective anti-tumor effects ^50^ instead of DNA origami nanoparticles themselves (Fig. S30A, B). There is also no significant elevation of anti-PEG IgG observed in mouse serum (Fig. S30C). These results indicated DoriVac may serve as a safe, effective, and modular platform for presentation of various neoantigens to induce therapeutic anti-tumor efficacy.

### 6. DoriVac combined with immune checkpoint inhibitor anti-PD-L1 exhibited synergistic, durable T cell responses

Since we detected increased PD-L1 expression on DCs in the draining LN of the DoriVac-treated groups (Fig. 6A), we hypothesized that combining the DoriVac with immune checkpoint inhibition would further enhance the therapeutic effects (Fig. 6B). Indeed, the combination with αPD-L1 (but not αPD-1) significantly inhibited tumor growth and induced tumor regression in 4 out of 5 mice on B16-OVA melanoma model (Fig. 6C, D, Fig. S31). All 4 mice with tumor regression survived and showed no sign of recurrence (Fig. 6E). All the surviving mice were rechallenged 4 months later with 2 × 10^5^ B16-OVA tumor cells, resulted in complete tumor remission (Fig. S31D). The enzyme-linked immune absorbent spot (ELISpot) assays performed one year after the initial tumor cell inoculation revealed that only DoriVac-included groups could induce SIINFEKL-specific CD8 T cell activation (Fig. S31E). Other than B16-OVA model, DoriVac presented enhanced therapeutic efficacy compared to bolus vaccine groups in EG7- OVA lymphoma tumor model (Fig. S32A). Meanwhile, all the mice survived from the DoriVac treatment in combination with αPD-L1 (Fig. 6F). ELISpot results on the splenocytes and peripheral blood mononuclear cells (PBMCs) of the survived mice at 6 months disclosed the long-term durable CD4 and CD8 T cell memories in the DoriVac-applied groups (Fig. 6G, H, Fig. S32B-D). These results demonstrate that DoriVac could induce both innate and adaptive immune cell activation with a durable T cell response. In addition, the antigen-specific immune response induced by DoriVac synergized with immune checkpoint inhibition for improved cancer immunotherapy.

**Figure 6.**
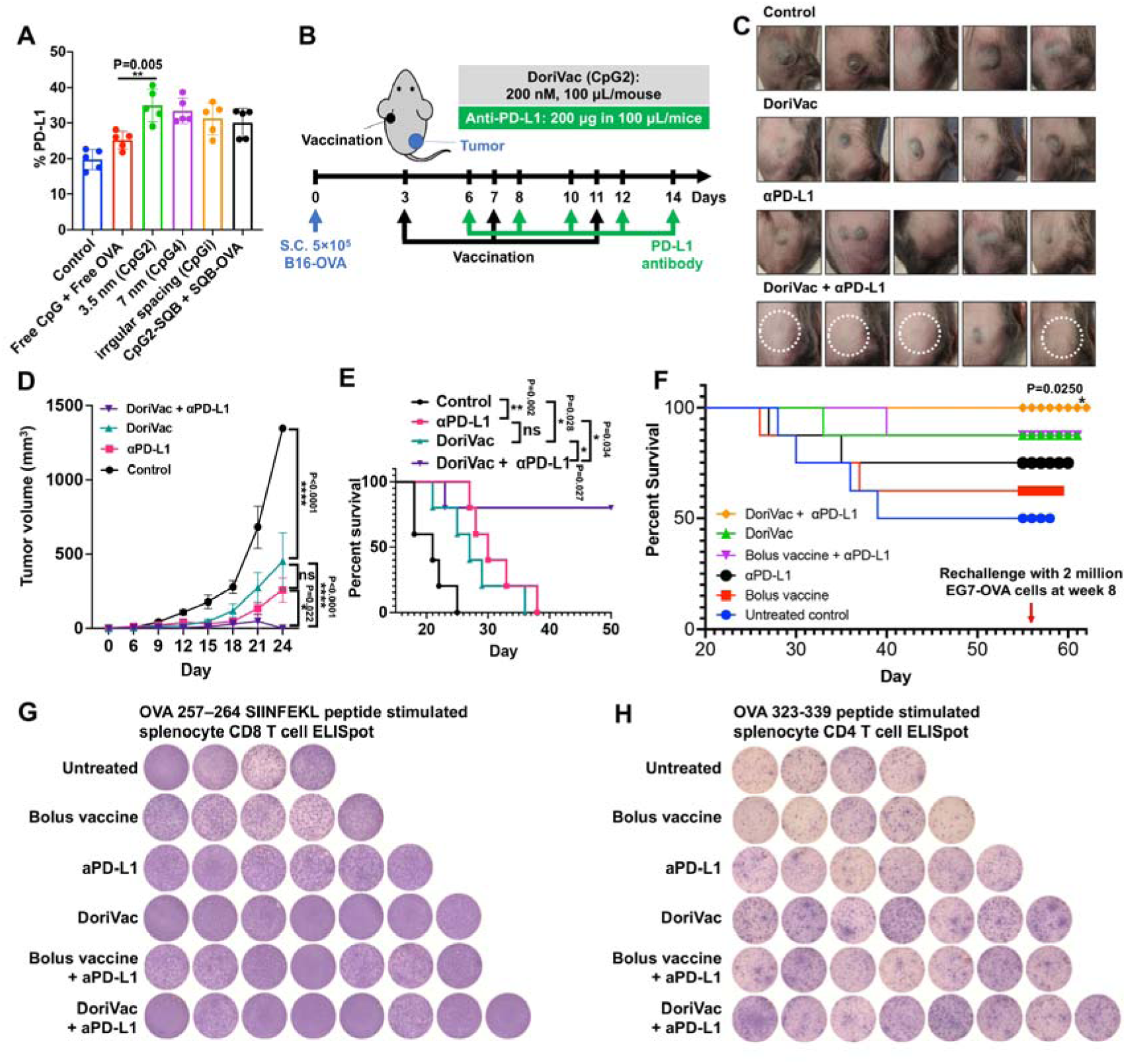
DoriVac combined with immune checkpoint inhibitor anti-PD-L1 exhibited synergistic, durable T cell responses. (**A**) Percentages of PD-L1^+^ cells on dendritic cells (DCs) in the lymph node. PD-L1 expression was increased on the DCs in the DNA origami vaccine-treated groups, correlating with the increase in the activated DC population, which provided a rationale for combination with anti-PD-L1 immune checkpoint inhibitor. (**B**) Experimental design schematic for the combination treatment with anti-PD-L1 antibody (αPD-L1). CpG2 DoriVac (200 nM) was subcutaneously administrated on days 3, 7, and 11 and anti-PD-L1 (200 µg/mice) was administered on the days 6, 8, 10, 12, and 14 to B16-OVA tumor-bearing mice. (**C, D**) The tumor photos (white circle indicates faded tumor) and associated graph of subcutaneous tumor growth in C57BL/6 mice. The combination therapy significantly inhibited tumor growth and induced tumor regression in 80% (4 out of 5) of mice (analyzed by two-way ANOVA). (**E**) Percent survival of B16-OVA tumor-bearing mice in various treatment groups (analyzed Log-rank test). All four mice with demonstrated tumor regression survived with no sign of recurrence and survived a tumor rechallenge administered at four months. (**F**) Percent survival of EG7-OVA tumor-bearing mice in various treatment groups. All the mice survived from the DoriVac treatment in combination with αPD-L1. (**G, H**) IFNγ ELISpot results on the splenocytes of the survived mice at 6 months showed more CD8 and CD4 T cell spots (as indicated by the purple color) in all the DoriVac applied groups.

## Discussion

Recent advances have illustrated the potential benefits of DNA origami in the context of cancer immunotherapy ^20, 21, 51^. Several studies have demonstrated that the spatial arrangements of ligands on DNA origami resulted in significant differences in activation of their cognate receptors and downstream signaling in cells ^52–54^. In this study, we explored a square block DNA origami architecture for comparison of spacing of CpG ligands in a range from 2.5 nm to 7 nm, as well as co-delivery of antigen, resulting in Th1-polarized immune responses (Fig. 1B, Fig. S33). Overall, DoriVac improved both CD4 and CD8 T cell proliferation, activation, and cytokine secretion, most noticeably with a CpG spacing of 3.5 nm and 18 CpGs per SQB. Previous studies also investigated other CpG engineering strategies, such as 3’-3’ linked CpG immunomer ^45^, CpG side chains of DNA duplex ^46^, DNA dendrimer with looped CpG ^55^, and G-quadruplex-based CpG ^56, 57^. All these studies indicated that higher-order CpG display may be needed for enhanced immune activation. In this work, we report a finely tuned nanospacing of CpG that is narrower than that explored in previous DNA origami studies. We speculate that the mechanistic basis for the preferred 3.5 nm spacing may relate to the required distance between binding sites on a TLR dimer, based on the CpG-TLR9 crystal structure, and also to geometrical constraints of higher-order TLR9 multimerization ^10^. More generally, co-presenting antigen and adjuvant with precisely defined configurations and stoichiometries on DNA origami provides a tool to characterize critical spatial parameters of danger signals and antigens for future vaccine development. Our comprehensive pre-clinical results confirm that the improved activation, cross-presentation, and cytokine secretion induced on DCs by DoriVac with optimal CpG configurations translate into superior T cell priming and polarization (compared to bolus vaccine, liposomal controls, and suboptimal CpG configurations on SQB). We also observed NK cell activation, which offers additional anti-tumor benefits. The effect was more pronounced when the antigen was co-delivered on the same particle as the CpG adjuvant, demonstrating that co-delivery is a critical aspect of this modular platform.

Some previous studies reported that CpG could assist with elimination of large tumors. These cases were either achieved by aggressive application of CpG both peritumorally and contralaterally ^58^ or preceded by surgical excision of the primary tumor ^59^. CpG-functionalized nanoparticle vaccines have also been broadly investigated recent years ^60^. For example, CpG and HPV E7 peptide co-delivered by mannose-modified liposomes decreased tumor growth of established large TC-1 grafted tumors in a mouse model ^61^. As of 2022, several CpG based drugs for cancer are in clinical trials, with none yet having been approved by the FDA ^62–64^. Display of CpG on DNA origami has been studied by several research groups. Schuller et al attached up to 62 CpG sequences on hollow 30-helix DNA origami to stimulate cellular immunity ^5^. CpG displayed on origami, compared to free CpG, induced greater secretion of IL-6 and IL-12 in DC cells. More recently, Comberlato et al. designed spatial patterns of CpG motifs on DNA origami, with 7 nm spacing that significantly activated TLR9 in RAW264.7 macrophages ^14^. This work represents the earliest published report of how varying CpG nanospacing on DNA origami can affect immune cell activation. Two other recent publications explored CpG display on DNA origami without variation in nanospacing, and both reported therapeutic efficacy in mouse models. One study applied CpG on a pH-responsive DNA origami nanodevice together with antigens, leading to a potent anti-tumoral T-cell response ^21^. The other study applied DNA origami to co-deliver the receptor binding domain of SARS-CoV-2 along with CpGs, leading to robust protective immunity against SARS-CoV-2 ^65^.

We achieved notable therapeutic effects in various tumor models by applying a modest dose of antigen and adjuvant (only 5.4 µg of antigen and 2.2 µg of CpG adjuvant) delivered by a DNA origami nanoparticle, which is a much smaller dose than that used in prior murine studies (100 µg antigen and 100 µg of CpG) ^48^. A similar finding was reported in a previous study, where 4 μg of CpG was fabricated on ultrasmall (∼25–50 nm) polymeric nanoparticles and co-delivered with antigen, resulting in substantial protection of mice from syngeneic tumor challenge ^66^. Another recent study used spherical nucleic acid (SNA) nanoparticles that contained 9 nmol (∼60 µg) of CpG in their animal efficacy study ^67^. In contrast, we were able to achieve a protective immune response against tumors using as little as 0.36 nmol (2.2 µg) of CpG in DoriVac. This highlights an advantage of DNA origami-based vaccines in achieving effective immune responses with minimal amounts of adjuvant. The significant therapeutic effects observed in the context of a low dose of adjuvant suggest that the DNA origami vaccine could potentially reduce adjuvant-related toxicity ^17, 34, 35^. We foresee that neoantigens identified for patients could be quickly and conveniently loaded to prefabricated, adjuvant-containing DNA origami to produce customized cancer vaccines that can be readily combined with FDA approved immune checkpoint inhibitors. This technology platform can also be applied to investigate other cellular ligand-receptor interactions where nanoscale ligand spacing is of critical importance to receptor signaling and subsequent immune polarization. Our study provides evidence that this DNA origami vaccine platform may prove useful for other indications outside of the context of cancer, including infectious diseases, because of its capability to generate robust and durable CD4 and CD8 memory T cell responses.

## Materials and Methods

### Fabrication of SQBs

SQBs were designed using square-lattice CaDNAno and assembled using previously published methods for folding of 3D DNA origami ^68^. Construction plans for SQB, scaffold and staple sequences are listed in Figs. S1–2 and Tables S1–3, respectively. Short synthetic DNA staple strands were purchased on a 100 nmole or 10 nmole scale from Integrated DNA Technologies, Inc.. Scaffold p8634 was produced in-house using previously published protocols ^69^, and purified from endotoxins by extraction with 2% Triton-X114 ^70^. To assemble the structures, DNA staple strands were mixed to a final concentration of 100 nM or 500 nM per strand (as higher concentrations are better for large scale folding as needed for *in vivo* applications). The optimized folding conditions used were 5 mM Tris, 1 mM ethylenediaminetetraacetic acid (EDTA; pH 8.0), 12 mM MgCl_2_, 20–100 nM scaffold, 5 times excess core staple strands (in excess compared to the scaffold concentration), 10 times excess handle conjugated staples and 20 times excess CpG-containing staples. Folding was performed in a thermocycler with the following program: denaturing at 80°C for 15 minutes, and then annealing from 50°C to 40°C decreasing at -0.1°C every 10 minutes and 48 seconds for a total folding time of 18 hours. All constructs were purified using PEG-8000 purification (below). The quality of SQBs was analyzed via agarose gel electrophoresis and negative-stain transmission electron microscopy (TEM).

### Fluorescent labeling of SQBs

SQBs were labeled with Cy5 fluorophores. DNA oligonucleotides /5AmMC6/GGGATAAGTTGATTGCAGAGC-3’(anti-handle) were modified with a 5’ amine and covalently coupled to Cy5 fluorophores via N-hydroxysuccinimide (NHS) ester coupling (Lumiprobe). DNA oligonucleotide (1 mM in ddH_2_O) was mixed with 10 times excess of Cy5- NHS (10 mM in DMSO) and supplemented with 10% of NaHCO_3_ (1M, buffer at pH 8.0). The reaction was carried out in the dark overnight at room temperature shaking at 400 rpm. Zeba size-exclusion and desalting columns (7K MWCO; Thermo Scientific, Waltham, MA) were used to remove unreacted dye through centrifugation at 1000 *g* for 2 minutes. The columns were washed with 400 µL of ddH2O three times before use, according to manufacturer’s protocol. The Cy5 conjugated oligonucleotides were added to the SQB folding pool with 5 times excess to ensure complete conjugation. Eight staples linked with complementary handle strands (5’- GCTCTGCAATCAACTTATCCC-3’) were used to capture the Cy5-linked strands.

### CpG-containing staple strands attachment to DNA origami SQB

CpG-containing staple strands were appended on one end of double helices on the flat side of the SQBs. For most of the staples on the flat face of the SQB, 10 thymine residues were added to the end of the staples on the flat face of the SQB (known as a poly-thymine modification) to minimize aggregation of origami. The CpG oligonucleotides with nuclease resistant phosphorothioate (PS) backbone (5’-tccatgacgttcctgacgtt-3’) replaced the poly T at designed positions for nanoscale patterning as described. The CpG strands were added to the 5’ ends of the staples (Table S3), except for where indicated below (Fig. S5). The CpG-containing staples were either ordered from IDT or home-made by a splint guided T4 ligation (Fig. S5A–D). For CpG1, to ensure that the CpG orientation was uniform, 3’-3’ linkage of CpG with some staples were applied by dibenzocyclooctyne (DBCO)-azide (Sigma, US) click chemistry (Fig. S5E–H). When folding the SQBs, the CpG-containing staple strands were 20 times in excess compared to the scaffold concentration to ensure complete conjugation of the CpG-containing strands.

### Ovalbumin (OVA) conjugation and quantification

For OVA conjugation, 24 sites on the extruding site of SQB were replaced with 5’- CGTCCCCTTTTAACCCTAGAA-3’ handle oligo at poly T positions. The complementary anti-handle oligo 5’-TTCTAGGGTTAAAAGGGGACG-3’ was modified with an amine group at the 5’ end. This oligo was conjugated with succinimidyl 4-(N-maleimidomethyl)cyclohexane-1- carboxylate (SMCC linker through NH_2_-NHS ester reaction. The SMCC-linked oligo was purified through Nap column (GE Healthcare Life Sciences). The purified SMCC-linked oligo was conjugated with OVA through cysteine (thiol)-maleimide group coupling. The maleimide group of SMCC-modified anti-handle can be attached to 4 cysteine residues on OVA (Fig. 1D, Fig. S6). SMCC-linked oligo that was not conjugated with OVA was washed away through 30K Amicon filter (Sigma) filtration for up to 7 times. The conjugated OVA-oligo was incubated with the DNA origami SQB at 37°C for 2 hours at 3 times excess. Free OVA-oligo was removed by PEG precipitation, as described below. To quantify the OVA protein conjugation, a standard curve of fluorophore intensity versus the molar concentration of the AF488-OVA (ThermoFisher, O34781) was made. Briefly, we tested the different AF488 reading by Nanodrop after diluting the protein to a range of concentration that is close to the theoretical concentration of conjugated OVA. The linear curve correlates the OVA molar concentration and the fluorophore intensity.

Three individual experiments were executed to get a repeatable equation for calculation. After OVA was fabricated to the SQB and the vaccine was purified, the AF488 intensity was detected by using the equation to calculation the OVA molar concentration. By dividing the concentration of SQB, the number of OVA per SQB is estimated (Fig. S7).

### SQB purification by PEG precipitation

After annealing, SQB monomers or OVA or peptide conjugated CpG-SQBs were purified from excess staples or excess OVA via PEG precipitation. Typically, 1 × TE buffer (5 mM Tris, pH 8.0 and 1 mM EDTA) containing 15% (w/v) PEG-8000 (Fisher Scientific, BP2331), and 510 mM NaCl was added to the origami sample at 1:1 volume in an Eppendorf tube and mixed gently. Note that it is essential to use PEG-8000 containing less than 5% water. The concentration of MgCl_2_ was adjusted by adding high concentration MgCl_2_ stock to the PEG buffer to obtain 10 mM MgCl_2_ concentration in the final mixture. The mixture was centrifuged at 16,000*g* for 25 minutes at room temperature. The supernatant was carefully removed, and the pellet was re-suspended in 1 × TE buffer supplemented with 10 mM MgCl_2_. This purification procedure was either carried out once or twice, depending on the desired purity level. This procedure is also used to concentrate the SQB monomer when high concentration is desired. The final SQB concentration after PEG precipitation was determined by NanoDrop.

### K10-PEG5k coating of SQBs

SQBs with or without OVA was mixed with oligolysine-PEG5k, in short K10-PEG5k (methoxy-poly(ethylene glycol)-block-poly(L-lysine hydrochloride); n=113, x=10) (Alamanda polymers) such that nitrogen in amines: phosphates in DNA ratio was 1:1, according to the published method ^32^. Samples were incubated at room temperature for 1 hour. After coating, the concentration of SQB was calculated based on the volume increase. K10-PEG5k coating does not affect vaccine efficacy (data not shown). To remove K10-PEG5k for agarose gel verification in Fig. 1H, the nuclease was inactivated by 5 mM EGTA and 10% β-mercaptoethanol in 10 mM MgCl_2_ incubating at 37 °C for 1 hour and then the K10-PEG5k shell was removed by incubating the samples with chondroitin sulfate ^31^ at 37 °C for 2 hours.

### DNase I degradation assay

SQBs (1 µg) were incubated with 1.0 U/µL DNase I (NEB) with 10 × DNase I buffer diluted in water (Gibco). Samples were incubated at 37°C for 30 minutes and then analyzed using 15% denaturing polyacrylamide (PAGE) gel to observe the CpG loading efficiency comparing to the calculated theoretical CpG oligos. Denaturing PAGE gel (15%) was homemade using 9 mL urea concentrate, 4.5 mL urea diluent, 1.5 mL urea buffer (all three from Fisher Scientific), 10 µL tetramethylethylenediamine (TEMED) and 150 µL 10 wt% ammonium persulfate (APS) casting into mini-cassette (ThermoFisher Novex). Loading wells were generated by inserting the comb into the cassette.

### Confocal imaging

Localization of OVA and SQB in the BMDCs was imaged using a Zeiss 710 confocal microscope equipped with 405 diode (405 nm), Ar (458, 488, 514 nm), and white light (633 nm) lasers for live cell tracking. For staining of late endosomes, cells were exposed to CellLight™ Late Endosomes-RFP (Life technologies, Catalog no. C10589) one day before. The cells were cultured with different version of vaccines, and the AF488 from OVA and Cy5 conjugated on SQB were detected for their colocalization with late endosome.

### BMDC isolation and stimulation

C57BL/6 mice at the age of 6–8 months were sacrificed, and the femurs and tibias were obtained. The muscle and connective tissues were removed from the femurs and tibias. A 25G syringe needle was used to flush the inner bone and collect the marrow. The solution was pipetted to release marrow; the marrow clot was then filtered through 40 µm cell strainer. The cells were spun down at 300 *g* for 5 minutes. The cells were resuspended at a density of 3 million cell per mL in RPMI1640 medium supplemented with 10% low endotoxin FBS (Gibco), 50 μM β-mercaptoethanol, 20 ng/mL granulocyte-macrophage colony-stimulating factor (GM-CSF; R&D Systems) and penicillin–streptomycin. Note that red blood cells were not removed. Fresh culture medium was added at day 3 and day 5. On day 7, the floating immature bone marrow derived dendritic cells (BMDCs) were collected for stimulation. BMDCs were seeded in 96 well with the density of 0.1 million in 100 or 200 µL medium. 0.01–1 nM of different DNA origami vaccine and controls were applied for BMDC stimulation for 1–2 days, depending on the experiment set-up. In the majority of our *in vitro* cell culture studies (Fig. 1-3, Fig. S13-S20), we have applied 1 nM SQB origami. For the control bolus vaccine, we have used an equivalent amount of CpG (usually 18 nM CpG if not specified otherwise) and antigens (4-10 nM of antigens). The supernatant was collected and stored in -80°C freezer for later ELISA analysis. The cells were collected for antibody staining and flow cytometry analysis or for downstream T cell co-culture. For cell culture study, there are usually 5 replicates in each group. For each n=5 *in vitro* cell-culture study, a single preparation of cells was harvested from several animals, combined, and then split into 5 replicate wells. Furthermore, for each of these studies, we have repeated 3–4 times with 3–5 replicates each.

### Isolation of OT-I and OT-II T cells and co-culture with BMDCs

The transgenic T cell receptors of OT-I and OT-II mouse were designed to recognize OVA residues 257–264 in the context of H2-Kb (MHC I) and residues 323–339 in the context of I-Ab (MHC II). 6–8 week C57BL/6-Tg(TcraTcrb)1100Mjb/J OT-I mice and B6.Cg-Tg(TcraTcrb)425Cbn/J OT-II mice were purchased from Jackson Laboratory. Mice were sacrificed according to established procedures and the spleens were obtained. The spleens were processed into single cell suspension by mashing the spleen using the top of the plunger of a syringe in a 70 µm cell strainer. CD8 OT-I cells and CD4 OT-II were isolated by MojoSort kit purchased from BioLegend (#480035, #480033) according to the manufacturer protocol. Approximately 2–5 million cells were isolated from each spleen. The isolated cells were labeled with carboxyfluorescein succinimidyl ester (CFSE; Biolegend #423801), incubated in 37°C for 20 minutes. Then the CFSE was bleached by the T cell culture medium, RPMI1640 supplemented with 10% FBS and penicillin–streptomycin. For co-culture, the culture medium for the BMDCs was removed completely. OT-I and OT-II T were added into the wells in fresh T cell culture medium at the number of 0.3-0.5 million based on the yield of isolation (BMDC versus T cells, 1:3-5), and allowed to culture together for 2 days. After 2 days of co-culture, the supernatant was collected and stored in -80°C freezer for ELISA and the cells were collected for antibody staining and flow cytometry analysis. Some T cells were applied for a tumor cell killing assay.

### Tumor cells and *in vitro* tumor cell killing assay

B16-OVA melanoma cells are a kind gift from Dr. David Mooney (Harvard University). B16- OVA cells were cultured at 37 °C in DMEM supplemented with 10% heat-inactivated FBS (Thermo Fisher Scientific), 100 U/mL penicillin and streptomycin (Mediatech), 2 mM L-glutamine (Thermo Fisher Scientific). For *in vitro* tumor cell killing experiment, 3000 tumor cells in 100 µL medium were seeded in 96 well plate one day before CD8 T cell co-culture. The next day, 30000 of the previously activated OT-I CD8 T cell suspension from the BMDC co-culture was added into the tumor cell wells and supplemented with 70 µL additional culture medium. The ratio of tumor cell and OT-I CD8 T cells was 1:10 and cells were cultured together for up to three days, depending on the experimental protocol. The live tumor cells were quantified via confocal microscopy on day 2 or day 3.

### Vaccine distribution

The *in vivo* distribution of DoriVac was assessed in healthy C57BL/6 mice. Mice were subcutaneously injected with DoriVac (200 μg/kg, one injection) at the left shoulder. After a 2, 4, 24 and 48 hour post-injection period, respectively, mice were sacrificed, followed by extraction of major organs (LNs, kidney, liver, spleen, and lung). Fluorescence intensities for Alexa Fluor488 (AF488) conjugated on OVA and Cy5 conjugated on SQB from organs were analyzed using an IVIS Lumina Series III system (PerkinElmer) and quantified using Living Image software (PerkinElmer).

### Flow cytometry

BMDC or T cell suspensions collected as described above were washed with PBS and stained with Zombie UV viability dye (BioLegend, #423108) and then washed with cell staining buffer (BioLegend, #420201). The cells were then stained with cell surface antibodies conjugated with various fluorophores (Table S4). Intracellular staining was initiated by fixation and/or permeabilization reagents (BioLegend, #424401). Cells were then stained with antibodies conjugated with various fluorophores. Antibodies (Table S4) were arranged into different panels, appropriate compensations were set up to compensate for fluorescent emission overlap, and the stained cells were analyzed on Fortessa X-20 (BD Biosciences). Storage events were gated on the population of interest. Flow data was analyzed using FlowJo V10. We followed a generic way to gate on all the cells excluding debris, then singlets and live cells. The gating for single maker or double-marker positive population was out of the live cell population or other mother populations.

### ELISA

Cytokines IL-12, IL-10, IL-6, TNFα, IL-4, TGFβ, IL-2, and IFNγ were evaluated in the culture supernatant, in the serum or the LN tissue by Quantikine ELISA kit (R&D Systems). Following the manufacturer’s instructions, the samples were diluted according to protein concentration, reacted with the appropriate antibodies and the intensity determined via a microplate reader.

### Luminex

The samples for Luminex experiments are either from human cell culture supernatant or from blood plasma. Blood samples collected from mice were collected into heparin-lined tubes and put on ice until processed. Blood was then centrifuged at 4°C, 1500rpm for 5 minutes. The top plasma layer was then collected and transferred to tubes and stored at -80°C until ready to be used. Cytokines were detected using either Bio-Plex Pro™ Mouse Cytokine Th1/Th2 Assay (Bio-Rad, M6000003J7) or the Mouse Magnetic Luminex Assay (R&D Systems, LXSAMSM20) by following methods provided by manufacturer. Human cytokines were detected using the Human Th1/Th2 11-plex Luminex Performance Assay (R&D Systems, LKTM008) by following the manufacterer’s methods.

### RNA sequencing and data analysis

RNA sequencing (RNA-seq) was carried out by Genewiz using an ultra-low input RNA-seq package that includes polyA selection and sequencing on an Illumina HiSeq with 150-bp pair-ended reads. Sequence reads were trimmed to remove possible adapter sequences and nucleotides with poor quality using Trimmomatic v.0.36. The trimmed reads were mapped to the Mouse GRCm38 reference genome using the STAR aligner v.2.5.2b. Unique gene hit counts were calculated by using feature Counts from the Subread package v.1.5.2 followed by differential expression analysis using DESeq2. Gene ontology analysis was performed using Database for Annotation, Visualization, and Integrated Discovery (DAVID). Principal component analysis (PCA) analysis and transcriptomic heat maps were graphed using R4.0.2.

### Animal model and treatment

C57BL/6J female mice (5–6 weeks) were purchased from the Jackson Laboratory and maintained in the Harvard Medical School animal facility. For therapeutic vaccination, after one-week accommodation in the animal facility, 5 × 10^5^ B16-OVA, 2 × 10^5^ B16F10 or 1 × 10^6^ EG7- OVA cells were administered subcutaneously (s.c.) on the right flank of the mice. The establishment of the tumor model was identified by visualization of tumor mass after the tumor cell inoculation. All tumor-bearing mice were randomly divided into multiple groups (n=5-8 in each group) and received appropriate vaccine treatment on day 3, 7 and 14 (or otherwise stated). Tumor growth and mouse survival were recorded. For CD8 depletion, mice were intraperitoneally (i.p.) given 100 μg of anti-CD8 antibody every other day for three total injections, starting the day before vaccination. For sampling, the last treatment was moved to day 11 and the mice were sacrificed on day 15 to make sure the sampling could be completed before the mice in control group started to die. For prophylactic vaccination, mice were treated on day 0 and 7 and then inoculated with 2 × 10^5^ B16-OVA tumor cells on day 14. For anti-PD-L1 or anti-PD-1 combination study, the mice were given 5 doses subcutaneously close to vaccine site at 200 μg per dose diluted in 100 µL PBS on day 6, 8, 10, 12, and 14 in addition to vaccine administration as outlined in the vaccine schedule in Figure 4. All animal studies were approved by the Institutional Animal Care and Use Committee (IACUC) of Harvard Medical School.

### Tumor growth and mouse survival

Tumor growth was evaluated by calculating the tumor volume using caliper measurements of length, width, and height of the tumor every 3–4 days after tumor inoculation. Survival time was calculated as life span from the day of tumor inoculation (day 0). The mice were euthanized in the cases of large tumor size (more than 20 mm in any single dimension) and/or poor body condition were observed. The life spans of the mice were recorded at the day of euthanasia.

### IFN**γ** ELIspot

After splenocytes or PBMDCs were harvested and processed according to protocols above, the cells were plated into a 96-well round-bottom plate, with each well containing cells from an individual mouse in 200 μL of media. We stimulated each well with 2 μg/mL of the associated peptide. The MHC I OVA peptide epitope (OVA peptide (257-264), sequence: SIINFEKLC, ThermoFisher) was used to probe antigen-specific CD8+ T cell responses, while the MHC II OVA peptide epitope (OVA peptide (323-339), sequence: ISQAVHAAHAEINEAGR, Thermo Fisher) was used to probe antigen-specific CD4^+^ T cell responses. After 48 hours of stimulated pre-incubation at 37°C, we collected the cells (1200 rmp, 5 minutes, 4°C) and resuspended in 100 μL of sterile splenocyte media. We incubated the cells for 36 hours at 37°C and then process the ELIspot plate according to the manufacturer’s instructions (RND Systems, Mouse IFNγ ELIspot kit, #505841). Any wells that were dry after the incubation steps were removed from the analysis. The plate was analyzed via an ELIspot plate reader by Dana Farber Cancer Institute’s Center for Immuno-oncology (CIO) Translational Immunogenics Laboratory (TIGL). The number of spot forming units (SFUs) per initial number of cells plated was quantified.

### Statistical analyses

GraphPad Prism 9 was used to make graphs, analyze the statistics, and calculate P values in most cases. One-way ANOVA was applied for flow cytometry data and ELISA data with Tukey’s post hoc multiple comparisons test. The analysis of mouse tumor volume trajectory was performed using a generalized linear mixed effect model, fit to a log transformed response looking at tumor volume growth rate across days. This analysis was executed using R version 4.04 and utilizing the lme4 package version 1.1-26 for all mixed modeling estimation. Survival curves were analyzed by Kaplan-Meier method and log-rank test. A two-tailed student t-test was applied to analyze the significance between two groups. P value ≤ 0.05 was considered statistically significant. ‘*’ refers to P ≤ 0.05; ‘**’ refers to P ≤ 0.01; ‘***’ refers to P ≤ 0.001; ‘****’ refers to P ≤ 0.0001. Error bars represent standard error of the mean (SEM).

## Acknowledgments

We thank Qi Yan, Zhao Zhao, Tian Zhang, Pascal Lill, Pranav Prabhala, Leo Chou, Jaeseung Han, Dionis Minev, Benjamin Everhart, Jie Deng, David Zhang, Kwasi Adu-Berchie, Hawa Dembele, Anjali Rajwar, and Kow Simpson for aiding in labor support, experimental design, exploring experiments, or manuscript proof-reading. We also thank Maurice Perez, Michael Carr, and Eric Zigon for their assistance in the lab management and facility usage. We thank Maartje Bastings and Aileen Li for exploration, before the current project initiated, of CpG-functionalized DNA-origami barrels for immune stimulation.

## Author contributions

Y.C.Z. developed and planned the experiments, carried out the vaccine fabrication and validation, and wrote the manuscript. J.H.R. and W.M.S provided experimental and theoretical guidance and edited the manuscript. O.J.Y. assisted Y.C.Z. in performing experiments, analyzing data and manuscript editing. C.M.W., F.M.A., J.I.M. and GI assisted with the DNA origami design, modeling and fabrication. H.B. performed the RNA sequencing analyzing. M.O.D, M.S, M.S, A.R.G., A.V. and M.S. helped with animal study design, modeling and sampling. T.C.F assisted with confocal experiment. Y.C. assisted with 3D modeling and manuscript editing. G.G.F and D.N. guided statistical analysis. C.J.W. and D.B.K. guided the experiment design and offered manuscript editing. D.M. and I.C.K. provided project support and manuscript editing.

## Data and materials availability

Data supporting the findings of this study are presented in the paper and the supplementary materials.

## Supplementary Materials

Figures S1-S33

Tables S1-S4

Supplemental methods

