## Supplementary Information for "Fine tuning of CpG spatial distribution with DNA origami for improved therapeutic cancer vaccination"

**Authors:** Yang C. Zeng^1,3,4^, Olivia J. Young^1,3,4,7^, Chris M. Wintersinger^1,4^, Frances M. Anastassacos^3,4^, James I. MacDonald^3^, Giorgia Isinelli^3,8^, Maxence O. Dellacherie^5^, Miguel Sobral^5^, Haiqing Bai^3^, Amanda R. Graveline^3^, Andyna Vernet^3^, Melinda Sanchez^3^, Kathleen Mulligan^1^, Youngjin Choi^2^, Thomas C. Ferrante^3^, Derin B. Keskin^6^, Geoffrey G. Fell^9^, Donna Neuberg^9^, Catherine J. Wu^6^, David J. Mooney^3,5^, Ick Chan Kwon^1,2,10^, Ju Hee Ryu^1,2^*, William M. Shih^1,3,4^*

**This PDF file includes:**

Figs. S1 to S33

Tables S1 to S5

Supplemental methods

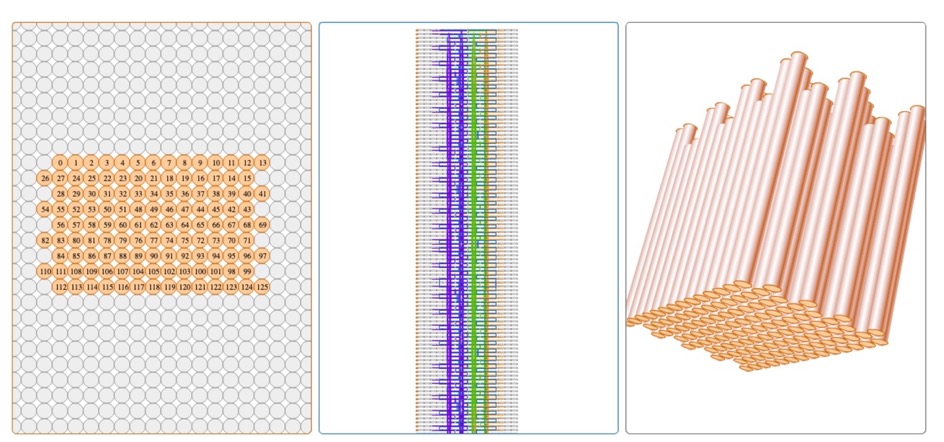

**Fig. S1. CaDNAnoSQ SQB design.** This is a screen shot of the basic SQB design. The structure was designed using CaDNAnoSQ and Python software. The scaffold is p8634, whose corresponding DNA sequence is presented at Table S1.

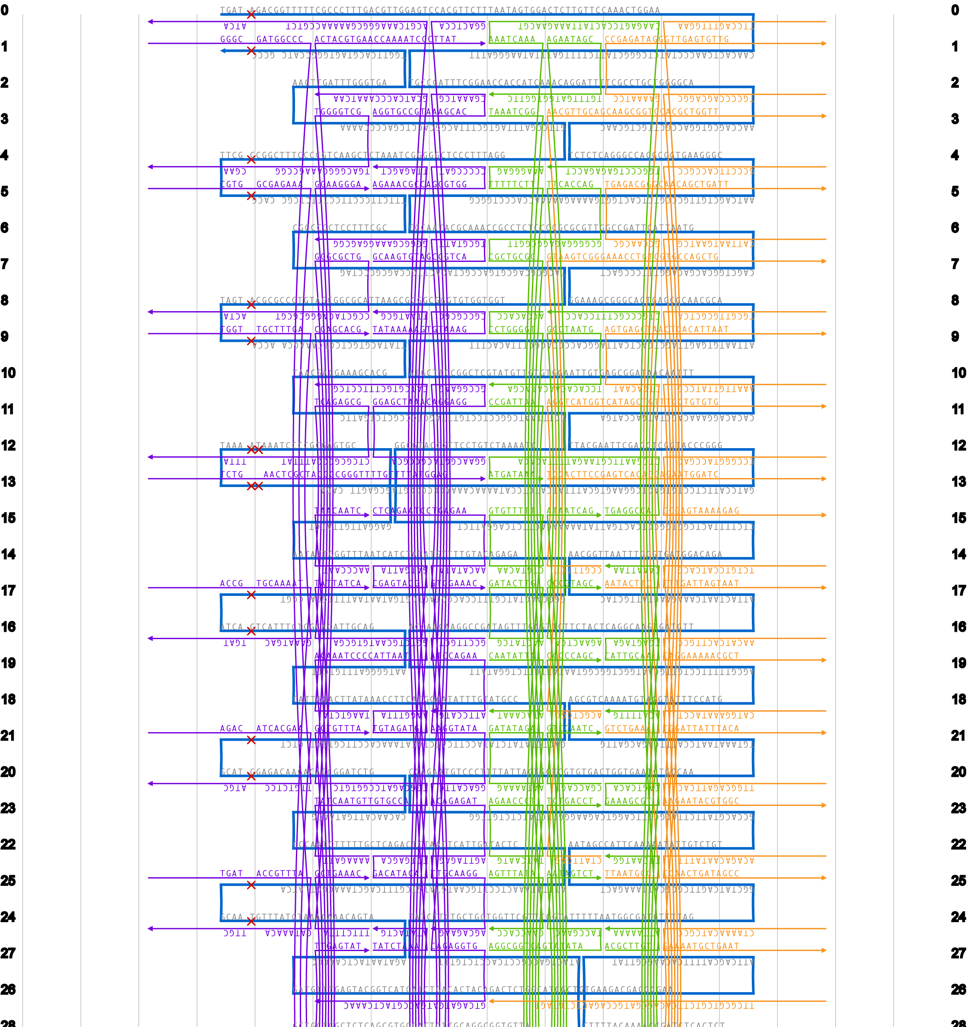

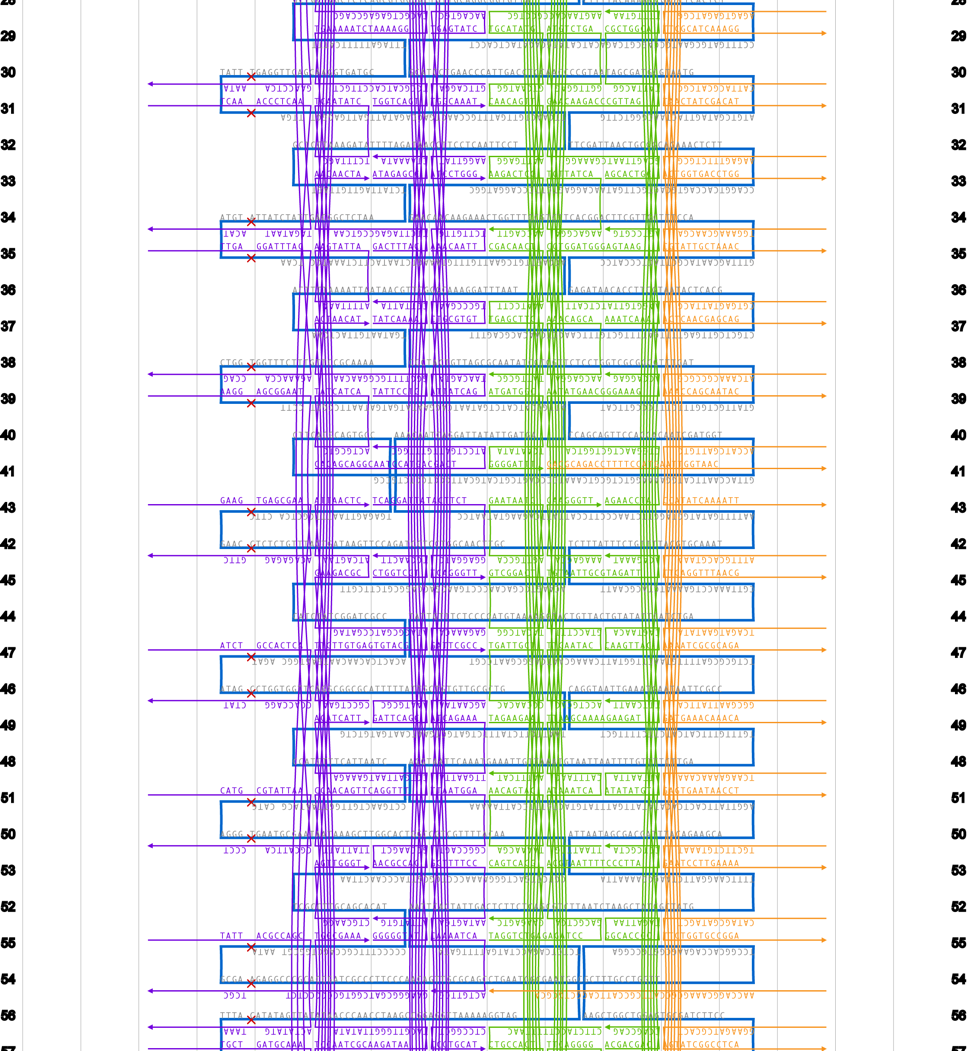

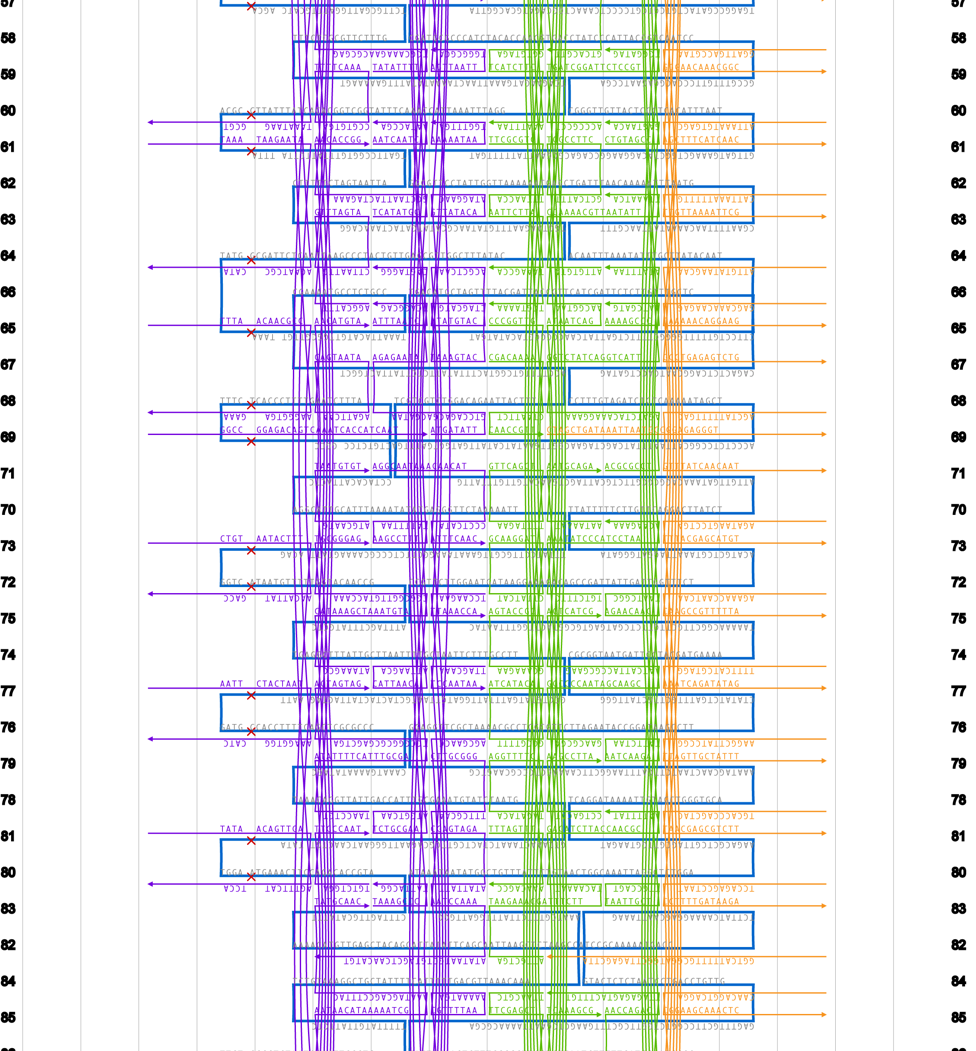

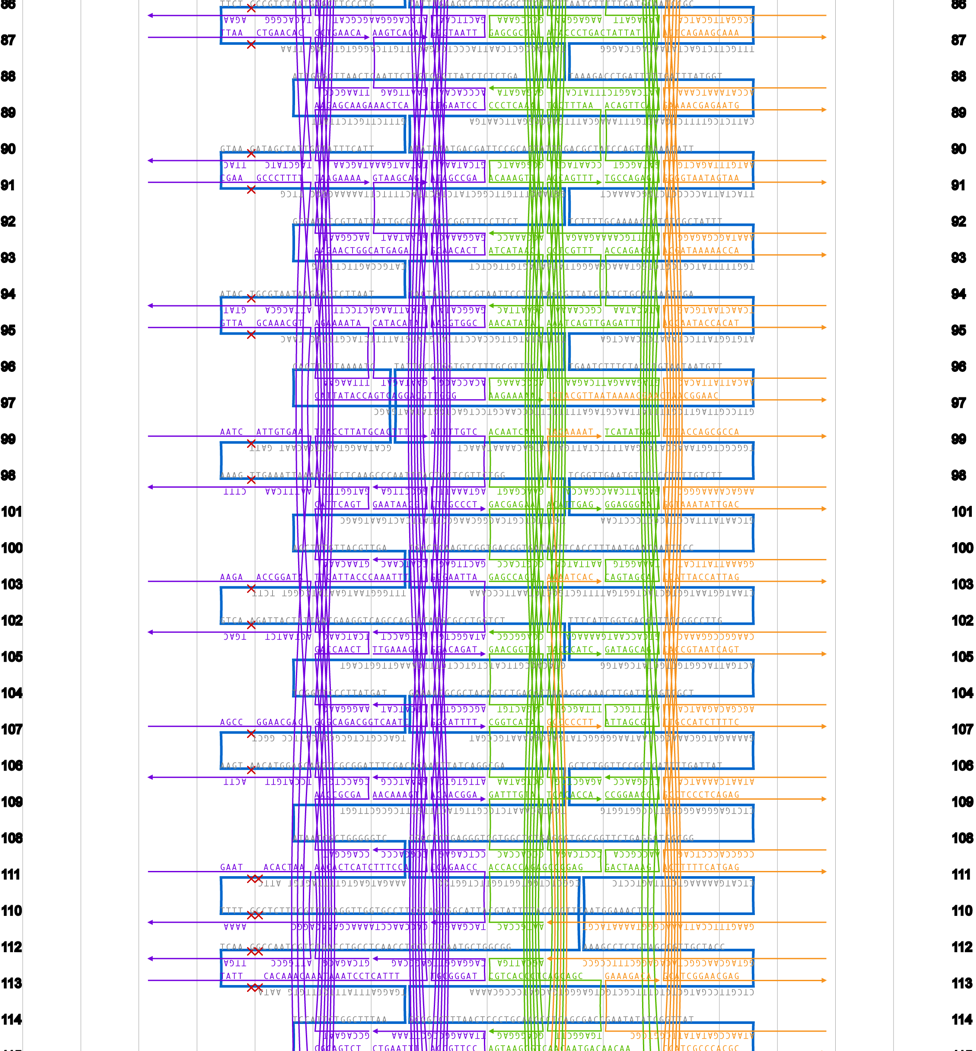

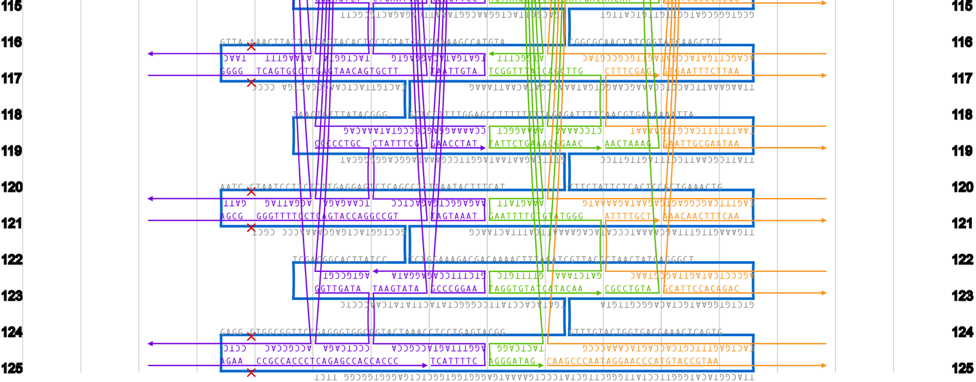

**Fig. S2. Routing strategy of SQB as displayed on the CaDNAnoSQ interface.** For more detail, refer to the .json file.

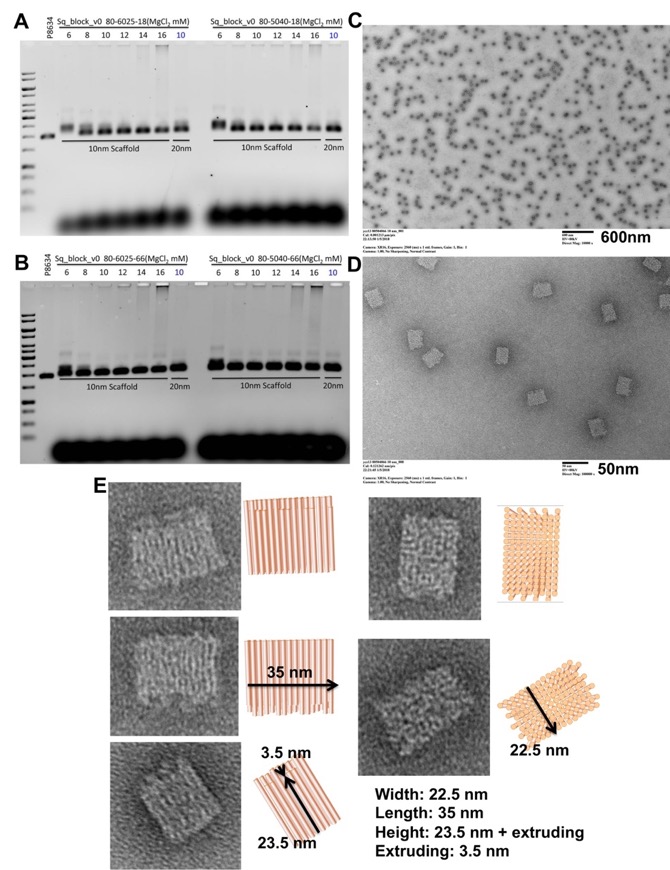

**Fig. S3. SQB folding optimization.** (**A, B**) The SQBs were annealed under various temperature ramps, time courses and MgCl_2_ concentrations. Time course for **(A)** 18 hours, **(B)** 66 hours and temperature condition is left panel (temperature ramp from 60 to 25°C), right panel (50 to 40°C). MgCl_2_ concentration are denoted at the top of agarose gel images. The optimal condition was determined as the condition which produced a high folding efficiency and minimal aggregation of the resulting SQB. The optimal temperature ramp conditions were determined to be: 80°C for 15 minutes, and then a declining temperature ramp starting at 50°C and declining to 40°C with -0.1°C every 10 minutes and 48 seconds. The optimal time course was 18 hours and the optimal MgCl_2_ concentration was 12 mM. (**C, D**) TEM images of the optimally folded SQBs. (**C**) There is no obvious SQB aggregation. (**D**) A higher magnification image showing the SQB structure. (**E**) The high magnification images of a single SQB and the precise dimensions.

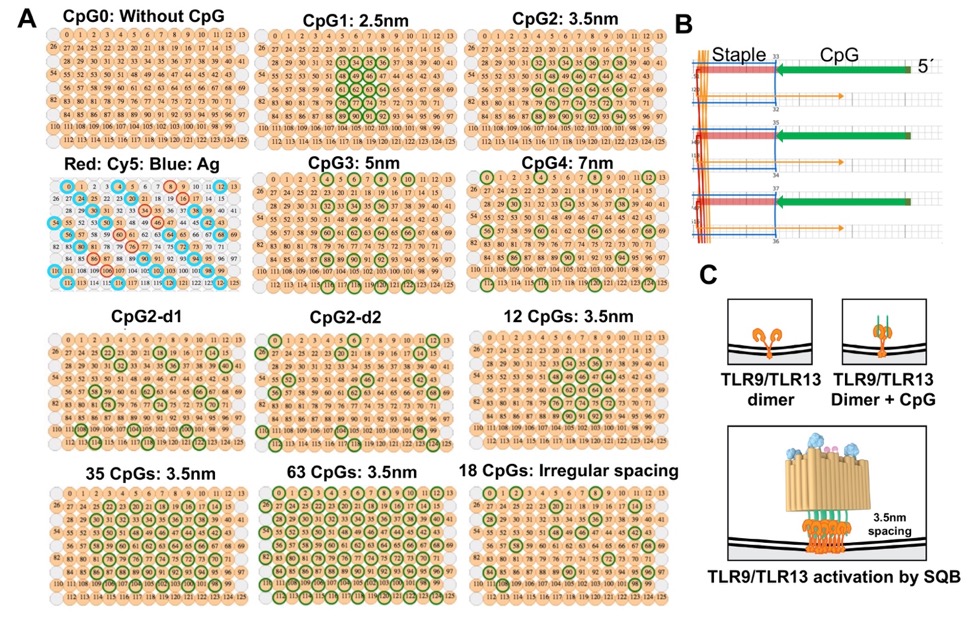

**Fig. S4. CpG spatial distribution on the flat face of the SQB.** (**A**) These figures show the Cy5 dye, antigen (Ag) and CpG positions on DNA origami. CpG is conjugated on the end of the double helices on the flat face of the SQB and Cy5 and antigen are conjugated on the opposite site of CpG face which has extruding helices. The CpG strands was fabricated as an extension of the staple strands at desired positions. The 126 helices are labeled between 0 and 125. The sequences of the CpG staples and positions are found in Table S3. DNA origami without CpG conjugation is denoted as CpG0. DNA origami labeled as CpG1, 2, 3, 4 has eighteen CpG strands with various spacing distance 2.5, 3.5, 5, 7 nm, respectively. CpG2-d1 and CpG2-d2 has wider spacing for CpG dimers (dimer CpG spaced at 3.5 nm). Finally, DNA origamis with different number of CpG strands (12, 18, 35, 63) are denoted as 12 CpGs, 18 CpGs (same as CpG2), 35 CpGs and 63 CpGs, respectively. 18 CpGs with random position of CpG strands is denoted as 18 CpGs with irregular spacing (CpGi). (**B**) A schematic figure from CaDNAno file showing CpG attachment from CpG2 on the end of double helices of No. 32, 34 and 36. All the CpGs fabricated are single stranded, the CpGs are added during phosphoramidite synthesis of the staple strands by the vendor (IDT), or else added via splint-templated ligation. (**C**) Proposed model of CpG and TLR9/TLR13 interaction that induces Th1 polarization when CpG is spaced at 3.5 nm.

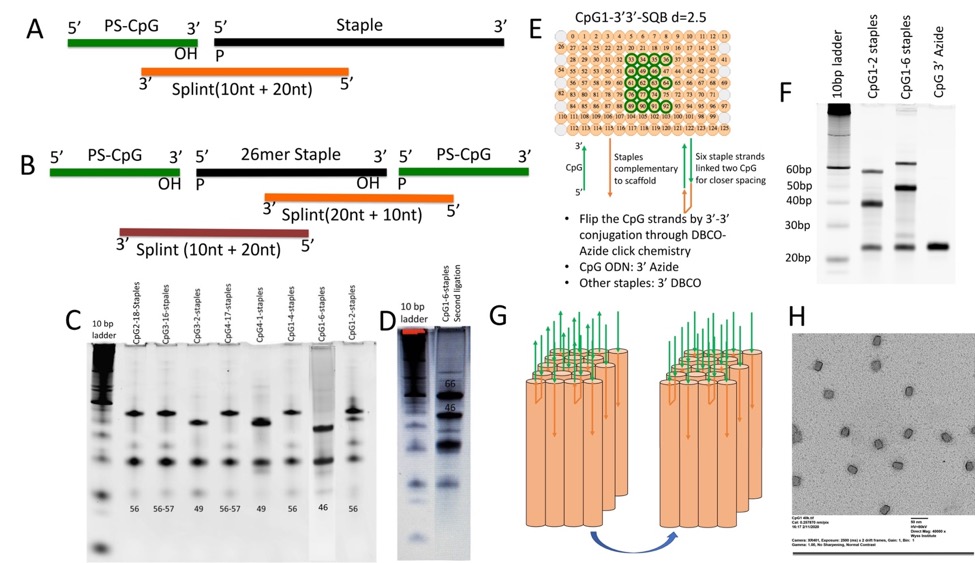

**Fig. S5. CpG ligation.** Most experiments in this study relied on commercially produced CpG-containing strands. We also self-assembled the CpG strands by ligating the CpG with original staples, and purifying the ligated staples via denatured PAGE gel. This technique is a viable option to minimize the costs associated with CpG-SQB fabrication. (**A**) Ligation strategy for all CpG patterns except for CpG1. The CpG has a 5´ end extruding from the origami. (**B**) Ligation strategy for CpG1 (2.5nm) where some of the staples linked CpG on both 5´ and 3´ end. (**C**) PAGE gel demonstrating the ligation results. The top band is the ligated full-length product. The second faint band is the unreacted original staples without poly Ts. The third bright band is all the splint DNA after denaturation. The faint forth band is the excess of CpG strand. (**D**) PAGE gel corresponding to the alternative ligation strategy for CpG1 with a CpG linked to both the 5´ and 3´ staple ends. (**E**) The ligation method described in (B) cannot generate uniform orientation of CpG and obtain the dense CpG patterned associated with CpG1. To this end, we applied DBCO-azide click chemistry to ligate 3´ and 3´ ends of staples and CpG strands to ensure all CpG strands have the same orientation, with the 5´ ends extruding from the SQB. The associated staples were modified with DBCO on their 3´ ends, and CpGs were modified with Azide on their 3´ ends. We also investigated a variant CpG2L (i.e. 3.5 nm spacing) with the 3´-3´ DBCO-azide linkage for all eighteen CpG extensions, and found the ability of these nanoparticles to induce Th1 polarization was diminished compared to the CpG2. Therefore we cannot conclude that the effects observed on dendritic cells between CpG1 and CpG2 are due solely to differences in spacing. (**F**) Denaturing PAGE gel result showing the successful conjugated strands (upper band). (**G**) A schematic demonstrating the alternative fabrication strategy that results in uniform CpG orientation for CpG1. Green arrows indicate CpG strands, head is the 3´ end and tail is the 5´ end. After ligation strategy with click chemistry, CpG strands have uniform orientation. (**H**) TEM imaging of CpG1-SQB, demonstrating monodispersed SQBs.

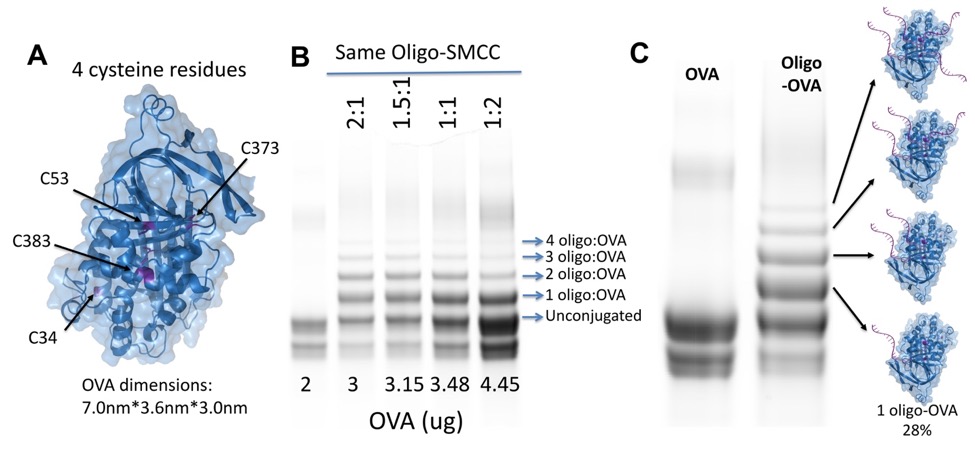

**Fig. S6. Ovalbumin (OVA) conjugation optimization.** There are 4 cysteine residues on the whole protein ovalbumin which could be conjugated to DNA oligonucleotides via an SMCC linkage reaction. Our aim was to generate a majority of the OVA product with a single oligonucleotide attached (1 oligo-OVA) for SQB appending. (**A**) The 3D structure of OVA protein. Cysteine residues, which can be linked with SMCC and subsequently conjugated to a DNA oligonucleotide, are labeled. (**B**) Oligo-SMCC and OVA were reacted at various ratios. The SDS PAGE gel showed the unconjugated OVA protein or OVA conjugated with 1~4 oligos. (**C**) SDS PAGE gel of the OVA-oligo reaction products. A 1:1 molar ratio of OVA to oligo-SMCC increased the yield of the singly appended 1 oligo-OVA protein compared to more conjugated forms of the OVA and unconjugated OVA. The 1 OVA-oligo product had a yield of around 25–28% and was the dominant SMCC reaction product under the optimized conditions.

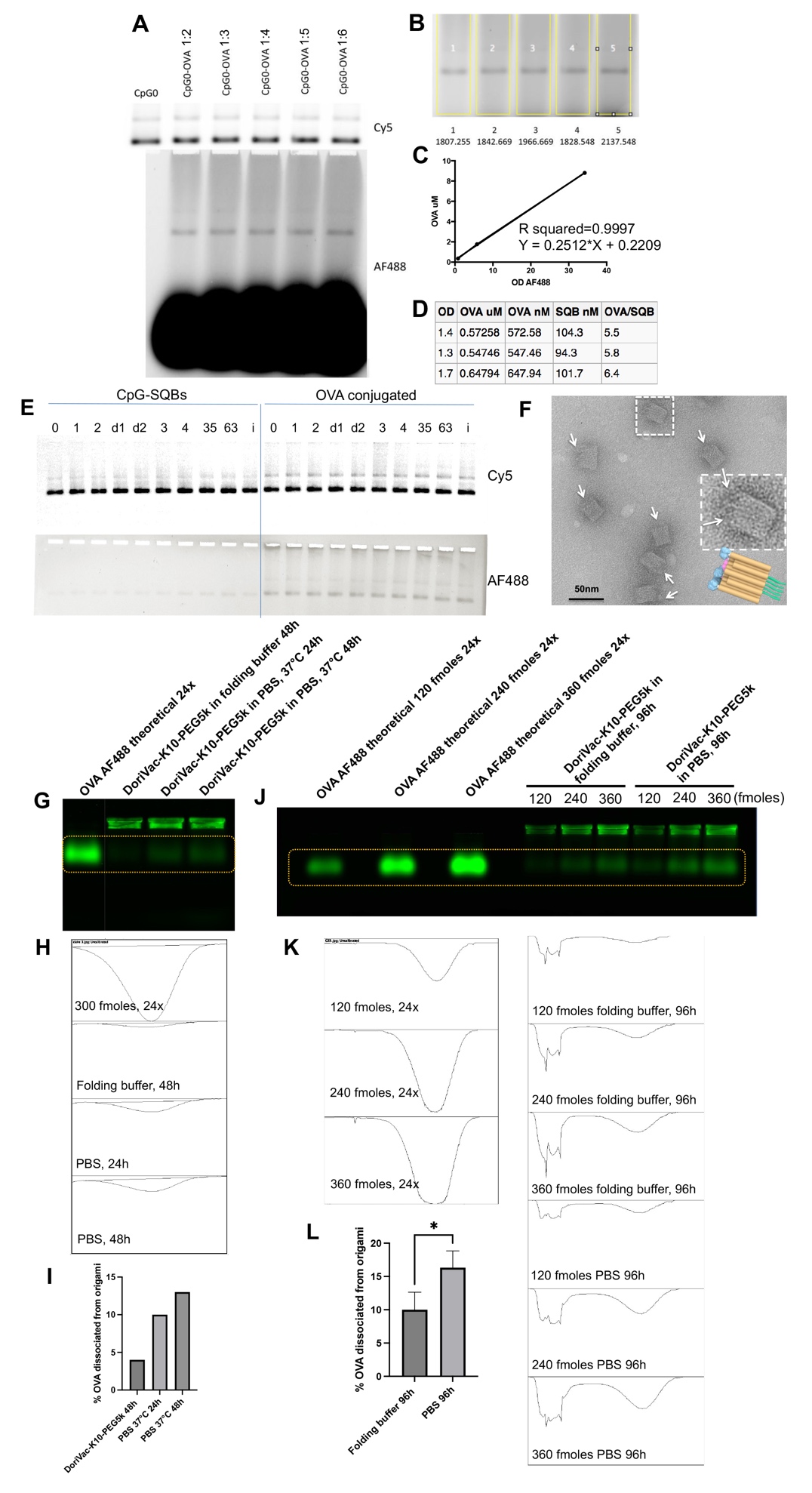

**Fig. S7. OVA conjugation, purification, and concentration verification.** The OVA-oligo conjugate was mixed with purified CpG-SQB directly and incubated for 2 hours at 37°C. (**A, B**) Various concentrations of OVA-oligo were mixed with the SQB to determine the appropriate amount to ensure complete hybridization. The Alexa Fluor 488 (AF488) fluorescent intensity was quantified via ImageJ and the appropriate amount of OVA-oligo for optimal conjugation was determined to be three times in excess of SQB multiplied by five times due to the yield of 1 OVA-oligo. (**C**) The AF488 intensity correlates with the number of conjugated OVA. We quantified the AF488 intensity to determine the average amount of OVA conjugated to the SQB. (**D**) We performed three different conjugation experiments of OVA to SQB and calculated independently the amount of OVA conjugated to the SQB. Due to the size constraints of the OVA protein, the average of these three trials showed that 6 OVA proteins can be appropriately conjugated to the SQB, while we were able increase this number to 12 OVA proteins in later experiment by improving the purity of the OVA-oligo. (**E**) All constructs showed AF488 signal after conjugation with OVA. CpG conjugated without OVA does not show strong AF488 signal (left panel). OVA conjugation correlates with increased AF488 signal (right panel). (d1 and d2 refer to CpG2-d1 and CpG2-d2, 35 and 63 refer to 35 and 63 CpG strands were conjugated onto SQB at spacing of 3.5 nm) (**F**) TEM images of conjugated CpG2-OVA-SQBs as the complete DoriVac vaccine. The arrows delineate the OVA protein. (**G-I**) To assess the handle/anti-handle stability, we conducted a PBS incubation experiment on OVA-conjugated DoriVac. The DoriVac was initially coated with K10-PEG5k and subsequently diluted 50 times (volume dilution) in PBS buffer. We evaluated the OVA conjugation using agarose gel analysis. Over the course of incubating the DoriVac at 37 ℃, we observed a minor release of OVA from the origami structure. To quantify the amount of OVA protein dissociated from the origami, we utilized Image J to measure the fluorescent intensity of AF488 conjugated on OVA (**H**). This intensity measurement of OVA in the agarose gel allowed us to compare it with the theoretical intensity of 24 OVA molecules (**I**). It is important to note that the K10-PEG5k coating neutralizes the charge of DNA origami, enabling them to remain in the well. As a result, the fluorescence detected in the well indicates the presence of OVA that remains conjugated to the DNA origami. (**J-L**) To further investigate, we conducted a comparison between the sample incubated in PBS for 4 days and that incubated in folding buffer. For a more precise analysis of OVA dissociation, we utilized a series of sample dilutions during gel examination. Our findings revealed that OVA dissociation occurs more rapidly in PBS than in the folding buffer through image J fluorescence intensity analysis (**K, L**). These results underscore the importance of enhancing the preservation method for DNA origami after conjugation with cargos, especially for long-term storage, to ensure successful clinical translation.

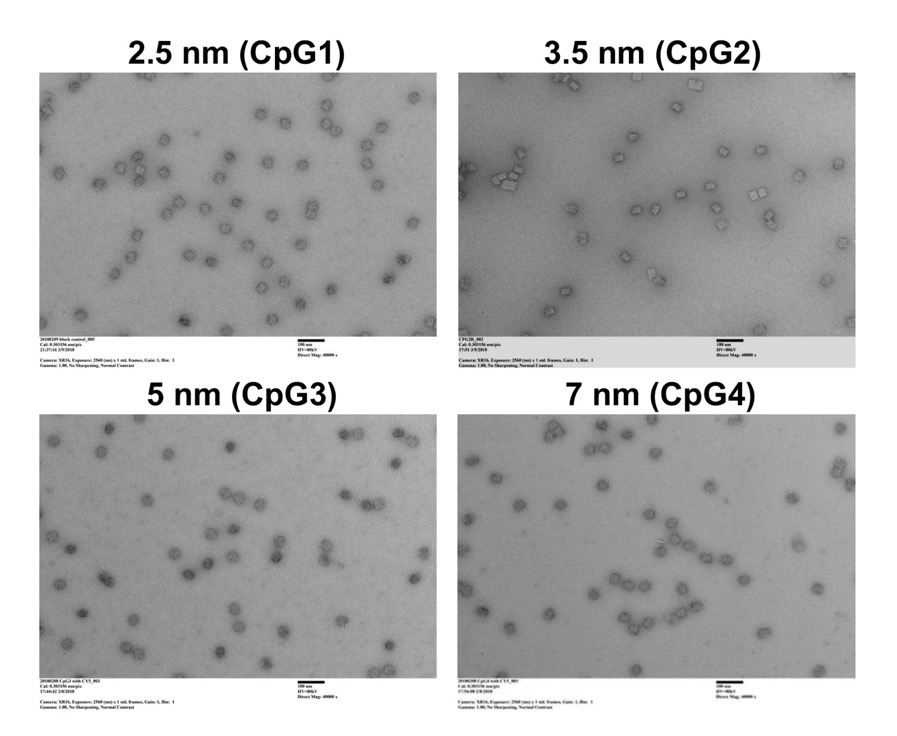

**Fig. S8. Folding of CpG-SQB constructs with varied CpG spacing.** TEM images of CpG1 to CpG4.

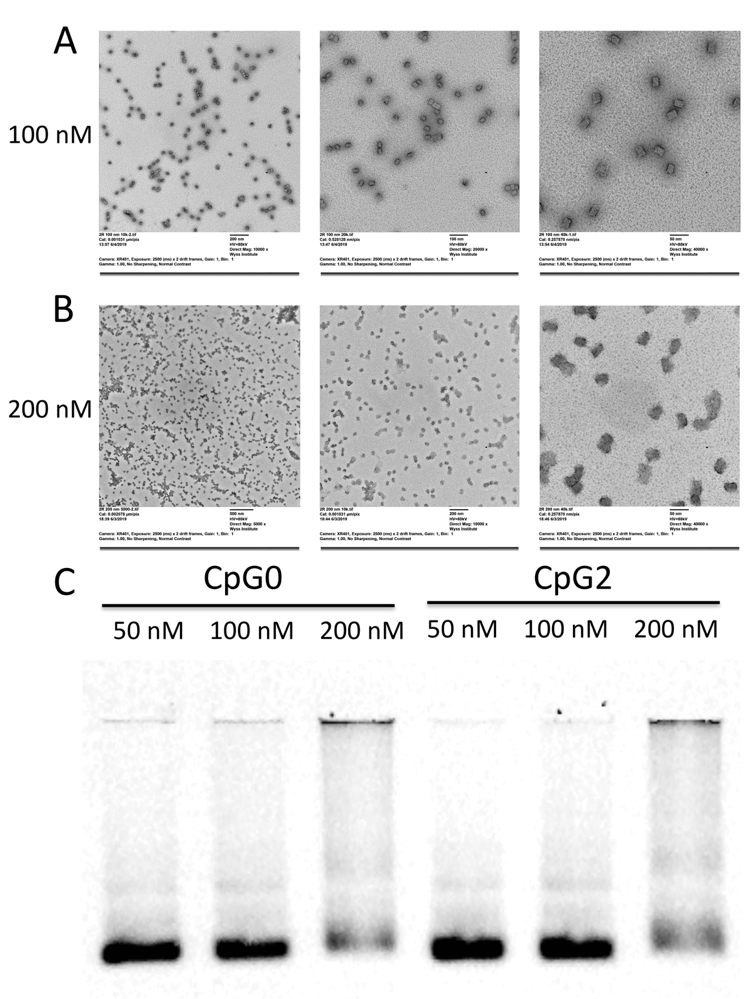

**Fig. S9. Folding of CpG-SQB with higher concentration for *in vivo* application.** We increased the scaffold concentration for CpG-SQB folding to as high as 100 nM scaffold concentration without jeopardizing structure integrity. For optimal folding, The scaffold was cultured with 5 times excess basic staple strands and 10-20 times excess of the functional staple strands for linking cargos. (**A, B**) TEM images of SQBs folded with 100 or 200 nM concentration of scaffold. (**C**) The agarose gel result of SQBs folded at different scaffold concentrations. 100 nM is the highest scaffold concentration that maintains structural integrity. At 200 nM scaffold concentration, the folding quality appears to be lower than for 100 nM.

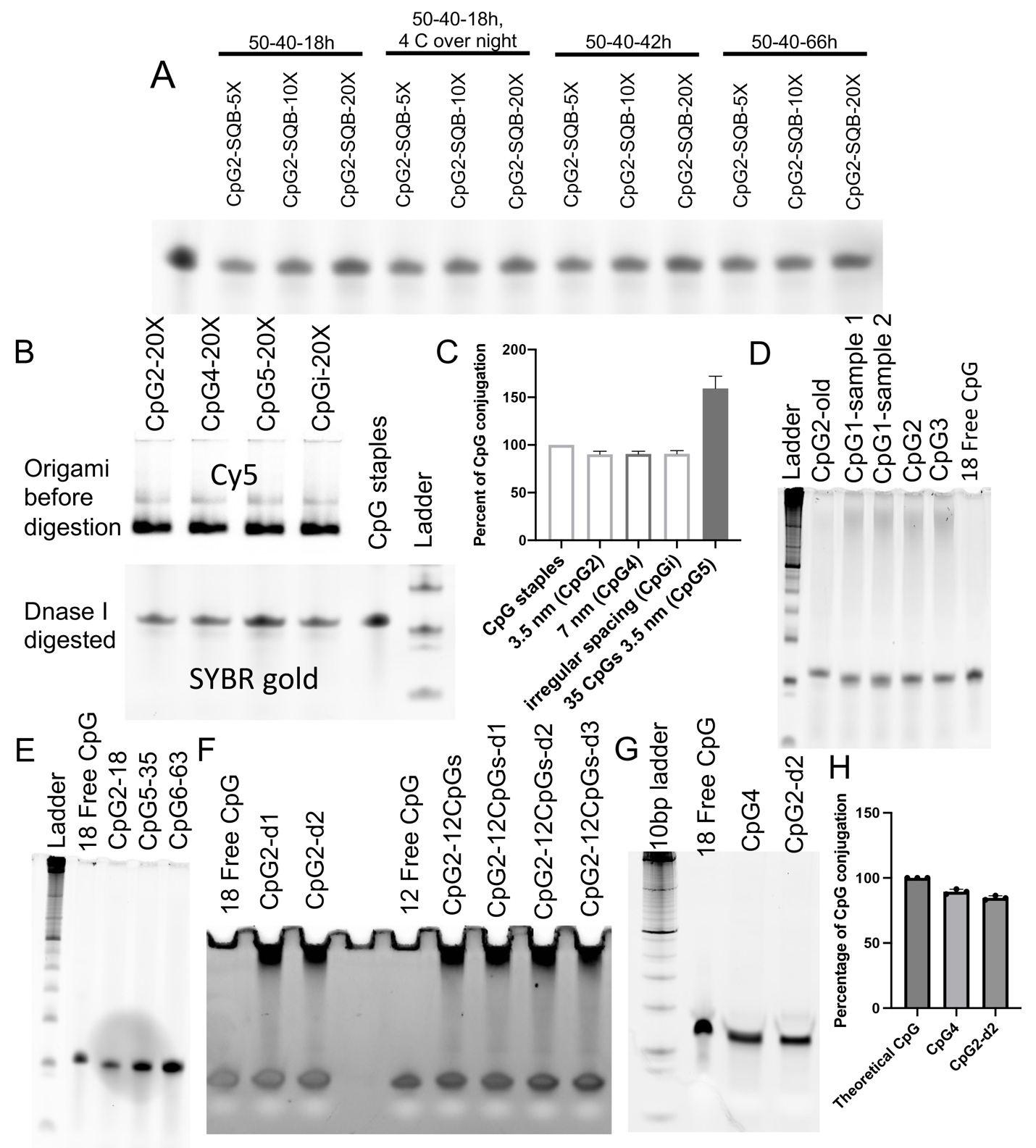

**Fig. S10. CpG loading efficiency.** This experiment verified the experimental number of CpG strands loaded onto the SQB DNA origami. We attempted various amounts of excess CpG-containing staple strands and verified 20 times in excess of the theoretical limit of CpG-containing staple strands could achieve ~90% loading efficiency. (**A**) The SQBs were digested by DNase I, which degrades the SQB, but not the associated CpG. CpG has a PS backbone that is not targeted by DNase. Various temperature ramps, annealing times and different amounts of excess CpG-containing staple strands were tested. After DNase I digestion, the solution was analyzed via denaturing PAGE gel at 250 V for 40 minutes. (**B**) Upper: agarose gel of various CpG-SQB constructs before DNase I digestion, denaturing PAGE gel of various CpG-SQB constructs after DNase I digestion. The four lanes correspond to four different constructs: CpG-SQB folded with various CpG spacings (CpG2: 3.5 nm, 18 CpGs; CpG4: 7 nm, 18 CpGs; CpGi: irregular spacing, 18 CpGs) and various amounts of CpG per SQB (CpG5: 35 CpG, 3.5 nm). The agarose gel band corresponds to the dye on SQB (upper). The corresponding denaturing gel was stained with SYBR gold (lower). (**C**) The band intensity was calculated by ImageJ and compared to the band intensity corresponding to the theoretical amount of conjugated CpG. 100% loading efficiency indicates that all 18 possible CpG sites were conjugated. “150% loading efficiency” in this case indicates that 27 CpG were successfully conjugated for a CpG-SQB design that has 35 possible CpG conjugation sites (indicating approximately 80% actual loading efficiency). (**D-F**) Denaturing PAGE gel results of CpG loading efficiency comparing more samples corresponding to the CpG designs shown in Fig. S4 and Fig. S20. (**G, H**) Denaturing PAGE gel and quantification results showing CpG abundancy of CpG4 and CpG2-d2 after being cultured in the complete culture medium with 10% FBS for 2 hours. The CpG conjugation efficiency remained at a similar level when compared to that measured before incubation with cell media (c.f. Fig. S10 B,F,G). “18 free CpG” sample means intact 18 free CpGs without culturing, indicating theoretical CpG amount.

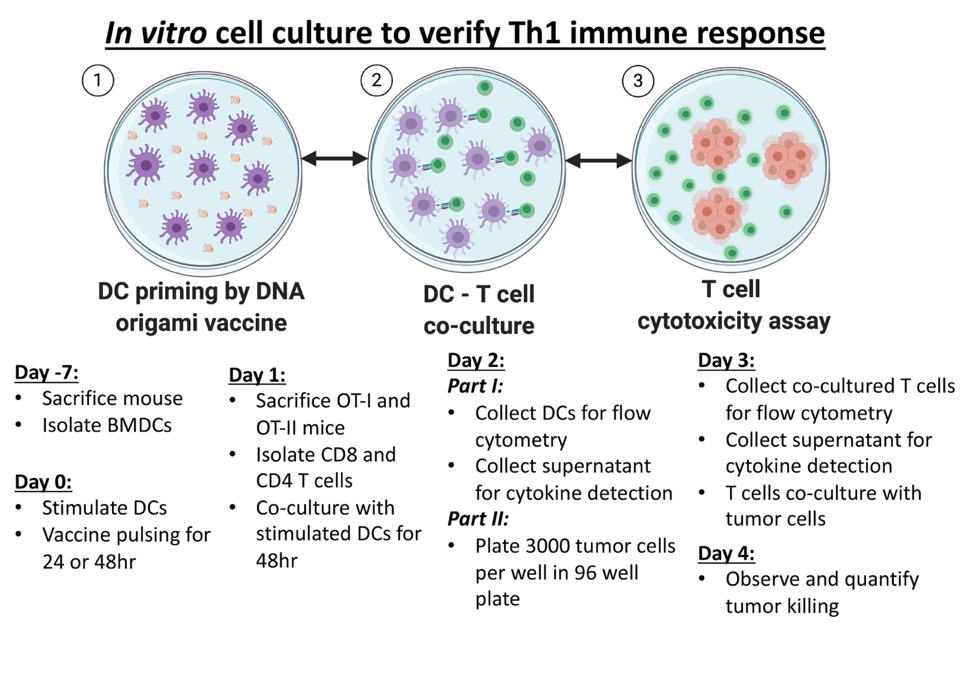

**Fig. S11. Serial cell co-culture study design to verify Th1 immune polarization by SQB stimulation.** The cell culture study took place in sequence. Dendritic cells (DCs) were isolated from mouse bone marrow to obtain immature DCs. BMDCs were stimulated by various DNA origami vaccine constructs. The DC supernatant was removed and the pulsed BMDCs were co-cultured with freshly isolated OT-I CD8 or OT-II CD4 T cells for 48 hours. SQB-stimulated DCs present the OVA antigen to T cells and provide co-stimulatory signals for T cell activation and proliferation. The DCs and T cells and associated supernatant were collected for flow cytometry and ELISA. Amplified and activated T cells were then co-cultured with B16-OVA tumor cells in a 1:10 (T cell: tumor cell) ratio. Tumor cells were labeled with calcein green fluorescent dye. Tumor killing effects were observed by fluorescence release or via live cell counting.

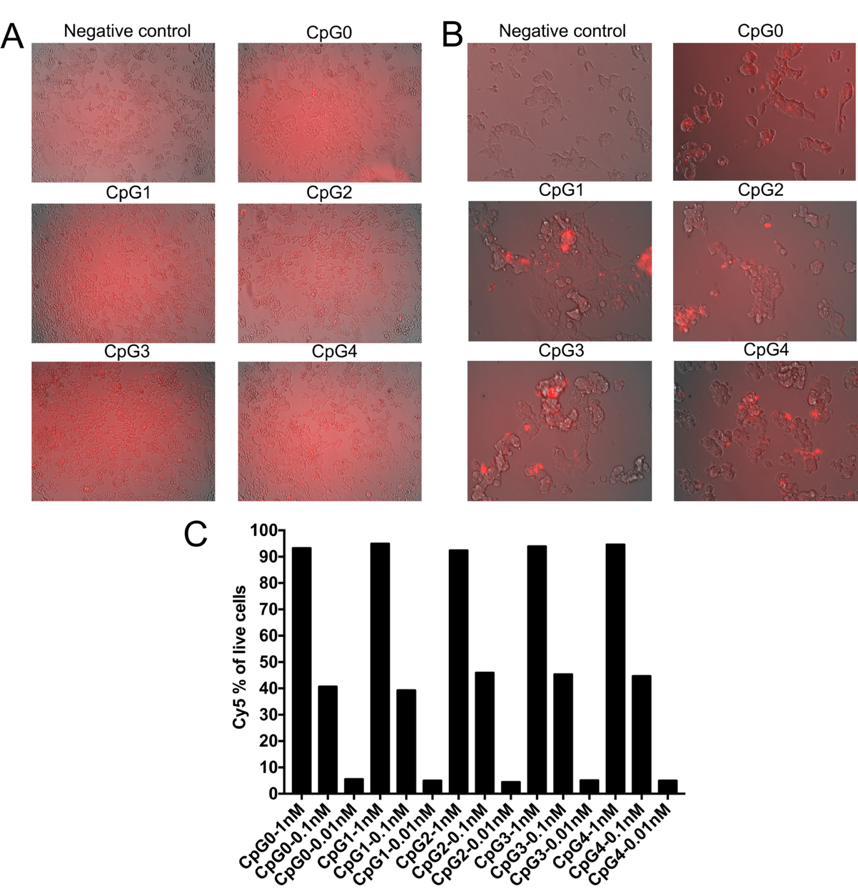

**Fig. S12. CpG-SQB uptake in three cell types.** Our group recently discovered that low-salt denaturation of DNA origami can be prevented, and nuclease digestion greatly slowed, by electrostatic coating of DNA nanostructures with PEGylated oligolysine (K10-PEG5k) (*4*). This finding extends of the utility of DNA origami *in vivo*. 293 T cells and HeLa cells grow robustly after adding K10-PEG5k coated DNA origami structures (CpG0-SQB)**.** 293 T cells have more pseudopods for endocytosis compared to HeLa cells, leading to more efficient uptake. (**A**) Uptake of CpG-SQB in HeLa cells. Images taken by fluorescent microscope (10×). (**B**) Uptake of CpG-SQB in 293T cells. Images taken by fluorescent microscope (20×). (**C**) Percentage of Cy5-expressing BMDCs analyzed by flow cytometry, after culture with various SQB concentrations for 24 hours. 1 nM of SQB DNA origami almost saturated the BMDCs, as demonstrated by the greater than 90% uptake efficiency. Consequently, 1 nM DNA origami were applied for all subsequent cell-culture experiments.

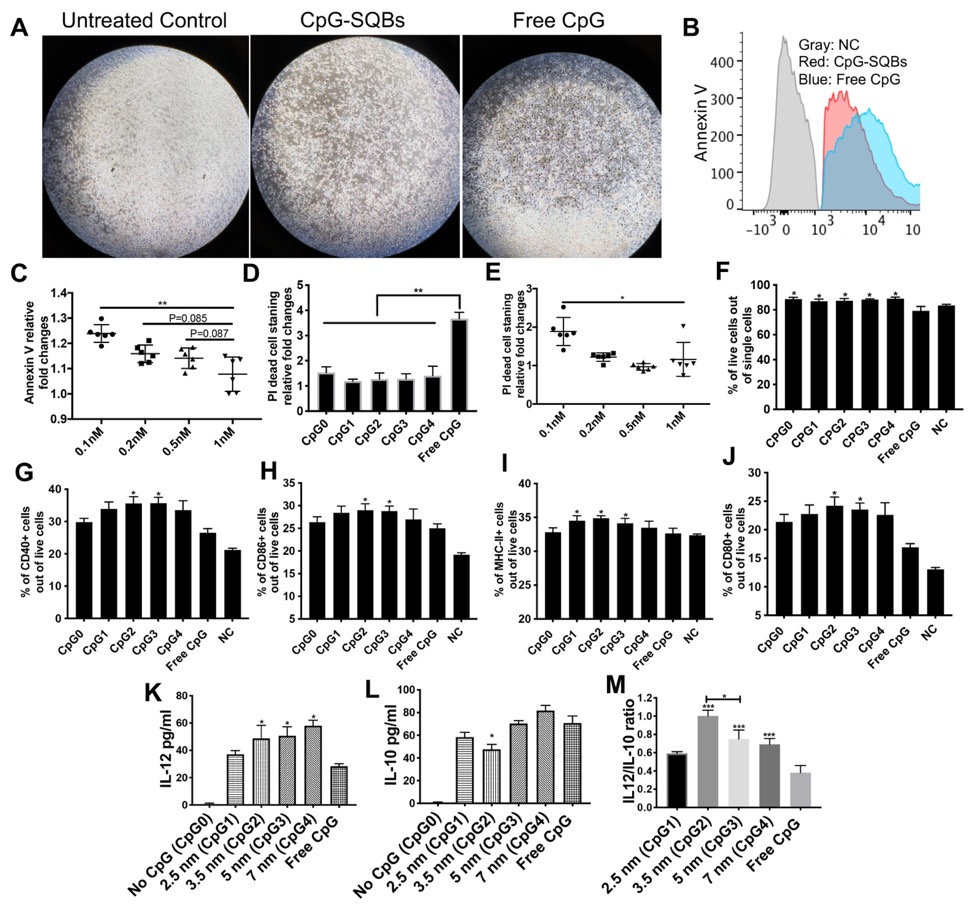

**Fig. S13. BMDC maturation was stimulated by CpG-SQBs.** CpG conjugated SQBs (without conjugated antigen) were applied to stimulate immature bone marrow derived dendritic cells (BMDCs). Various SQB constructs containing CpGs (based on the concentration of 1 nM SQB) or free CpG (18 nM) was treated. (**A**) Representative microscope images showing BMDCs after two-day stimulation by various constructs and relevant controls. Magnification: 10×. (**B, D**) Free CpG induced severe cell apoptosis and cell death, as detected by Annexin V and PI staining. (**C, E**) Higher concentration of CpG-SQBs showed decreased cell apoptosis and cell death. (**F**) BMDC viability as detected by flow cytometry after 48 hours of co-culture. (**G-J**) Flow cytometry data showing frequencies of CD40, CD86, MHC II and CD80, after 2-day stimulation by CpG-SQB constructs with various spacing. 3.5 nm and 5 nm spacing of CpG on SQB showed significantly increased DC maturation compared to an equivalent amount of free CpG. (**K-M**) Amount of IL-12 and IL-10 secretion in the BMDC culture supernatant after one day of stimulation, as quantified by ELISA. The IL-12/IL-10 ratio, a critical marker of Th1 polarization, revealed that 3.5 nm spacing (CpG2) was optimal. CpG0 refers to SQB without CpG and antigen in this Figure.

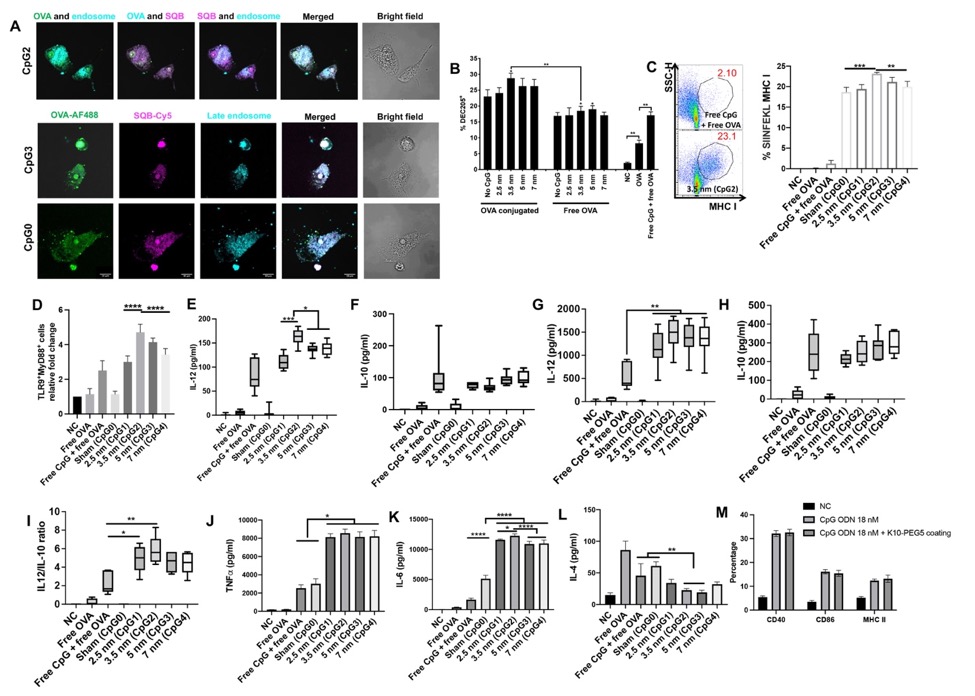

**Fig. S14. BMDCs activation by various DNA origami vaccine constructs and relevant controls.** (**A**) Confocal images showing the colocalization of DoriVac in the endosome and ER. (**B**) Frequency of DEC205^+^ cells in BMDCs treated with DoriVac and various CpG-SQB controls. DEC205 was significantly increased when BMDCs were cultured with CpG-SQB with 3.5 nm CpG spacing. DoriVac co-delivering of antigen and adjuvant also increased DEC205 expression**.** (**C**) Representative SIINFEKL MHC I expressing cell population scatter plots (left panel) and corresponding frequencies in the total cell population. (**D**) Frequencies of double-positive TLR9 and MyD88 expressing population, detected via antibody intracellular staining. CpG2 induced more TLR9 activation and downstream MyD88 expression. (**E**, **F**) ELISA detection of the Th1-polarizing IL-12 cytokine and the Th2-polarizing IL-10 cytokine after 1-day stimulation with various DNA origami vaccine constructs (based on the concentration of 1nM SQB), free OVA (4 nM) or free CpG (18 nM). (**G, H**) ELISA detection of the Th1-polarizing IL-12 cytokine and the Th2-polarizing IL-10 cytokine after 2-day stimulation. (**I**) IL-12 versus IL-10 ratio on day 2. (**J**, **K**) TNFα and IL-6 detection by ELISA within the BMDC supernatant. TNFα expression in all the SQB vaccine groups was significantly higher than the corresponding bolus vaccine (free CpG and free OVA). CpG2 stimulation demonstrated the most increased IL-6 expression compared to controls. (**L**) IL-4 detection by ELISA within the BMDC supernatant. CpG2 and CpG3 stimulation demonstrated the lowest IL-4 expression. (**M**) Positive population (%) of BMDC maturation markers after 1-day stimulation by free CpG or CpG coated with K10-PEG5k. Sham (CpG0) refers to SQB conjugated with OVA antigen but without CpG in this Figure.

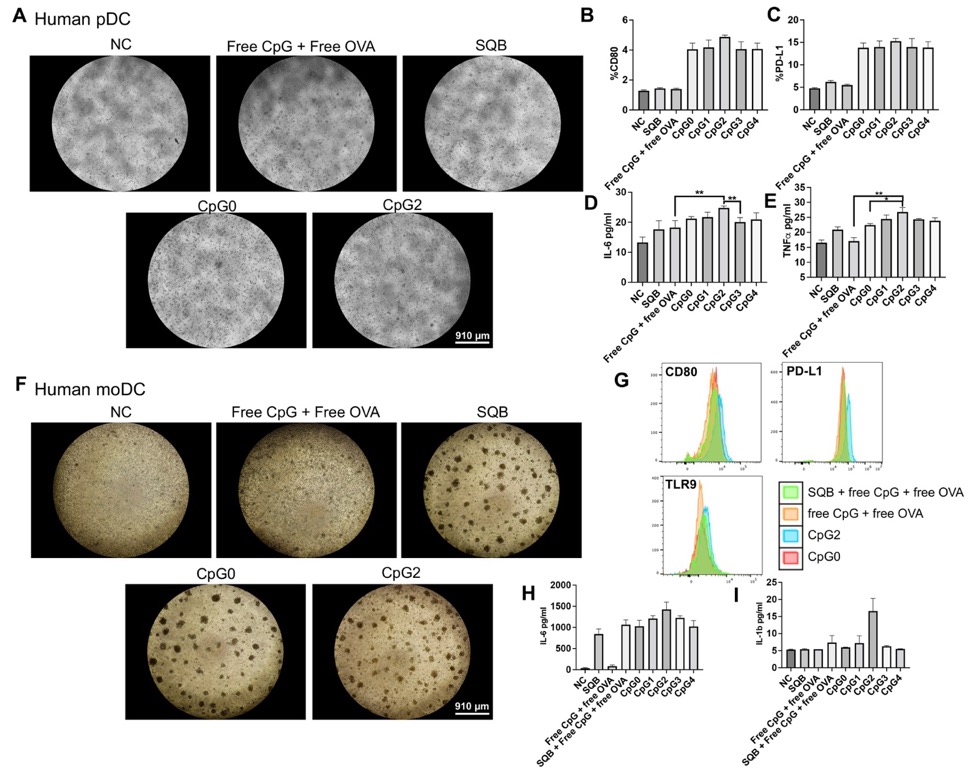

**Fig.S15**. **Spacing effects on human DCs.** (**A**) Representative images showing human plasmacytoid DCs (pDCs) being stimulated under different conditions. Obvious cell clots after stimulation were observed in CpG2 group. All the other spacing showed similar cell morphology compared to CpG2. (**B, C**) Flow results of CD80 and PD-L1 expression on human pDCs. (**D, E**) Luminex results showing IL-6 and TNFα. We did not observe obvious differences in IL-12 and IL-10 cytokine levels likely because the cell number is limited. (**F**) Representative images showing human monocyte derived DCs (moDCs) after various stimulation conditions. (**G**) Flow results of CD80, PD-L1 and TLR9 showed improved cell stimulation by CpG2. All the other spacing showed similar stimulation compared to CpG2. However, the cytokine results showed that CpG2 stimulated the strongest IL-6 (**H**) and IL-1b (**I**) secretion. Various DNA origami vaccine constructs (based on the concentration of 1 nM SQB), free OVA (10 nM) or free CpG (18 nM) was treated.

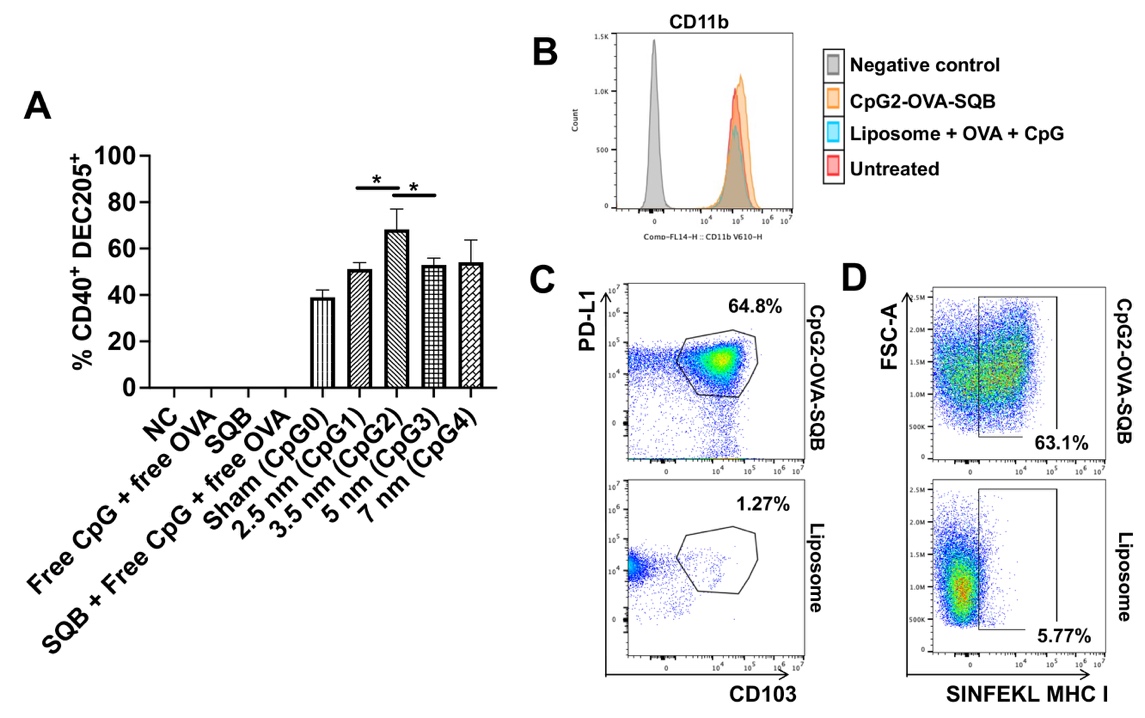

**Fig.S16**. **RAW264.7 cell activation and** **SQB versus liposome in RAW264.7.** (**A**) Double-positive CD40^+^DEC205^+^ population detected in BMDCs by flow cytometry. Comparing DoriVac with liposome carrying the equivalent amounts of CpG and OVA, DoriVac showed enhanced CD11b expression (**B**), as well as PD-L1^+^CD103^+^ population (**C**). (**D**) More SIINFEKL epitopes were presented through MHC I. Various DNA origami vaccine constructs (based on the concentration of 1 nM SQB), free OVA (10 nM) or free CpG (18 nM) was treated.

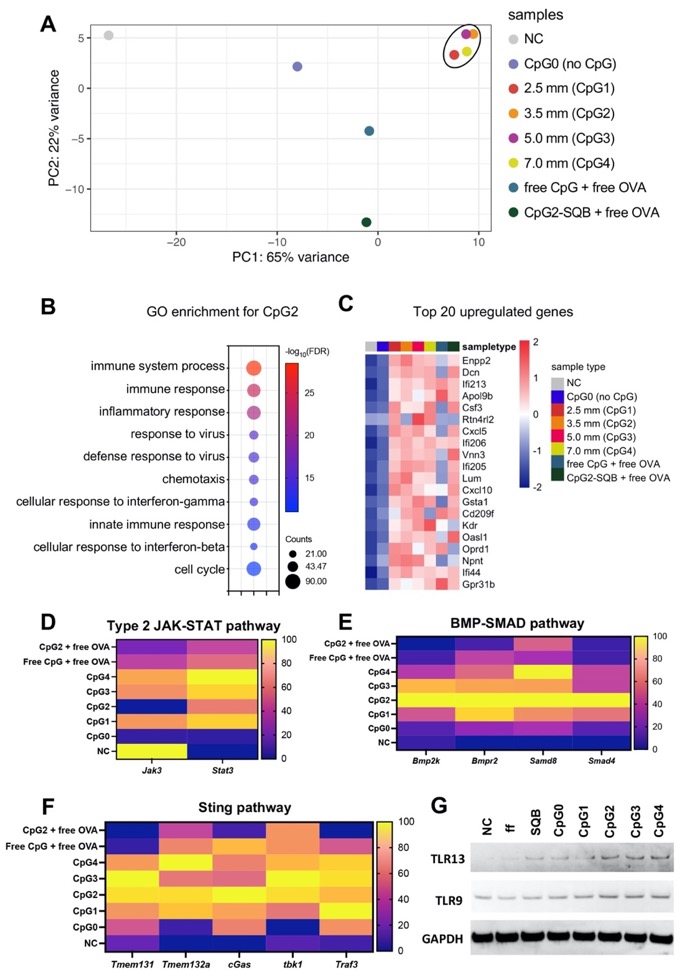

**Fig. S17. RNA-sequencing analysis to characterize transcription changes induced by various origami vaccines.** (**A**) Principal component analysis of the BMDC samples stimulated by various SQB constructs. (**B**) Gene Ontology (GO) enrichment of differentially expressed genes between the CpG2 stimulated group and the unstimulated negative control. (**C**) Heat map of the top 20 differentially upregulated genes in CpG2. (**D-F**) Analysis of related genes in different signaling pathways. (**G**) Western blot results showing the expression of both TLR9 and TLR13. We observe a greater quantity of TLRs were stimulated in CpG2, CpG3 and CpG4 groups (ff: free CpG + free OVA). Various DNA origami vaccine constructs (based on the concentration of 1 nM SQB), free OVA (4 nM) or free CpG (18 nM) was treated.

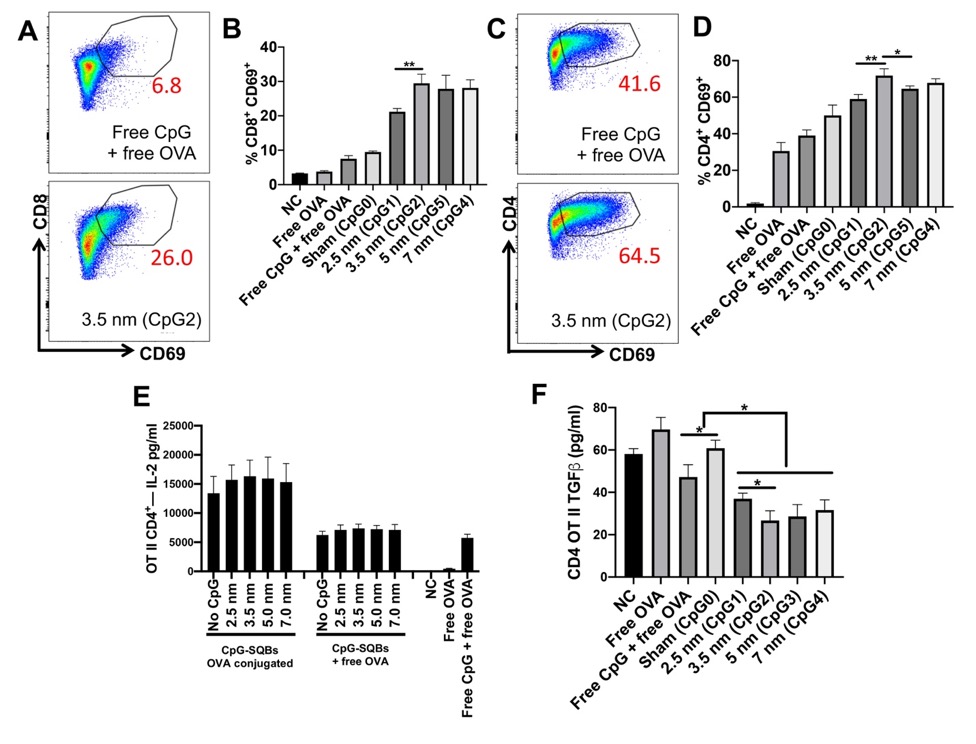

**Fig. S18. T cell activation by various DoriVac and corresponding controls.** (**A-D**) Representative CD69 scatter plots of CD8 and CD4 T cells and corresponding frequencies of CD69-expressing cells. (**E**) ELISA detection of IL-2 in the supernatant of CD4 OT II cells co-cultured with pulsed BMDCs for 2 days. The IL-2 secretion showed negligible difference among different spacings. OVA conjugated CpG-SQBs (left panel) induced the highest IL-2 level. (**F**) ELISA detection of TGFβ in the supernatant of CD4 OT II cells co-cultured with pulsed BMDCs for 2 days. All DoriVac demonstrated less TGFβ secretion compared to bolus vaccine and SQB without CpG (CpG0, OVA conjugated). Various DNA origami vaccine constructs (based on the concentration of 1 nM SQB), free OVA (4 nM) or free CpG (18 nM) was treated.

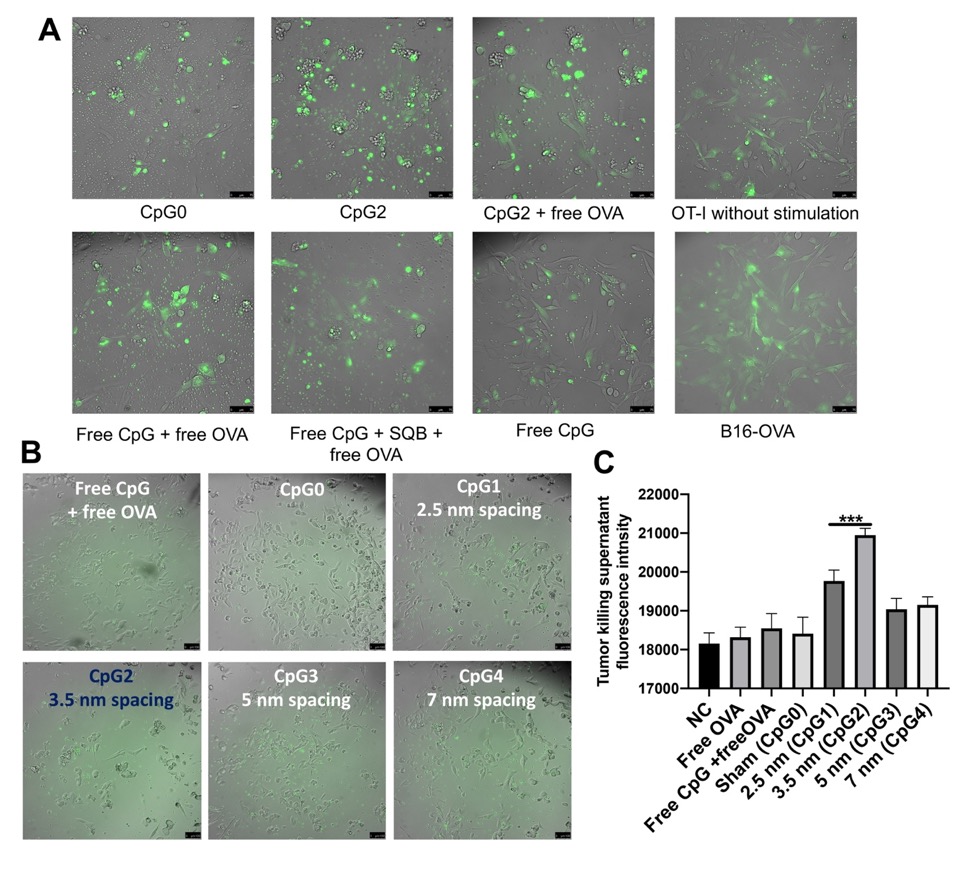

**Fig. S19. *In vitro* tumor killing by vaccine constructs with various CpG spacing.** (**A**) The images of activated T cell - tumor cell co-culture after 2 days. T cells were activated by co-culturing with BMDCs stimulated by DoriVac, control vaccine or unstimulated (as shown in labeling). T cells are imaged as small green dots. The B16-OVA tumor cells were stained with calcein green. Cell spreading is characteristic of healthy tumor cells. Tumor cell aggregations are characteristic of cell killing by CD8 T cells. (**B**) The images represent a different iteration of the experiment. Tumor killing was observed on day 2. Stimulation by DoriVac with various spacings of CpG was compared. (**C**) Calcein green intensity in the supernatant of various groups. Increased calcein is associated with greater T cell activity and increased tumor cell killing, as tumor cell killing leads to increased calcein green release. Various DNA origami vaccine constructs (based on the concentration of 1 nM SQB), free OVA (4 nM) or free CpG (18 nM) was treated.

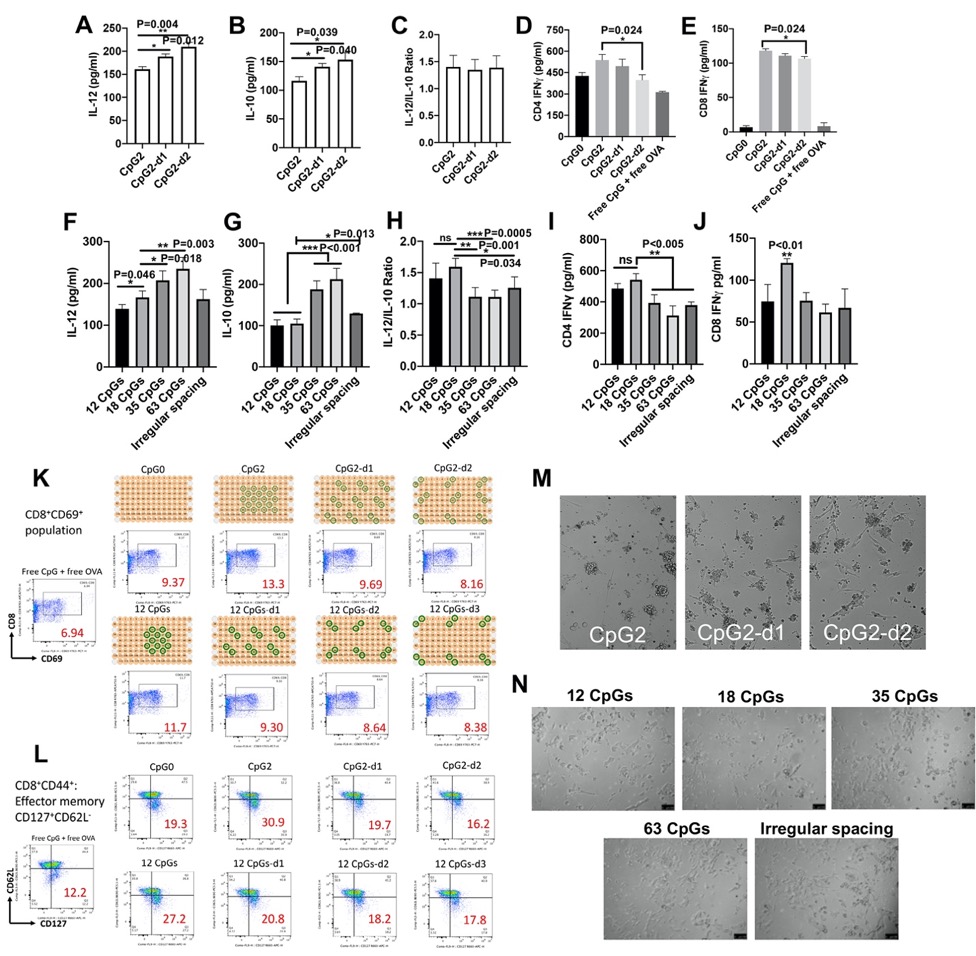

**Fig. S20. DC and T cell activation and *in vitro* tumor killing by various CpG dimer patterns and densities.** DNA origami SQBs are a great tool to study the dimer-trigger unit where two CpG molecules are closely associated in a dimer. Different spacing strategies (as shown in Fig. S4) are easily applied on the SQB platform with controlled CpG orientation to investigate higher-order multimerization. The OT I CD8 T cells were primed by BMDCs treated with various CpG-SQB constructs. (**A**, **B**) ELISA detection of IL-12 and IL-10 secreted from BMDCs after 1-day stimulation. IL-12 and IL-10 secretion both increased when CpG dimer spacing increased. (**C**) The IL-12/IL-10 ratio (n=5). The IL-12/IL-10 ratio was constant as CpG dimer spacing increased, in contrast to the effect observed as CpG monomer spacing increased (decreased IL-12/IL-10 ratio as shown in Fig. 2G) (**D, E**) IFNγ quantification by ELISA from CD4 OT-II and CD8 OT-I T cells after 2-day coculture. (**F, G**) ELISA detection of IL-12 and IL-10 secreted from BMDCs after 1-day stimulation by different number of CpGs on DoriVac. IL-12 and IL-10 secretion both increased in the context of additional CpGs. However, IL-10 secretion was greatly induced with 35 and 63 CpGs. (**H**) The IL-12/IL-10 ratio (n=5). The IL-12/IL-10 ratio decreased as the number of CpGs increased. When the number of CpG was increased from 12 to 18, the IL-12/IL-10 ratio remained constant. Irregular spacing also increased IL-10 secretion and decreased the IL-12/IL-10 ratio compared to CpG2. (**I, J**) Quantification of IFNγ from the supernatant of co-cultured OT-II CD4^+^ or OT-I CD8^+^ T cells with pulsed BMDCs for 2 days by ELISA. 12 CpGs and 18 CpGs showed no IFNγ secretion difference from CD4 OT-II T cells, but 18 CpGs induced more IFNγ secretion from CD8 OT-I T cells. 35, 63 and irregularly spaced CpGs showed less IFNγ expression in both CD4 OT-II and CD8 OT-I T cells, suggesting that 18 CpGs maximizes IFNγ secretion. (**K**) Representative flow plot of CD69 expression (a CD8 T cell activation marker) in different treatment groups. 12 CpGs and other 3 dimer patterns were also examined. (**L**) Representative flow plot of effector memory cells as indicated by the frequency of the effector memory CD8^+^CD44^+^CD127^+^CD62L^-^ population. CD8 T cell activation decreased as the dimer spacing increased, suggesting that 3.5 nm spacing of the CpG dimer is optimal. (**M**) Increased spacing of the CpG dimers decreased the amount of observed tumor-cell killing (aggregated cells) associated with activated CD8 T cells. (**N**) 18 CpGs per SQB demonstrated the most tumor-cell killing activity compared to the relevant controls. Increased CpG density (35 or 63 CpGs per SQB) resulted in decreased tumor killing after sequential co-culture. Decreased CpG density (12 or less CpGs per SQB) also resulted in decreased tumor killing with sequential co-culture, suggesting that 18 CpGs per SQB is optimal for tumor-cell killing activity and aligning with previous results. Various DNA origami vaccine constructs (based on the concentration of 1 nM SQB), free OVA (6 nM) or free CpG (18 nM) was treated.

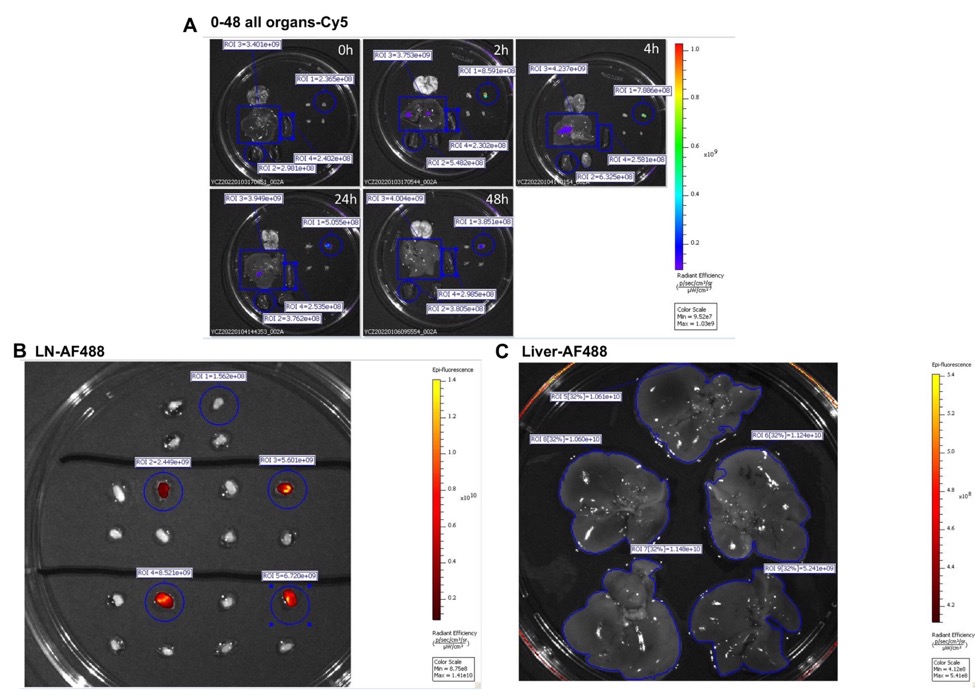

**Fig. S21. DoriVac distribution.** The vaccine was administrated at 0 hour with a concentration of 600 nM in 100 ul on the left shoulder of the C57BL6 mice. The AF488 and Cy5 fluorescent signal was recorded by IVIS at 2, 4, 24 and 48 hours. AF488 was labeled on OVA protein conjugated on the SQB. Cy5 was conjugated directly on the SQB. (**A**) Representative image showing the various organs’ Cy5 signal at different time point. All the results were standardized to avoid individual bias. The schematic of ROI gating was also showed. The vaccine dominantly accumulated in draining lymph nodes (blue circle). (**B**) Representative image showing the LN AF488 signal gating. (**C**) Representative image showing the liver AF488 signal gating.

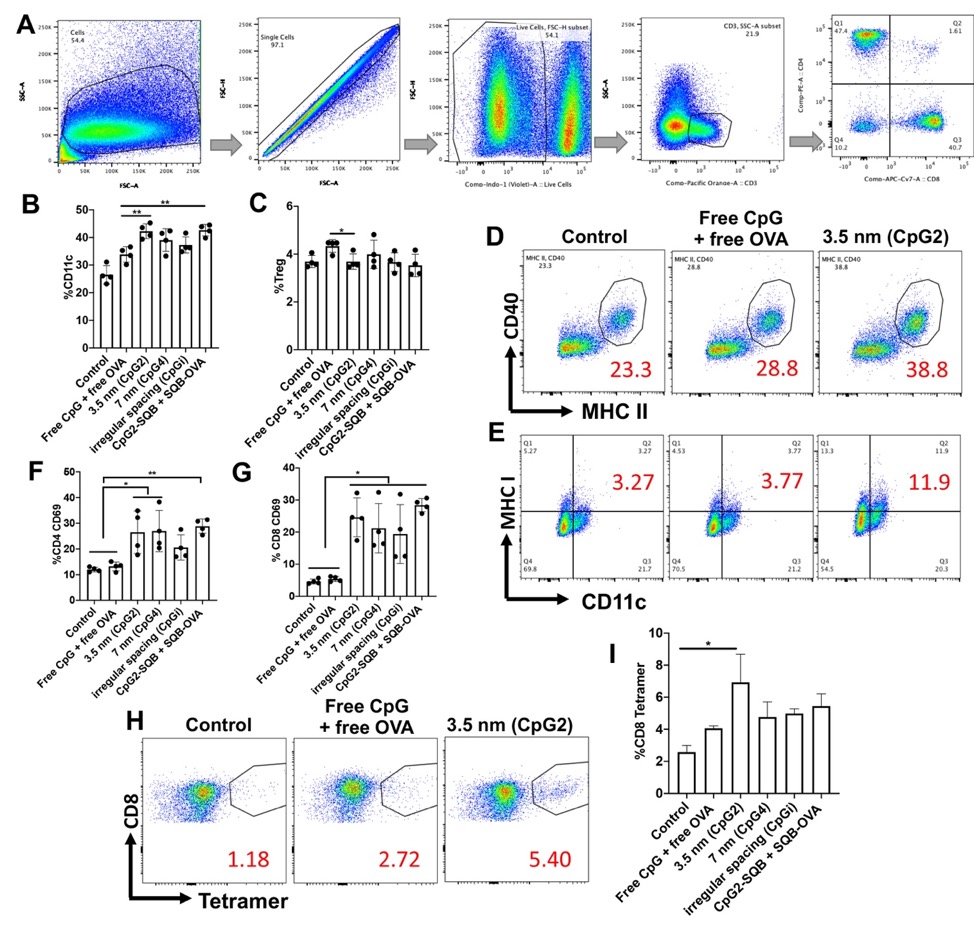

**Fig. S22. Vaccination in naïve healthy mice.** The mice were vaccinated by various versions of the DoriVac and bolus vaccine. The lymph nodes (LNs) were processed into single cell suspension for flow cytometry analysis on day 1 or day 8 after vaccination. Blood cells were also collected on day 8 for CD8 tetramer cell analysis. (**A**) Cell gating strategy for lymphocytes. The subset of CD3^+^ or CD11c^+^ cells was gated from all live cells. Based on CD3 gating, the subpopulations of CD4 and CD8 T cells were gated. (**B, C**) Frequencies of CD11c^+^ DCs and CD4^+^CD25^+^FoxP3^+^ Treg population. (**D, E**) Representative scatter plots of the CD40 and MHC II double positive population and SINFEKL MHC I population of DCs from three different treatment groups. (**F, G**) Frequencies of CD69 positive cells in CD8 and CD4 T cells in the draining LN 1 day after vaccination. (**H**) Representative scatter plots of CD8 tetramer cells found in the LN 8 days after vaccination. Note that DC and T cell activation markers showed no difference compared to untreated control on day 8 after vaccination, suggesting that vaccination boosts are necessary for durable immune responses. (**I**) Frequency of CD8 tetramer population in the blood cells, collected 8 days after vaccination.

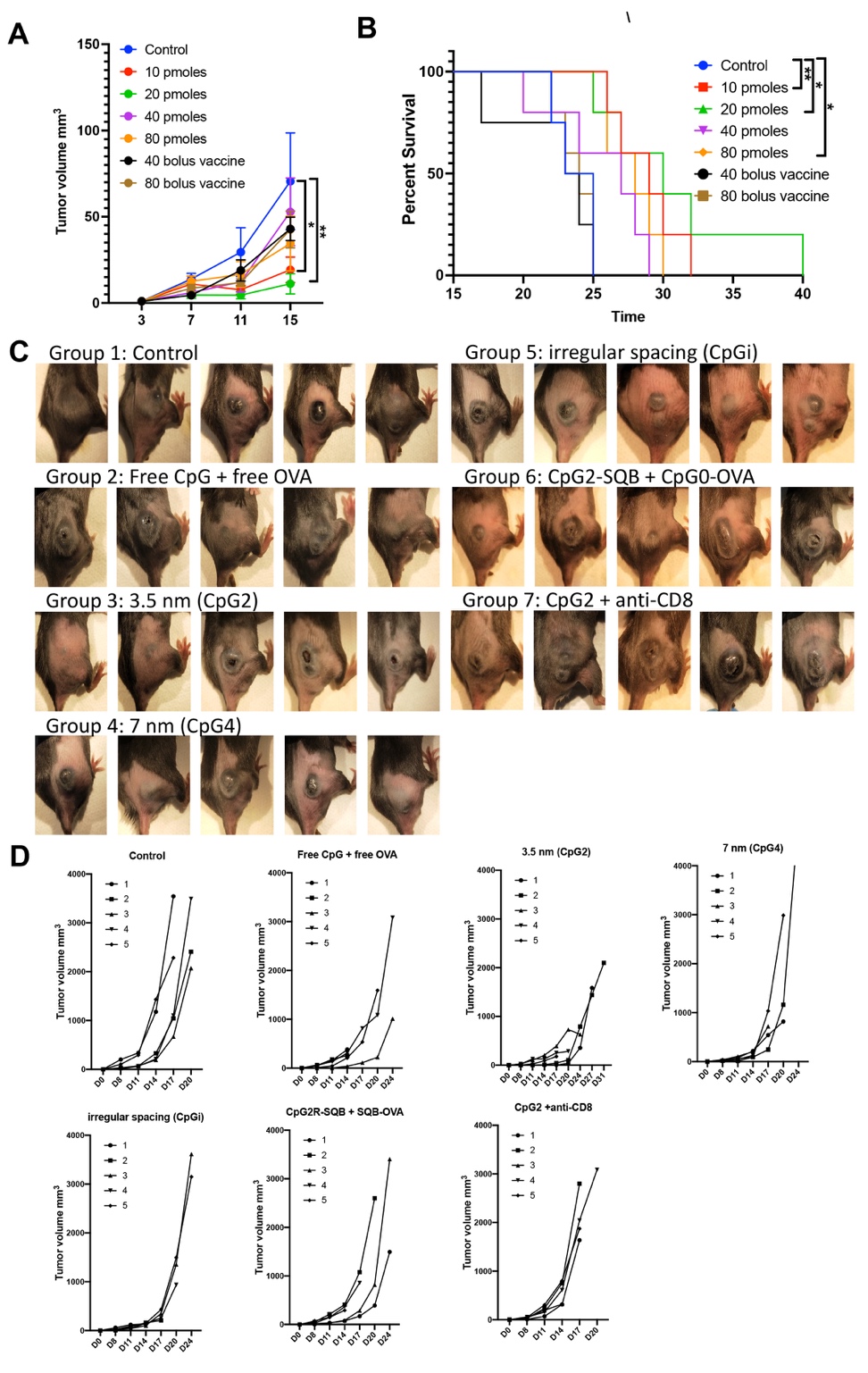

**Fig. S23. Therapeutic vaccination effects of DoriVac in the B16-OVA melanoma tumor model.** (**A,B**) Different vaccine concentrations were tested for a total of three doses in B16-OVA model. 40 and 80 bolus vaccine (Free CpG + Free OVA) corresponding the same amount of antigen and adjuvant in 40 pmoles and 80 pmoles DoriVac. The tumor growth and mice survival results disclose that 10 and 20 pmoles DoriVac induced the most beneficial immune response. Increasingly, the increased DoriVac amount does not lead to further benefits. The bolus vaccines did not show any benefits in this study. (**C**) Full panel images showing tumor growth in various groups 15 days after tumor inoculation. (**D**) Tumor growth curves were plotted for each individual mouse in each group.

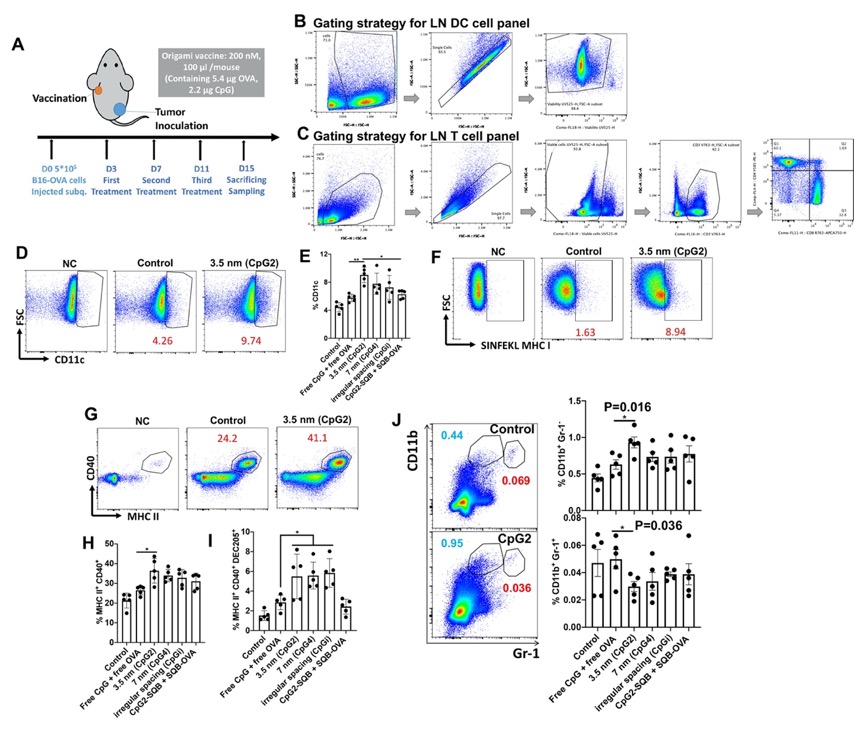

**Fig. S24. Additional lymphocyte analysis from treated tumor-bearing mice.** (**A**) For immune profiling experiments, mice were treated with vaccines on day 3, 7 and 11, and then sacrificed on day 15. This is a slightly different vaccination strategy than was used for the treatment experiments to ensure that all mice were alive at the time of sacrifice. (**B**) Cell gating strategy for lymphocyte DC panel. (**C**) Cell gating strategy for lymphocyte T cell panels. Live cells were gated for CD3^+^ cells. CD4 and CD8 T cells were gated based on the CD3 gating, and then subsequently gated for T cell subpopulations. (**D, E**) Representative images from flow scatter plots and corresponding frequencies of CD11c positive population. (**F**) Representative scatter plots of SINFEKL MHC I population. (**G, H**) Representative scatter plots and corresponding frequencies of CD40 and MHC II double positive population. CpG2 showed the most significant DC maturation, as demonstrated by the frequency of CD40^+^MHC II^+^. (**I**) DEC205^+^ subset was gated out of CD40^+^ MHC II^+^ cells. Codelivery of antigen with CpG adjuvant through SQB could greatly increase antigen uptake indicated by enhanced DEC205 expression compared to OVA delivered freely. **(J)** Representative scatter plots (left) and percentage quantifications (right) of myeloid-derived suppressor CD11b^+^Gr-1^+^ cells (red) and CD11b^+^Gr-1^-^ monocytes (blue) in the LN.

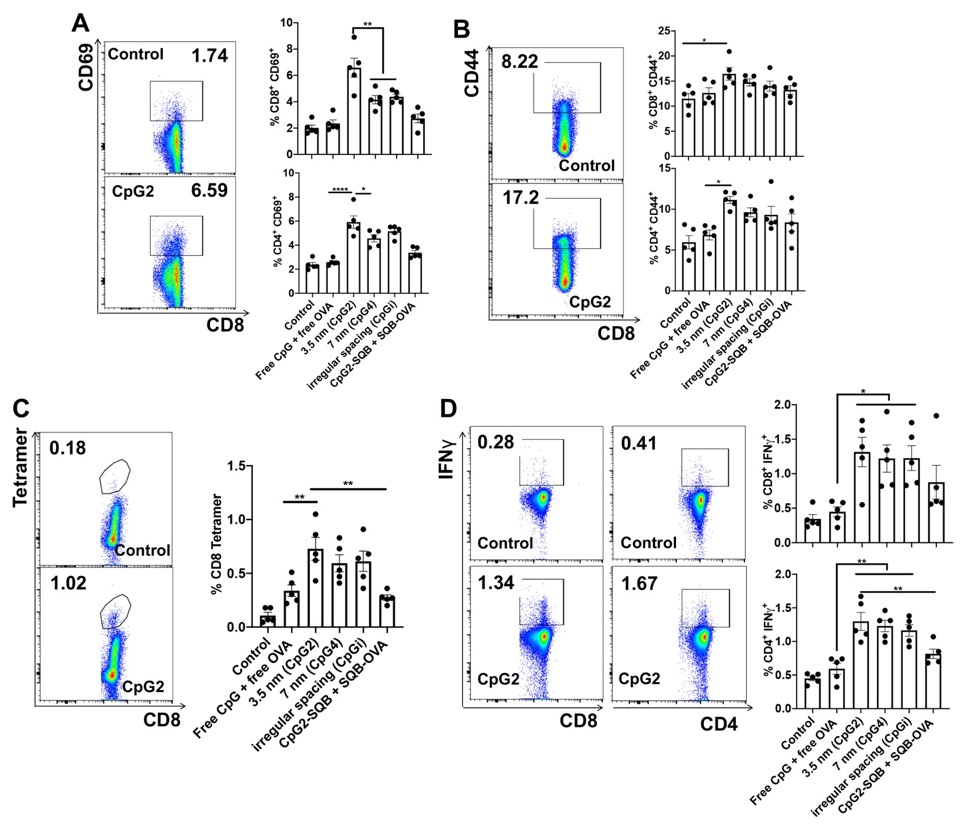

**Fig. S25. Additional lymphocyte T cell analysis from treated tumor-bearing mice.** (**A**) Representative CD69^+^ scatter plots of CD8^+^ cells (left) and percentages of CD69^+^ cells in CD8^+^ and CD4^+^ T cells (right). (**B**) Representative CD44^+^ plots of CD8^+^ cells (left) and percentages of CD44^+^ cells in CD8^+^ and CD4^+^ T cells (right). (**C**) Representative SIINFEKL tetramer plots on CD8^+^ cells (left) and percentages of tetramer cells in CD8^+^ T cells (right). (**D**) Representative IFNγ^+^ scatter plots of CD8^+^ or CD4^+^ T cells (left) and percentages of IFNγ^+^ cells in CD8^+^ or CD4^+^ T cells (right). IFNγ-expressing CD4 Th1 cells and CD8 cell were both significantly increased in all the origami vaccine groups co-delivering OVA and CpG.

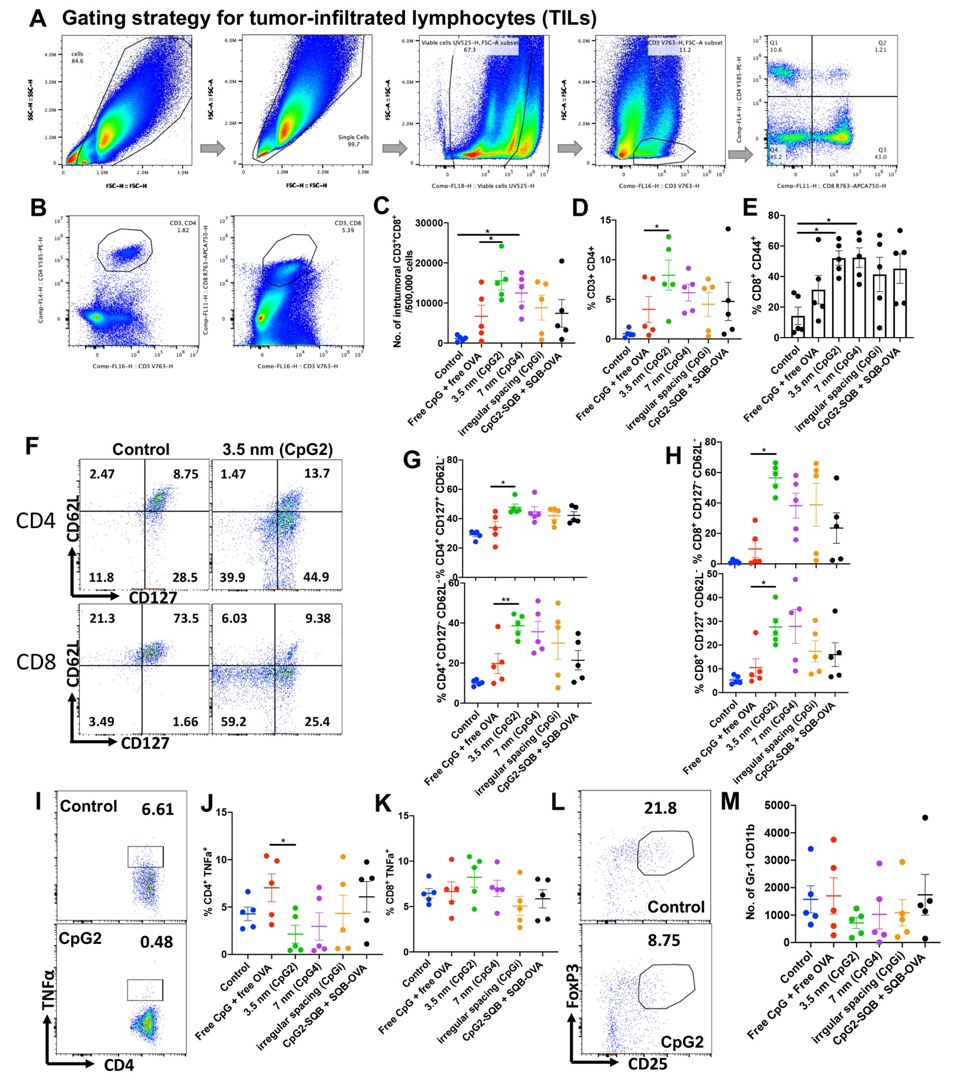

**Fig. S26. Extensive tumor infiltrating lymphocytes (TILs) analysis from treated tumor-bearing mice.** (**A**) The tumor tissue was collected on day 15 and processed into single cell suspension for flow cytometry. Cell gating strategy were shown. CD3^+^ cells were gated out of live cells directly. Based on CD3 gating, CD4 and CD8 T cells were gated for T cell subpopulations. (**B**) In some cases, CD3^+^CD4^+^ or CD3^+^CD8^+^ cells were gated separately. (**C, D**) Number or frequency of infiltrated CD4 and CD8 T cell populations. (**E**) CpG2 and CpG4 groups showed increased frequency of the CD8^+^CD44^+^ memory cell population. (**F**) Gating for T memory cell sub-populations by CD62L and CD127. (**G**) CpG2 showed significantly increased T effector memory CD4^+^CD127^+^CD62L^-^ T cell and effector CD4^+^CD127^-^CD62L^-^ T cell population compared to relevant controls. (**H**) CpG2 showed significantly increased CD8^+^CD127^-^CD62L^-^ effector and effector memory CD8^+^CD127^+^CD62L^-^ T cell population compared to relevant controls. (**I**) Representative scatter plots of CD4^+^ TNFα^+^ population. (**J**) The CD4^+^ TNFα^+^ population was significantly downregulated in CpG2 group. (**K**) The CD8^+^ TNFα^+^ subpopulation frequency did not show an obvious difference among various groups. (**L**) Representative scatter plots of the CD4^+^ Treg subpopulation in the tumor environment of treated mice. (**M**) Quantification of Gr-1^+^CD11b^+^ myeloid-derived suppressor cells in tumor tissue.

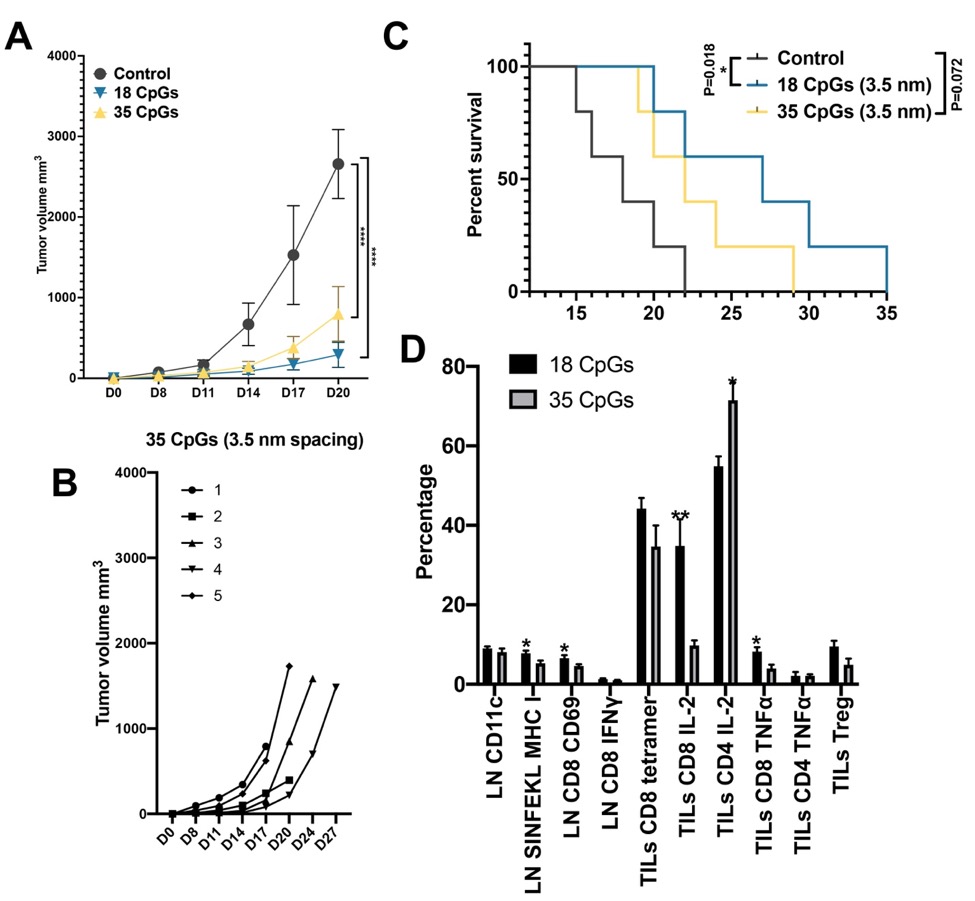

**Fig. S27. 35 CpGs at 3.5 nm spacing showed reduced therapeutic effects compared to 18 CpGs at the same spacing when administered on the DNA origami SQB platform.** (**A**) Tumor growth was not significantly different between 18 CpGs and 35 CpGs treated groups. (**B**) Tumor growth curves of all 5 mice treated with the 35 CpGs per SQB vaccine showed similar profiles (Tumor growth curve of mice treated with 18 CpGs per SQB vaccine is shown in Fig. S23E: CpG2). (**C**) Survival curves of the DoriVac treated groups with 35 CpGs compared to 18 CpGs. The DoriVac with 35 CpGs per SQB group had median survival of 22 days versus 26 days in 18 CpGs group. (**D**) Immune cell profiling showed significant differences between the different treatment groups. The differences were mainly associated with cross-presentation and CD8 T cell response. 35 CpGs induced less CD8 activation but increased the CD4^+^ IL-2^+^ population.

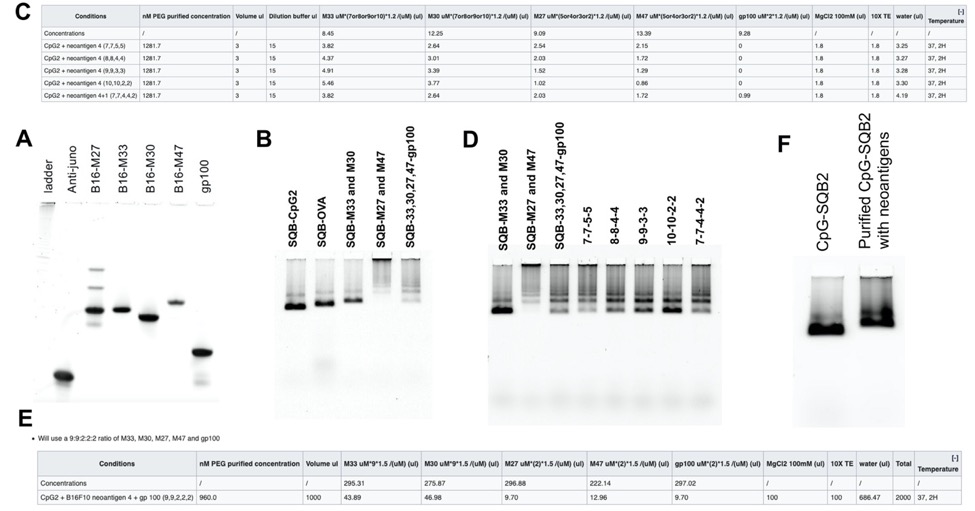

**Fig. S28. Fabrication of neoantigen vaccines.** (**A**) Denaturing page gel showing conjugated neoantigen peptides or tumor associated peptide (gp100) with single stranded oligo DNA using DBCO-Azide click chemistry. These products were purified through band cutting and ethanol precipitation. (**B**) Agarose gel result showing conjugation of neoantigens with CpG2-SQB. Conjugation with M33 and M30 did not lead to aggregation while, conjugation with M27 and M47 did. (**C**) A protocol showing how different proportions of the peptides were tested, while maintaining a constant total number of MHC I and MHC II peptides identical. (**D**) Agarose gel result showing the testing of different proportions of peptides. (**E**) A protocol demonstrating how we fabricated the vaccine eventually for the animal study. (**F**) Agarose gel result showing purified neoantigen DoriVac compared to the CpG2-SQB.

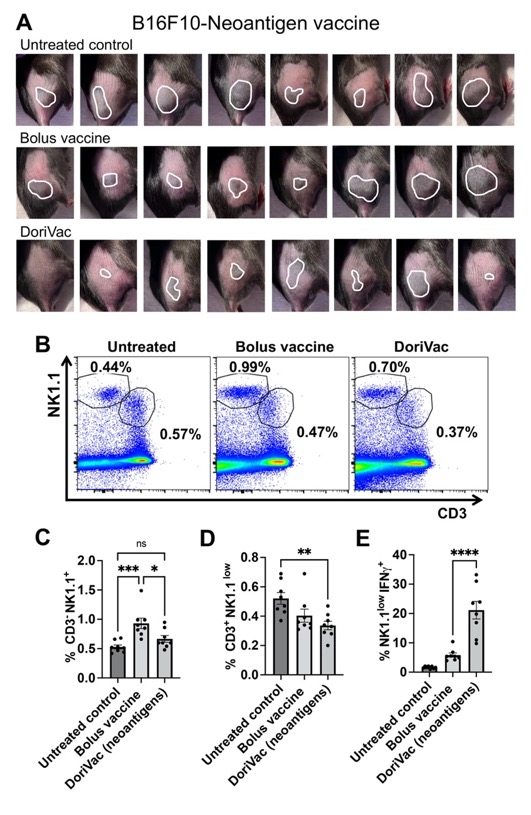

**Fig. S29. Neoantigen modified DoriVac treatment in B16F10 melanoma model.**(**A**) Tumor images were taken on day 12 post tumor inoculation. (**B-D**) Representative scatter plots and percentages of NK cells in the draining LN. NK1.1 and CD3 were plotted together to evaluate the NK1.1^+^CD3^-^ and NK1.1^low^CD3^+^ populations. (**E**) Percentages of IFNγ expression NK cells in the NK1.1^low^CD3^+^ population.

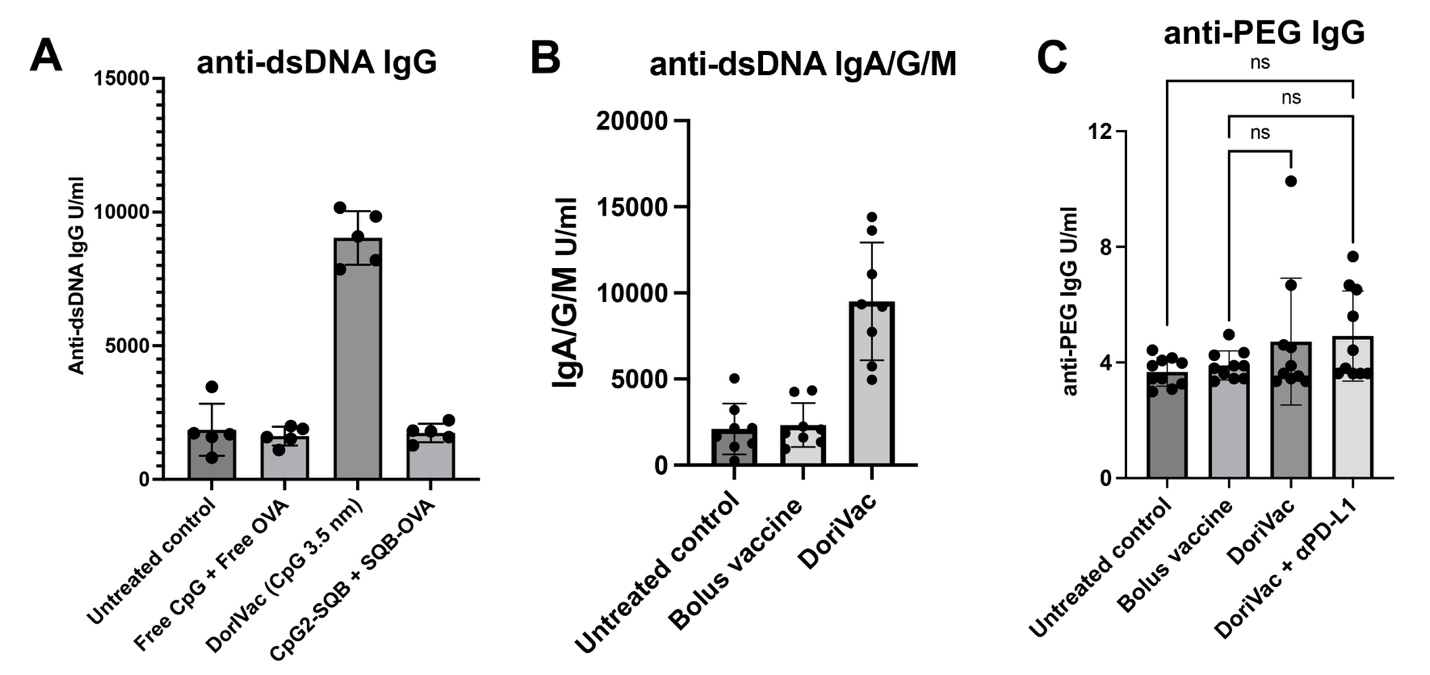

**Fig. S30. Immunogenicity of DNA origami.** (**A**) The anti-dsDNA IgG antibody titer was measured by ELISA after administering 3 doses of 20 pmol DoriVac in the B16OVA tumor model. Note that the group of CpG2-SQB + SQB-OVA was treated with 40 pmole of SQB nanoparticles to include same amount of OVA and CpG with DoriVac. However, this group didn’t show increased anti-dsDNA IgG, indicating that the anti-dsDNA IgG might not be correlated with DNA origami nanoparticle used for treatment. By analyzing anti-tumor effect occurred by DoriVac, the increase in anti-dsDNA IgG may be attributed to their anti-tumor effects. (**B**) The anti-dsDNA IgA/G/M antibody titer in the blood was measured after administering 3 doses of 20 pmol DoriVac in the B16F10 tumor model. The titer showed a four-fold increase compared to the control and bolus vaccine groups. (**C**) The anti-PEG IgG antibody titer was measured by ELISA after administering 3 doses of 20 pmol DoriVac in the B16OVA tumor model. No significant elevation of anti-PEG IgG was observed in mouse serum.

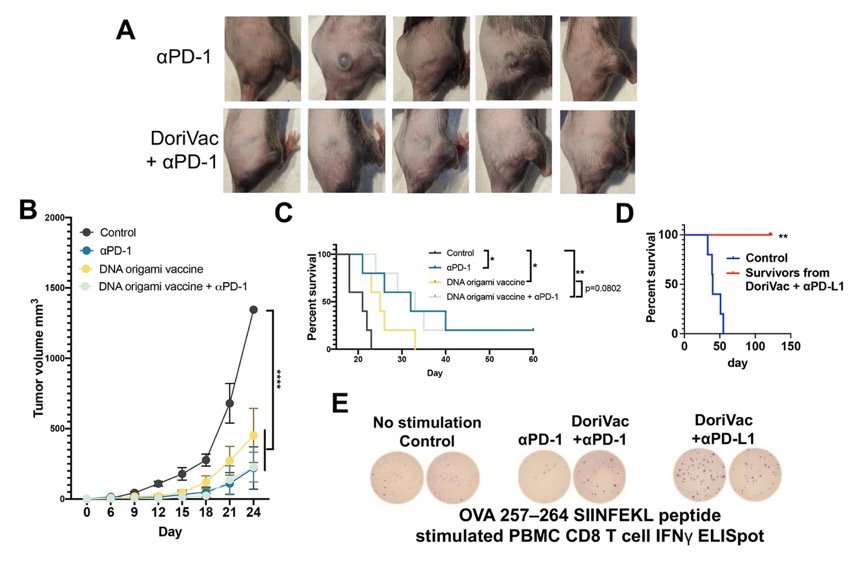

**Fig. S31. DoriVac treatment combined with αPD-L1 or αPD-1 for melanoma.** (**A**) Tumor images were taken on day 12 post tumor inoculation. (**B**) Tumor growth curve. DoriVac combination with αPD-1 did not exhibit synergy effects. αPD-1 was applied at the same time points and doses as αPD-L1. (**C**) Percent survival of B16-OVA tumor-bearing mice in various treatment groups. The median survival of untreated control, DoriVac treated, αPD-1treated and combination treated groups are 21, 25, 32, and 33 days, respectively. One mouse from the αPD-1 monotherapy group and one mouse from the combination group showed durable tumor regression and survived. n=5. (**D**) The survival curve of mice from DoriVac and αPD-L1 combination group compared to controls. The survived mice were rechallenged with 2×10^5^ B16-OVA tumor cells after 4 months. All the survived mice from the treatment groups (αPD-1(n=1), DoriVac + αPD-1(n=1) and DoriVac + αPD-L1 (n=4)) showed complete tumor remission after rechallenging. (**E**) One year after tumor rechallenging, before we sacrificed the mice, we took blood samples and performed an IFNγ ELISpot experiment on the survived mice PBMCs, stimulating with SIINFEKL peptides to look at the quantity of antigen-specific T cell responses. All the DoriVac applied groups showed obvious CD8 T cell responses as evidenced by the IFNγ spots. Two mice from DoriVac + αPD-L1 group were sacrificed in advance for student practice purposes and were not able to be tested for ELISpot.

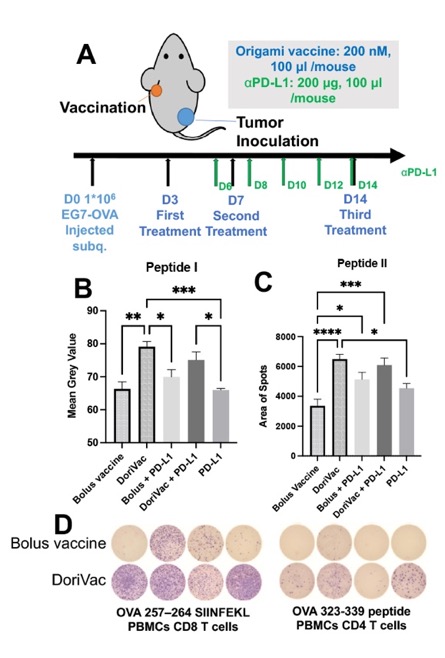

**Fig. S32. DoriVac treatment in EG7-OVA lymphoma model.**(**A**) Schematic delineating mouse EG7-OVA lymphoma model setup and therapeutic vaccination treatment plan in C57BL/6 mice. (**B, C**) Statistical summary of the splenocyte ELISpot results of Fig. 6G, H. (**D**) ELISpot results showing the memory CD8 T cells in PBMCs 6 months after the tumor rechallenge in EG7-OVA model, comparing the bolus vaccine and DoriVac group.

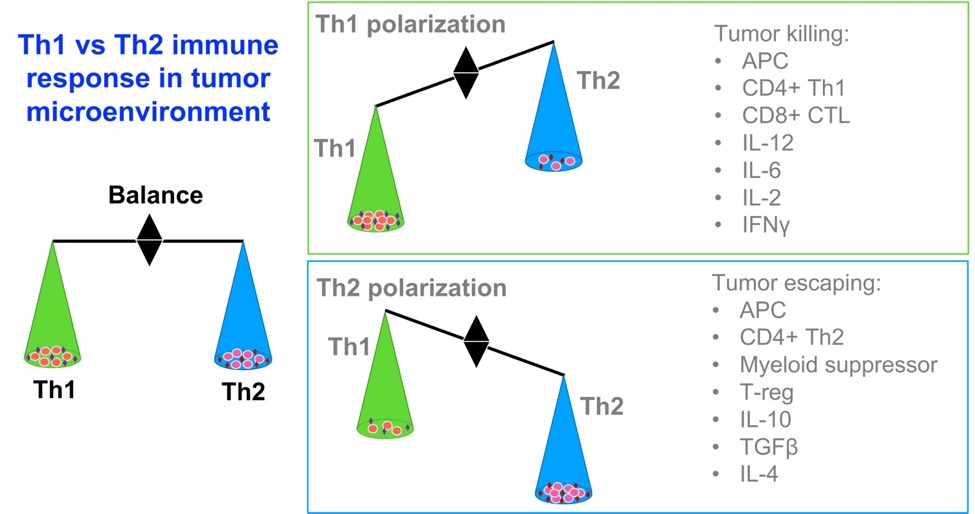

**Fig. S33. Summary of Th1 versus Th2 immune polarization. (A)** Conventionally, type I (Th1) and type II (Th2) refer to two categories of CD4 positive T helper cells. However, in the context of the tumor immune microenvironment, Th1 and Th2 refer to two types of immune polarization characterized by specific infiltrating cells and cytokines. Th1 polarized immune responses are associated with increased DC maturation, Th1 polarization of CD4 cells, CD8-mediated killing of tumor cells and increased cancer regression. IL-12, IL-6, IFNγ and IL-2 are identified as critical Th1-polarizing cytokines. In the context of Th2 immune polarization, the tumor can escape immune surveillance. The Th2-polarized microenvironment is characterized by Th2-polarized CD4 cells, T regulatory cells, myeloid suppressor cells and cytokines IL-10, TGFβ and IL-4, which serve to dampen the anti-tumor response. One factor that may affect Th1 or Th2 immune polarization is the choice of immune adjuvant, such as CpG and dsRNA. In this experiment, SQBs co-delivering antigen and adjuvant CpG at optimal spacing induces improved Th1-polarized immune response which is beneficial for anti-tumor immunity.

**DNA scaffold for SQB|** p8634 sequence

AATGCTACTACTATTAGTAGAATTGATGCCACCTTTTCAGCTCGCGCCCCAAATGAAAATATAGCTAAACAGGTTATTGACCATTTGCGAAATGTATCTAATGGTCAAACTAAATCTACTCGTTCGCAGAATTGGGAATCAACTGTTATATGGAATGAAACTTCCAGACACCGTACTTTAGTTGCATATTTAAAACATGTTGAGCTACAGCATTATATTCAGCAATTAAGCTCTAAGCCATCCGSCAAAAATGACCTCTTATCAAAAGGAGCAATTAAAGGTACTCTCTAATCCTGACCTGTTGGAGTTTGCTTCCGGTCTGGTTCGCTTTGAAGCTCGAATTAAAACGCGATATTTGAAGTCTTTCGGGCTTCCTCTTAATCTTTTTGATGCAATCCGCTTTGCTTCTGACTATAATAGTCAGGGTAAAGACCTGATTTTTGATTTATGGTCATTCTCGTTTTCTGAACTGTTTAAAGCATTTGAGGGGGATTCAATGAATATTTATGACGATTCCGCAGTATTGGACGCTATCCAGTCTAAACATTTTACTATTACCCCCTCTGGCAAAACTTCTTTTGCAAAAGCCTCTCGCTATTTTGGTTTTTATCGTCGTCTGGTAAACGAGGGTTATGATAGTGTTGCTCTTACTATGCCTCGTAATTCCTTTTGGCGTTATGTATCTGCATTAGTTGAATGTGGTATTCCTAAATCTCAACTGATGAATCTTTCTACCTGTAATAATGTTGTTCCGTTAGTTCGTTTTATTAACGTAGATTTTTCTTCCCAACGTCCTGACTGGTATAATGAGCCAGTTCTTAAAATCGCATAAGGTAATTCACAATGATTAAAGTTGAAATTAAACCATCTCAAGCCCAATTTACTACTCGTTCTGGTGTTTCTCGTCAGGGCAAGCCTTATTCACTGAATGAGCAGCTTTGTTACGTTGATTTGGGTAATGAATATCCGGTTCTTGTCAAGATTACTCTTGATGAAGGTCAGCCAGCCTATGCGCCTGGTCTGTACACCGTTCATCTGTCCTCTTTCAAAGTTGGTCAGTTCGGTTCCCTTATGATTGACCGTCTGCGCCTCGTTCCGGCTAAGTAACATGGAGCAGGTCGCGGATTTCGACACAATTTATCAGGCGATGATACAAATCTCCGTTGTACTTTGTTTCGCGCTTGGTATAATCGCTGGGGGTCAAAGATGAGTGTTTTAGTGTATTCTTTTGCCTCTTTCGTTTTAGGTTGGTGCCTTCGTAGTGGCATTACGTATTTTACCCGTTTAATGGAAACTTCCTCATGAAAAAGTCTTTAGTCCTCAAAGCCTCTGTAGCCGTTGCTACCCTCGTTCCGATGCTGTCTTTCGCTGCTGAGGGTGACGATCCCGCAAAAGCGGCCTTTAACTCCCTGCAAGCCTCAGCGACCGAATATATCGGTTATGCGTGGGCGATGGTTGTTGTCATTGTCGGCGCAACTATCGGTATCAAGCTGTTTAAGAAATTCACCTCGAAAGCAAGCTGATAAACCGATACAATTAAAGGCTCCTTTTGGAGCCTTTTTTTTGGAGATTTTCAACGTGAAAAAATTATTATTCGCAATTCCTTTAGTTGTTCCTTTCTATTCTCACTCCGCTGAAACTGTTGAAAGTTGTTTAGCAAAATCCCATACAGAAAATTCATTTACTAACGTCTGGAAAGACGACAAAACTTTAGATCGTTACGCTAACTATGAGGGCTGTCTGTGGAATGCTACAGGCGTTGTAGTTTGTACTGGTGACGAAACTCAGTGTTACGGTACATGGGTTCCTATTGGGCTTGCTATCCCTGAAAATGAGGGTGGTGGCTCTGAGGGTGGCGGTTCTGAGGGTGGCGGTTCTGAGGGTGGCGGTACTAAACCTCCTGAGTACGGTGATACACCTATTCCGGGCTATACTTATATCAACCCTCTCGACGGCACTTATCCGCCTGGTACTGAGCAAAACCCCGCTAATCCTAATCCTTCTCTTGAGGAGTCTCAGCCTCTTAATACTTTCATGTTTCAGAATAATAGGTTCCGAAATAGGCAGGGGGCATTAACTGTTTATACGGGCACTGTTACTCAAGGCACTGACCCCGTTAAAACTTATTACCAGTACACTCCTGTATCATCAAAAGCCATGTATGACGCTTACTGGAACGGTAAATTCAGAGACTGCGCTTTCCATTCTGGCTTTAATGAGGATTTATTTGTTTGTGAATATCAAGGCCAATCGTCTGACCTGCCTCAACCTCCTGTCAATGCTGGCGGCGGCTCTGGTGGTGGTTCTGGTGGCGGCTCTGAGGGTGGTGGCTCTGAGGGTGGCGGTTCTGAGGGTGGCGGCTCTGAGGGAGGCGGTTCCGGTGGTGGCTCTGGTTCCGGTGATTTTGATTATGAAAAGATGGCAAACGCTAATAAGGGGGCTATGACCGAAAATGCCGATGAAAACGCGCTACAGTCTGACGCTAAAGGCAAACTTGATTCTGTCGCTACTGATTACGGTGCTGCTATCGATGGTTTCATTGGTGACGTTTCCGGCCTTGCTAATGGTAATGGTGCTACTGGTGATTTTGCTGGCTCTAATTCCCAAATGGCTCAAGTCGGTGACGGTGATAATTCACCTTTAATGAATAATTTCCGTCAATATTTACCTTCCCTCCCTCAATCGGTTGAATGTCGCCCTTTTGTCTTTGGCGCTGGTAAACCATATGAATTTTCTATTGATTGTGACAAAATAAACTTATTCCGTGGTGTCTTTGCGTTTCTTTTATATGTTGCCACCTTTATGTATGTATTTTCTACGTTTGCTAACATACTGCGTAATAAGGAGTCTTAATCATGCCAGTTCTTTTGGGTATTCCGTTATTATTGCGTTTCCTCGGTTTCCTTCTGGTAACTTTGTTCGGCTATCTGCTTACTTTTCTTAAAAAGGGCTTCGGTAAGATAGCTATTGCTATTTCATTGTTTCTTGCTCTTATTATTGGGCTTAACTCAATTCTTGTGGGTTATCTCTCTGATATTAGCGCTCAATTACCCTCTGACTTTGTTCAGGGTGTTCAGTTAATTCTCCCGTCTAATGCGCTTCCCTGTTTTTATGTTATTCTCTCTGTAAAGGCTGCTATTTTCATTTTTGACGTTAAACAAAAAATCGTTTCTTATTTGGATTGGGATAAATAATATGGCTGTTTATTTTGTAACTGGCAAATTAGGCTCTGGAAAGACGCTCGTTAGCGTTGGTAAGATTCAGGATAAAATTGTAGCTGGGTGCAAAATAGCAACTAATCTTGATTTAAGGCTTCAAAACCTCCCGCAAGTCGGGAGGTTCGCTAAAACGCCTCGCGTTCTTAGAATACCGGATAAGCCTTCTATATCTGATTTGCTTGCTATTGGGCGCGGTAATGATTCCTACGATGAAAATAAAAACGGCTTGCTTGTTCTCGATGAGTGCGGTACTTGGTTTAATACCCGTTCTTGGAATGATAAGGAAAGACAGCCGATTATTGATTGGTTTCTACATGCTCGTAAATTAGGATGGGATATTATTTTTCTTGTTCAGGACTTATCTATTGTTGATAAACAGGCGCGTTCTGCATTAGCTGAACATGTTGTTTATTGTCGTCGTCTGGACAGAATTACTTTACCTTTTGTCGGTACTTTATATTCTCTTATTACTGGCTCGAAAATGCCTCTGCCTAAATTACATGTTGGCGTTGTTAAATATGGCGATTCTCAATTAAGCCCTACTGTTGAGCGTTGGCTTTATACTGGTAAGAATTTGTATAACGCATATGATACTAAACAGGCTTTTTCTAGTAATTATGATTCCGGTGTTTATTCTTATTTAACGCCTTATTTATCACACGGTCGGTATTTCAAACCATTAAATTTAGGTCAGAAGATGAAATTAACTAAAATATATTTGAAAAAGTTTTCTCGCGTTCTTTGTCTTGCGATTGGATTTGCATCAGCATTTACATATAGTTATATAACCCAACCTAAGCCGGAGGTTAAAAAGGTAGTCTCTCAGACCTATGATTTTGATAAATTCACTATTGACTCTTCTCAGCGTCTTAATCTAAGCTATCGCTATGTTTTCAAGGATTCTAAGGGAAAATTAATTAATAGCGACGATTTACAGAAGCAAGGTTATTCACTCACATATATTGATTTATGTACTGTTTCCATTAAAAAAGGTAATTCAAATGAAATTGTTAAATGTAATTAATTTTGTTTTCTTGATGTTTGTTTCATCATCTTCTTTTGCTCAGGTAATTGAAATGAATAATTCGCCTCTGCGCGATTTTGTAACTTGGTATTCAAAGCAATCAGGCGAATCCGTTATTGTTTCTCCCGATGTAAAAGGTACTGTTACTGTATATTCATCTGACGTTAAACCTGAAAATCTACGCAATTTCTTTATTTCTGTTTTACGTGCAAATAATTTTGATATGGTAGGTTCTAACCCTTCCATTATTCAGAAGTATAATCCAAACAATCAGGATTATATTGATGAATTGCCATCATCTGATAATCAGGAATATGATGATAATTCCGCTCCTTCTGGTGGTTTCTTTGTTCCGCAAAATGATAATGTTACTCAAACTTTTAAAATTAATAACGTTCGGGCAAAGGATTTAATACGAGTTGTCGAATTGTTTGTAAAGTCTAATACTTCTAAATCCTCAAATGTATTATCTATTGACGGCTCTAATCTATTAGTTGTTAGTGCTCCTAAAGATATTTTAGATAACCTTCCTCAATTCCTTTCAACTGTTGATTTGCCAACTGACCAGATATTGATTGAGGGTTTGATATTTGAGGTTCAGCAAGGTGATGCTTTAGATTTTTCATTTGCTGCTGGCTCTCAGCGTGGCACTGTTGCAGGCGGTGTTAATACTGACCGCCTCACCTCTGTTTTATCTTCTGCTGGTGGTTCGTTCGGTATTTTTAATGGCGATGTTTTAGGGCTATCAGTTCGCGCATTAAAGACTAATAGCCATTCAAAAATATTGTCTGTGCCACGTATTCTTACGCTTTCAGGTCAGAAGGGTTCTATCTCTGTTGGCCAGAATGTCCCTTTTATTACTGGTCGTGTGACTGGTGAATCTGCCAATGTAAATAATCCATTTCAGACGATTGAGCGTCAAAATGTAGGTATTTCCATGAGCGTTTTTCCTGTTGCAATGGCTGGCGGTAATATTGTTCTGGATATTACCAGCAAGGCCGATAGTTTGAGTTCTTCTACTCAGGCAAGTGATGTTATTACTAATCAAAGAAGTATTGCTACAACGGTTAATTTGCGTGATGGACAGACTCTTTTACTCGGTGGCCTCACTGATTATAAAAACACTTCTCAGGATTCTGGCGTACCGTTCCTGTCTAAAATCCCTTTAATCGGCCTCCTGTTTAGCTCCCGCTCTGATTCTAACGAGGAAAGCACGTTATACGTGCTCGTCAAAGCAACCATAGTACGCGCCCTGTAGCGGCGCATTAAGCGCGGCGGGTGTGGTGGTTACGCGCAGCGTGACCGCTACACTTGCCAGCGCCCTAGCGCCCGCTCCTTTCGCTTTCTTCCCTTCCTTTCTCGCCACGTTCGCCGGCTTTCCCCGTCAAGCTCTAAATCGGGGGCTCCCTTTAGGGTTCCGATTTAGTGCTTTACGGCACCTCGACCCCAAAAAACTTGATTTGGGTGATGGTTCACGTAGTGGGCCATCGCCCTGATAGACGGTTTTTCGCCCTTTGACGTTGGAGTCCACGTTCTTTAATAGTGGACTCTTGTTCCAAACTGGAACAACACTCAACCCTATCTCGGGCTATTCTTTTGATTTATAAGGGATTTTGCCGATTTCGGAACCACCATCAAACAGGATTTTCGCCTGCTGGGGCAAACCAGCGTGGACCGCTTGCTGCAACTCTCTCAGGGCCAGGCGGTGAAGGGCAATCAGCTGTTGCCCGTCTCACTGGTGAAAAGAAAAACCACCCTGGCGCCCAATACGCAAACCGCCTCTCCCCGCGCGTTGGCCGATTCATTAATGCAGCTGGCACGACAGGTTTCCCGACTGGAAAGCGGGCAGTGAGCGCAACGCAATTAATGTGAGTTAGCTCACTCATTAGGCACCCCAGGCTTTACACTTTATGCTTCCGGCTCGTATGTTGTGTGGAATTGTGAGCGGATAACAATTTCACACAGGAAACAGCTATGACCATGATTACGAATTCGAGCTCGGTACCCGGGGATCCATTCTCCTGTGACTCGGAAGTGCATTTATCATCTCCATAAAACAAAACCCGCCGTAGCGAGTTCAGATAAAATAAATCCCCGCGAGTGCGAGGATTGTTATGTAATATTGGGTTTAATCATCTATATGTTTTGTACAGAGAGGGCAAGTATCGTTTCCACCGTACTCGTGATAATAATTTTGCACGGTATCAGTCATTTCTCGCACATTGCAGAATGGGGATTTGTCTTCATTAGACTTATAAACCTTCATGGAATATTTGTATGCCGACTCTATATCTATACCTTCATCTACATAAACACCTTCGTGATGTCTGCATGGAGACAAGACACCGGATCTGCACAACATTGATAACGCCCAATCTTTTTGCTCAGACTCTAACTCATTGATACTCATTTATAAACTCCTTGCAATGTATGTCGTTTCAGCTAAACGGTATCAGCAATGTTTATGTAAAGAAACAGTAAGATAATACTCAACCCGATGTTTGAGTACGGTCATCATCTGACACTACAGACTCTGGCATCGCTGTGAAGACGACGCGAAATTCAGCATTTTCACAAGCGTTATCTTTTACAAAACCGATCTCACTCTCCTTTGATGCGAATGCCAGCGTCAGACATCATATGCAGATACTCACCTGCATCCTGAACCCATTGACCTCCAACCCCGTAATAGCGATGCGTAATGATGTCGATAGTTACTAACGGGTCTTGTTCGATTAACTGCCGCAGAAACTCTTCCAGGTCACCAGTGCAGTGCTTGATAACAGGAGTCTTCCCAGGATGGCGAACAACAAGAAACTGGTTTCCGTCTTCACGGACTTCGTTGCTTTCCAGTTTAGCAATACGCTTACTCCCATCCGAGATAACACCTTCGTAATACTCACGCTGCTCGTTGAGTTTTGATTTTGCTGTTTCAAGCTCAACACGCAGTTTCCCTACTGTTAGCGCAATATCCTCGTTCTCCTGGTCGCGGCGTTTGATGTATTGCTGGTTTCTTTCCCGTTCATCCAGCAGTTCCAGCACAATCGATGGTGTTACCAATTCATGGAAAAGGTCTGCGTCAAATCCCCAGTCGTCATGCATTGCCTGCTCTGCCGCTTCACGCAGTGCCTGAGAGTTAATTTCGCTCACTTCGAACCTCTCTGTTTACTGATAAGTTCCAGATCCTCCTGGCAACTTGCACAAGTCCGACAACCCTGAACGACCAGGCGTCTTCGTTCATCTATCGGATCGCCACACTCACAACAATGAGTGGCAGATATAGCCTGGTGGTTCAGGCGGCGCATTTTTATTGCTGTGTTGCGCTGTAATTCTTCTATTTCTGATGCTGAATCAATGATGTCTGCCATCTTTCATTAATCCCTGAACTGTTGGTTAATACGCATGAGGGTGAATGCGAATAATAAAGCTTGGCACTGGCCGTCGTTTTACAACGTCGTGACTGGGAAAACCCTGGCGTTACCCAACTTAATCGCCTTGCAGCACATCCCCCTTTCGCCAGCTGGCGTAATAGCGAAGAGGCCCGCACCGATCGCCCTTCCCAACAGTTGCGCAGCCTGAATGGCGAATGGCGCTTTGCCTGGTTTCCGGCACCAGAAGCGGTGCCGGAAAGCTGGCTGGAGTGCGATCTTCCTGAGGCCGATACTGTCGTCGTCCCCTCAAACTGGCAGATGCACGGTTACGATGCGCCCATCTACACCAACGTGACCTATCCCATTACGGTCAATCCGCCGTTTGTTCCCACGGAGAATCCGACGGGTTGTTACTCGCTCACATTTAATGTTGATGAAAGCTGGCTACAGGAAGGCCAGACGCGAATTATTTTTGATGGCGTTCCTATTGGTTAAAAAATGAGCTGATTTAACAAAAATTTAATGCGAATTTTAACAAAATATTAACGTTTACAATTTAAATATTTGCTTATACAATCTTCCTGTTTTTGGGGCTTTTCTGATTATCAACCGGGGTACATATGATTGACATGCTAGTTTTACGATTACCGTTCATCGATTCTCTTGTTTGCTCCAGACTCTCAGGCAATGACCTGATAGCCTTTGTAGATCTCTCAAAAATAGCTACCCTCTCCGGCATTAATTTATCAGCTAGAACGGTTGAATATCATATTGATGGTGATTTGACTGTCTCCGGCCTTTCTCACCCTTTTGAATCTTTACCTACACATTACTCAGGCATTGCATTTAAAATATATGAGGGTTCTAAAAATTTTTATCCTTGCGTTGAAATAAAGGCTTCTCCCGCAAAAGTATTACAGGGTCATAATGTTTTTGGTACAACCGATTTAGCTTTATGCTCTGAGGCTTTATTGCTTAATTTTGCTAATTCTTTGCCTTGCCTGTATGATTTATTGGATGTT

**Table S1. Sequence of P8634.**

| **Basic staple strands** | **Length and positions** |
| --- | --- |
| GGACTCCACAGAGGTGGTCAGATGATGACCGTACTCAAAC | 40mer, [0, 63] start, [26, 40] end |
| TTTTTTTTTTGGGCGATGGCCCCATAAACATTGCTTTTTTTTTT | 44mer, [1, 17] start, [24, 17] end |
| AGGTGCCGTAAAGCACCGAAATCGTTGCAAGGAGCAGAAG | 40mer, [3, 48] start, [24, 56] end |
| TTTTTTTTTTCGTGGCGAGAAATTGTCTCCATGCTTTTTTTTTT | 44mer, [5, 17] start, [20, 17] end |
| AGAAACGCCAGGGTGGCCCCCGATACAGAGATAGTTAGAGTGGCAAAT | 48mer, [5, 48] start, [31, 63] end |
| GCAAGTGTAGCGGTCATGCGTATTAAGGTATAGGACATTCATCCTGGG | 48mer, [7, 48] start, [33, 63] end |
| TTTTTTTTTTTGGTTGCTTTGAGAAATGACTGATTTTTTTTTTT | 44mer, [9, 17] start, [16, 17] end |
| TATAAAAAGTGTAAAGCGCCGCGCATCCAGAAATTCCATG | 40mer, [9, 48] start, [18, 56] end |
| GGAACGGTACGCCGCAGGAGCTAAACAGGAGGGCCGGAAGGTGGAAAC | 48mer, [12, 63] start, [17, 63] end |
| TTTTTTTTTTTCTGAACTCGCTACGGCGGGTTTTGTTTTATGGAG | 45mer, [13, 17] start, [13, 63] end |
| CTCAGAATCCTGAGAAAACATATAATTATCAGTAACAGTATCAGGGTT | 48mer, [15, 48] start, [45, 63] end |
| AACCCAATTCAGAGCGCTCGCGGGGATTTATTTTATTTTTTTTTT | 45mer, [14, 47] start, [12, 17] end |
| TTTTTTTTTTACCGTGCAAAATAGAAACCACCAGTTTTTTTTTT | 44mer, [17, 17] start, [38, 17] end |
| GCCTTGCTCTGCGTGTTGCCCGAAGATTCGCC | 32mer, [16, 63] start, [47, 63] end |
| TAAGTCTAGGGCGCTGCCGCTACAGGGCGCGTACTATTTTTTTTTT | 46mer, [18, 47] start, [8, 17] end |
| TTTTTTTTTTAGACATCACGAATAGATAATACATTTTTTTTTTT | 44mer, [21, 17] start, [34, 17] end |
| AAAAGATTTGGGGTCGTGACGGGGAAAGCCGGCGAATTTTTTTTTT | 46mer, [22, 47] start, [4, 17] end |
| TTTTTTTTTTTGATACCGTTTAGAACCTCAAATATTTTTTTTTT | 44mer, [25, 17] start, [30, 17] end |
| TTTCTTTAACTACGTGAACCAAAATCCCTTAT | 32mer, [24, 47] start, [1, 63] end |
| TATCTAAAACGTCAAAGGGCGAAAAACCGTCTATCATTTTTTTTTT | 46mer, [27, 48] start, [0, 17] end |
| TGAGTATCAACAGTGCCAAAATCAACTGTTGG | 32mer, [29, 56] start, [54, 56] end |
| GTTCAGGAGGTTTTCCAATAGTGACCGTGCAT | 32mer, [30, 63] start, [57, 63] end |
| TTTTTTTTTTTCAAACCCTCAACGCATTCACCCTTTTTTTTTTT | 44mer, [31, 17] start, [50, 17] end |
| TGGTCAGTTCTGAGCAGACATACAGCATCACCCAAATCAA | 40mer, [31, 48] start, [2, 40] end |
| AAGGTTATTTAATGGACGGCCAGTAGTTAATTTGGGCGCA | 40mer, [32, 63] start, [58, 56] end |
| ATAGAGCCTGGCAGATCCGGTGTCGGAAGGGA | 32mer, [33, 48] start, [5, 47] end |
| TTTTTTTTTTTTGAGGATTTAGCCACCAGGCTATTTTTTTTTTT | 44mer, [35, 17] start, [46, 17] end |
| GACTTTACAAGGTTTATGTAGATGGGGGCGAAAGGAGCGG | 40mer, [35, 48] start, [6, 40] end |
| AAACAATTTCTTGTTGATCAGAAATTGAATTAAAAAATAATGGTTTGA | 48mer, [35, 56] start, [60, 56] end |
| TATCAAAAGGTCTGCAATGTGCGACGAGCACG | 32mer, [37, 48] start, [9, 47] end |
| TTTTTTTTTTAAGGAGCGGAATACAGAGAGGTTCTTTTTTTTTT | 44mer, [39, 17] start, [42, 17] end |
| TATTCCTGGATGATTACGAGTACGCATCGTGCTTTCCTCG | 40mer, [39, 48] start, [10, 40] end |
| CAGAGCAGGCAATGCATGACGACTATCCTGATTGTTTGGC | 40mer, [41, 40] start, [40, 48] end |
| TTTTTTTTTTGAAGTGAGCGAAAAGGGTGAGAAATTTTTTTTTT | 44mer, [43, 17] start, [68, 17] end |
| TCAGGATTATACTTCTGGAGGATCTAAAGTACCTAGCATGATTTCAAC | 48mer, [43, 48] start, [73, 63] end |
| CTGGTCGTGGGTTTTGCGGAACAATATTATCA | 32mer, [45, 48] start, [17, 47] end |
| GAGAAACAATATGTACACGCTCAATTAAACCA | 32mer, [44, 63] start, [75, 63] end |
| TTTTTTTTTTATCTGCCACTCAAGAATCGCCATATTTTTTTTTT | 44mer, [47, 17] start, [64, 17] end |
| CGCCTGAAAAGTATTAATTTTAAAACAAATCCCCATTAATTTAATGCG | 48mer, [46, 47] start, [8, 48] end |
| AGCAATAAGTTATACAATAGGAACTCCAATAA | 32mer, [46, 63] start, [77, 63] end |
| TTTTTTTTTTCATGCGTATTAATAAATAAGGCGTTTTTTTTTTT | 44mer, [51, 17] start, [60, 17] end |
| TTATTATTTCAATATCTCTTTAGGTATCAATGTTGTGCCATTAGAGCT | 48mer, [50, 47] start, [4, 48] end |
| CTGCAAGGTGAAAAATCTAAAAGGATATACTG | 32mer, [52, 47] start, [24, 48] end |
| TTTTTTTTTTTATTACGCCAGCACTATATGTAAATTTTTTTTTT | 44mer, [55, 17] start, [56, 17] end |
| GGGGGTATCACGCTGAGAGCCAGCTTGAGTAT | 32mer, [55, 48] start, [27, 47] end |
| GAAGGGCGATCGGTGCGGGCCTCTTCGCTTTTTTTTTT | 38mer, [54, 55] start, [54, 17] end |
| CTCCGGCTAATCCAAAATATAATGCTGTAGCTCAACATGT | 40mer, [56, 63] start, [82, 40] end |
| TTTTTTTTTTTGCTGATGCAAAAGTTTCATTCCATTTTTTTTTT | 44mer, [57, 17] start, [80, 17] end |
| TATATTTTGCCAAGCTAACGCCAGTGCGCATCACCTTGCTGCTGAAAC | 48mer, [59, 48] start, [25, 47] end |
| TTTTTTTTTTTAAATAAGAATAAAAGGTGGCATCTTTTTTTTTT | 44mer, [61, 17] start, [76, 17] end |
| AATCAATCCCTGATTAATGAAAGAAACAACTA | 32mer, [61, 48] start, [33, 47] end |
| TCATATGCAAATGCGCGATTCAGCTTCTTAGAGCCGTCAAGGTGTTTA | 48mer, [63, 48] start, [21, 47] end |
| AGGCATTTGAAGACGCATCAGTAATATCATCAACTGCGTGTAACAATC | 48mer, [66, 47] start, [15, 47] end |
| TTTTTTTTTTTTTAACAACGCCAACATTATGACCTTTTTTTTTT | 44mer, [65, 17] start, [72, 17] end |
| ATTTAATCATAGGCGATCCGATAGAGTAACAT | 32mer, [65, 48] start, [37, 47] end |
| TTTTTTTTTTGGCCGGAGACAGTCAAATCACCATCAAT | 38mer, [69, 17] start, [69, 55] end |
| ATGATATTGTCCAGACGACGATAAAGAGAATATGGAACTT | 40mer, [69, 56] start, [42, 48] end |
| AGGCAATAAACAACATCCCTCATAAAGGTGGCGAGGCATACTTGCCCT | 48mer, [71, 48] start, [101, 63] end |
| ATGCAATGCAGTAATAAGATTCAAATTAACTC | 32mer, [70, 47] start, [43, 47] end |
| TTTTTTTTTTCTGTAATACTTTATTACGCAGTATTTTTTTTTTT | 44mer, [73, 17] start, [94, 17] end |
| TCCAAGAAGCAACACTGAGGAAACGGGAATTAATAGGCTGGAACCTAT | 48mer, [72, 63] start, [119, 63] end |
| ATAAAGCCGTTTAGTACTTAATTGTTGTTGTGAGTGTACGCGTTATTA | 48mer, [74, 47] start, [36, 48] end |
| ATTAAGCACATTAACAGCCTAATTACTAGAAAACATCATT | 40mer, [74, 55] start, [49, 47] end |
| TTAGCAAAATAGCCGAGTCATAAAGGACAGAT | 32mer, [74, 63] start, [105, 63] end |
| TTTTTTTTTTAATTCTACTAATTAGCTATCTTACTTTTTTTTTT | 44mer, [77, 17] start, [90, 17] end |
| AGCGAACCTTGAATCCACCCACAAGGCATTTT | 32mer, [76, 63] start, [107, 63] end |
| CTTGCGGGTTTCGCAAGGGTAATTGACTTCAAACAACGGA | 40mer, [79, 56] start, [109, 63] end |
| TAACCTGTTTTTCAAACCGTGTGACCAACAGTTCAGGTTTCTAAAATA | 48mer, [78, 47] start, [32, 48] end |
| ATGGTCAATCTGCGAATCGCAAAGAACGCGAGAGTTGGGT | 40mer, [78, 55] start, [53, 47] end |
| TTTTTTTTTTTATAACAGTTGATAGACGGGAGAATTTTTTTTTT | 44mer, [81, 17] start, [86, 17] end |
| CGAGTAGAATATTATTCGTTTTAAAAAAATGACCAGAACCTACGAAGG | 48mer, [81, 56] start, [110, 56] end |
| TGTCTGGATCCAATCGCAAGATAAATTATGTG | 32mer, [80, 47] start, [52, 48] end |
| TAAAGCCCTAGGTTGGGTTATATATGGCGAAA | 32mer, [83, 48] start, [55, 47] end |
| TTTTTTTTTTTTAACTGAACACTCCATGTTACTTTTTTTTTTTT | 44mer, [87, 17] start, [106, 17] end |
| TTAAGCCCATATTTTCATTTGCGAAATACCGA | 32mer, [88, 47] start, [60, 48] end |
| TTTTTTTTTTCGAAGCCCTTTTAGTAATCTTGACTTTTTTTTTT | 44mer, [91, 17] start, [102, 17] end |
| AACGGAATCATAAAGCTAAATGTACAGTAGGG | 32mer, [92, 47] start, [64, 48] end |
| TTTTTTTTTTGTTAGCAAACGTAATTTCAACTTTTTTTTTTTTT | 44mer, [95, 17] start, [98, 17] end |
| GAATAGATCATACATATATTTTAAAAGCCTTTTCAGGCAG | 40mer, [96, 55] start, [66, 48] end |
| CATTATACCAGTCAGGACGTTGGGACACCACG | 32mer, [97, 40] start, [96, 56] end |
| TTTTTTTTTTAATCATTGTGAAACCGCCACCCTCTTTTTTTTTT | 44mer, [99, 17] start, [124, 17] end |
| ATTTTGTCAGTAAATTGCCCGGAAGTCTTTCCAGAGGATA | 40mer, [99, 56] start, [122, 48] end |
| GAATAAGGGTAATTAAGACTCCTTTGCGGGAG | 32mer, [101, 48] start, [73, 47] end |
| GTAACAAAAAGAACTGGCATGAGACGGCGGTTGTACCAAAAACATGTA | 48mer, [100, 47] start, [65, 47] end |
| GACTTGAGTAGTAAATAAGAGGCTGAGACTCCCTATTTCGGCTGACCT | 48mer, [100, 63] start, [102, 48] end |
| TTTTTTTTTTAAGAACCGGATAAGGATTAGGATTTTTTTTTTTT | 44mer, [103, 17] start, [120, 17] end |
| TTGAAAGATATAATGAAATAGCAAAGTAGTAG | 32mer, [105, 48] start, [77, 47] end |
| AAGGGAACAAGAGCAAGAAACTCATCCGGGCGCGAGCTGAAACACCGG | 48mer, [104, 47] start, [61, 47] end |
| AGCGCGTTTAATTGTATGATGATACAGGAGTGCTGAATTTGAAATCCG | 48mer, [104, 63] start, [106, 48] end |
| TTTTTTTTTTAGCCGGAACGAGATAAGTTTTAACTTTTTTTTTT | 44mer, [107, 17] start, [116, 17] end |
| ATTGTGTCACCGTTCCTTAAAGGCCGCTTAAA | 32mer, [106, 63] start, [114, 48] end |
| AACAAAGTATACAGGGAAGCGCATTTCCCAAT | 32mer, [109, 48] start, [81, 47] end |
| CCAGCGATAATAACATAAAAATCGTATTACGG | 32mer, [108, 47] start, [80, 48] end |
| CCTCAGAGTGCGGGATCAGGAGGTTGAGGCAG | 32mer, [108, 63] start, [112, 48] end |
| TTTTTTTTTTGAATACACTAAATTGGCCTTGATTTTTTTTTT | 42mer, [111, 17] start, [112, 17] end |
| CACCAACCTAAAACGAAAGAGGCAAAATTTTTTTTTT | 37mer, [110, 55] start, [110, 17] end |
| GTCAGACGAACACTCATCTTTCCAAAATAGCAGCCTTTACTATGCAAC | 48mer, [112, 47] start, [83, 47] end |
| TTTTTTTTTTTATTCACAAACAAATAAATCCTCATTTCCGGACCC | 45mer, [113, 17] start, [108, 48] end |
| GCCAGAATAAGCGCGACGACCTGCCCTGAACA | 32mer, [114, 47] start, [87, 47] end |
| CGCAGTCTTACTGGTAGCGCAGACGGTCAATCGAATTGAGAAGTCAGA | 48mer, [115, 40] start, [87, 55] end |
| TTTTTTTTTTGGGGTCAGTGCCTTGAGTAACAGTGCTTTTCATCAT | 46mer, [117, 17] start, [104, 48] end |
| CCAAAAGGAGCCCCGTATAAACAGGACCAACTTCATCAAGTAAGAAAA | 48mer, [118, 63] start, [91, 47] end |
| CCCCCTGCTCAAGAGATTCATTACCCAAATTTGCAATAATGTAAGCAG | 48mer, [119, 40] start, [91, 55] end |
| TTTTTTTTTTAGCGGGGTTTTGCTCAGTACCAGGCCGTCCATCAAC | 46mer, [121, 17] start, [100, 48] end |
| AGTGCCGTCATTCAGTGATGGTTTAGAAAATATTTAAGAATAATGTGT | 48mer, [122, 47] start, [71, 47] end |
| GGTTGATACCCTCAGATTACCTTATGCAGTTT | 32mer, [123, 40] start, [99, 55] end |
| TTTTTTTTTTAGAACCGCCACCCTCAGAGCCACCACCC | 38mer, [125, 17] start, [125, 55] end |
| TCATTTTCAGGTTTAGTACCGCCATAAGTATAGGGCTTGA | 40mer, [125, 56] start, [98, 48] end |
| TAAATCGGTATCAATGAGAACCCTAATTGAGGAAGACTCCAATTTCAT | 48mer, [3, 64] start, [48, 64] end |
| AAAGGGAGTTTTTCTTAATAAAAGGATATAGAAACCAGTT | 40mer, [4, 71] start, [34, 64] end |
| ACCACACCCCTGGGGTAACTATCGGATACTTGTATTGCGC | 40mer, [8, 71] start, [38, 64] end |
| CCGATTAACTGTACAAGTGTTTTTTCAATATAGGGGATTT | 40mer, [11, 64] start, [41, 71] end |
| TGAGGCCACTCGAATTCGTAAGATTTTAGACAATGATAAA | 40mer, [15, 80] start, [13, 71] end |
| AGAACTCAGCCTAATGTCCACACAACATACGA | 32mer, [16, 79] start, [10, 64] end |
| CAATATTAAAATCCTTTGAGCTTGTACATCGGGTCGGACTTCGTAAAA | 48mer, [19, 64] start, [66, 64] end |
| CGACCAGTTTCACCAGGCGGGGAGAGGCGGTTCGCTGCGCATACAAAT | 48mer, [20, 79] start, [18, 64] end |
| AGTTTATAGTCAATGGCAACAGTTTAAAACGAAACAGTACAAATTTAA | 48mer, [25, 64] start, [60, 64] end |
| AGGCGGTCAGTATATATACCGAACAGAATAGCTGTTTGATGGTGGTTC | 48mer, [27, 64] start, [2, 64] end |
| ACGCTTGTCAAGAGTCCACTATTAAAGAACGTAAATCAAAGAACCACC | 48mer, [27, 80] start, [24, 64] end |
| AAGTAACACCGCCTGCTGCATATGGAAGAGTC | 32mer, [28, 79] start, [52, 64] end |
| AGCACTGCCAGTCACAGAAAGCGTTGGCCCTGAGAGACCT | 40mer, [33, 80] start, [4, 72] end |
| CGACAACTCGCAACACTGATTGCTTAAAGCCA | 32mer, [35, 64] start, [64, 64] end |
| AAATCAAATGAGTAGACATTGCAACTGCCCGCTTTCCACC | 40mer, [37, 80] start, [8, 72] end |
| ATGATGGCAGTTGCCAGAATAATGGTAATTCTCAACCGTT | 40mer, [39, 64] start, [69, 71] end |
| AGAACCTACTGGAACTGCTGGTCAATAATCAG | 32mer, [43, 80] start, [15, 79] end |
| AAAGAGCAAATATGAACGGGAAAGCAAATTAA | 32mer, [42, 79] start, [14, 80] end |
| ACCTGCAGCGTGGATGGGAGTAAGACATTTTG | 32mer, [46, 79] start, [18, 80] end |
| TAGAAGAATTTAACCAAATTCTTAGCAAAGAAAGTACCGCAGGAAACC | 48mer, [49, 64] start, [92, 64] end |
| TTAATTTGGAACAAGACCCGTTAGTTGAATGG | 32mer, [50, 79] start, [22, 80] end |
| CAGTCACGGGTGTAGATCATCTTCTAGATACAAGGTTTTGGAGAGATA | 48mer, [53, 64] start, [88, 64] end |
| ACGTAATTTTCCCTTATATTACGGCGCTGGCAATTAAAAA | 40mer, [53, 72] start, [24, 80] end |
| TAGGTCTGAGAGATCCGACGCTGAATGTCTGAGGTTGGAGAATAGTCT | 48mer, [55, 64] start, [25, 79] end |
| CTTCTACCTTTTTAACCTGCCAGTAAACAGCC | 32mer, [56, 79] start, [80, 64] end |
| ACGACGACTAGATTAAGGCACCGCTTTTGTAA | 32mer, [57, 80] start, [28, 80] end |
| ACCCGCCTATAAATCACATTTAACTGTTATCAAAGACGGAGTCCAATC | 48mer, [60, 79] start, [21, 79] end |
| TTCGCGTCGGCGTTTTATCATACAGCGGAATCACAAAGTTCCAGGCGC | 48mer, [61, 64] start, [102, 64] end |
| CTGTAGCCATTAATTAATATATGTGCAGTTAATCGAAAGGTCTGACCT | 48mer, [61, 80] start, [23, 79] end |
| ATTGTGTATTGAATACGTACCTTTAAACAGCAAACGAGGACCCGTAGC | 48mer, [64, 79] start, [17, 79] end |
| CCCGGTTGCTTATCATGCAAGGATGGAATTAC | 32mer, [65, 64] start, [94, 64] end |
| AAAAGCCCCAGTAACACAAGTTACAAGGTGTTATCTCATTCCGCCAGC | 48mer, [65, 80] start, [19, 79] end |
| CGACAAAATTTTAGAAGTTCAGCTACGCAAAGAAGAAAAA | 40mer, [67, 64] start, [97, 71] end |
| ACGCGCCTAGATCTACAAAGGAAAGAAGGGTT | 32mer, [71, 80] start, [43, 79] end |
| AATAAAATGGTCTATCAGGTCATTACAGAAAT | 32mer, [70, 79] start, [42, 80] end |
| TGTCTTTCATAATCAGAACGGTAATGTAATTGCGTAGATTACCAGGAG | 48mer, [72, 79] start, [38, 80] end |
| GAACGCGATGGCCTTCGCTCATTTTTAAGCAAAAGAAGATAGTCCGTG | 48mer, [76, 79] start, [34, 80] end |
| CCTGACATTGATCGGATTCTCCGTCGTCGCTA | 32mer, [78, 79] start, [50, 80] end |
| TTTAGTTTGCCCGAAAGAGCGCTACCTGATAACGGTCATAATGGCTTT | 48mer, [81, 64] start, [116, 64] end |
| ATTGCTGATAAGAAACGATTTCTTTACAAAATTTGAGGGGGTCACGTT | 48mer, [82, 71] start, [58, 72] end |
| TTAACGTCTTCGAGCTGCCACCACGATTTGTAGCAGGGAG | 40mer, [84, 71] start, [114, 64] end |
| AAGAGGAAGACATCTTACCAACGCTGGGATAG | 32mer, [86, 79] start, [58, 80] end |
| CCCTCAAACAGACTGTGAACGGTGAAAAGGCTTATTCTGAAACGGAAC | 48mer, [89, 64] start, [119, 79] end |
| TGCCAGAGAATCATTACCGCGAAGCCAAAACGTTAATATTTTTCAATT | 48mer, [91, 80] start, [46, 80] end |
| ATCATAACCCGTCACCGACGAGAAGTTTTGTCTAGGTGTATCATACAA | 48mer, [93, 64] start, [123, 79] end |
| AACATATAGAACGAGTACAATCAATACTCAGGAGGGATAG | 40mer, [95, 64] start, [125, 71] end |
| TCATATGGGTAGAAAGATTCAGAAAATGCAGA | 32mer, [99, 80] start, [71, 79] end |
| GGAGGGAAATACATAAACCAGACGTAATCGGCAGAACAAGATATTTAA | 48mer, [101, 80] start, [64, 80] end |
| AATTATCACCTCGTTTCGCCAAAAAAATATCCCATCCTAAAATCGATG | 48mer, [100, 79] start, [66, 80] end |
| TAAAGGTGCAGTAGCACTTTTGCAAAAGAAGAACTCATCG | 40mer, [100, 87] start, [75, 79] end |
| GAGCCAGCAAAGTATTGAATTTTCTGTATGGGGATCTAAAACATTGAG | 48mer, [103, 64] start, [101, 79] end |
| GATAGCAGGATAGCGTACAGTTCATATTCTAAAATCAAGAGAGTAACA | 48mer, [105, 80] start, [60, 80] end |
| TTTAGCGTTGCTTTAACCAATACTGGCCCCAATAGCAAGCTTAAATCA | 48mer, [104, 79] start, [62, 80] end |
| AGTTTGCCATTAGCGTAATCAGGTCTTTATCAAAGCCTTA | 40mer, [104, 87] start, [79, 79] end |
| AGAGCTCGATACCCTGACTATTATAATTTTAT | 32mer, [106, 79] start, [78, 80] end |
| CCGGAACCAAAAGATTAACCAGACTTGCCAGTTAATTGCTCCAGCCAG | 48mer, [109, 80] start, [56, 80] end |
| AACCGCCAGACTAAAGTTAGAGAGTACTTTGT | 32mer, [108, 87] start, [84, 72] end |
| AATGCCACACCACCAGAGCCGGAGCCCTCAGATCAAAGCG | 40mer, [110, 71] start, [85, 79] end |
| AGCATTGACGTCACCCTCAGCAGCTGAGGCTTTCACACCA | 40mer, [112, 71] start, [109, 79] end |
| AGTAAGCGTCAACAATGACAACAACCGGAACC | 32mer, [115, 64] start, [106, 80] end |
| TCGGTTTATCAGCTTGCTCCAAAATACCCATC | 32mer, [117, 64] start, [105, 79] end |
| AACTAAAGGTCACCAATGAAAAGAACCAGTTT | 32mer, [119, 80] start, [91, 79] end |
| CGCCTGTAGACATTCAACCGACCAAAATCAGTTGAGATTTACAAGAAA | 48mer, [123, 80] start, [70, 80] end |
| TTTTTTTTTTTTCCAGTTTGGAAGAAAATGCTGAATTTTTTTTTTT | 46mer, [0, 110] start, [27, 110] end |
| TTTTTTTTTTTGCCCCAGCAGGCCGAACTGATAGCCTTTTTTTTTT | 46mer, [2, 110] start, [25, 110] end |
| TTTTTTTTTTGCCCTTCACCGCCAAGAATACGTGGCTTTTTTTTTT | 46mer, [4, 110] start, [23, 110] end |
| TTTTTTTTTTCATTAATGAATCGTGGATTATTTACATTTTTTTTTT | 46mer, [6, 110] start, [21, 110] end |
| TTTTTTTTTTTGCGTTGCGCTCACAGGAAAAACGCTTTTTTTTTTT | 46mer, [8, 110] start, [19, 110] end |
| TTTTTTTTTTAAATTGTTATCCGTTTGATTAGTAATTTTTTTTTTT | 46mer, [10, 110] start, [17, 110] end |
| TTTTTTTTTTCCCGGGTACCGAGCCGAGTAAAAGAGTTTTTTTTTT | 46mer, [12, 110] start, [15, 110] end |
| TGCACTTCCGAGTCACAGGAGAATGGATCTTTTTTTTTT | 39mer, [13, 72] start, [13, 110] end |
| CCGTTTCTAGGTCATGGTCATAGCTGTTTCCTGTGTGTTTTTTTTTT | 47mer, [14, 79] start, [11, 110] end |
| TTTTTTTTTTTCTGTCCATCACGAAACCAGCAATACTTTTTTTTTT | 46mer, [14, 110] start, [39, 110] end |
| AATACTTCCTCACAATAGTGAGCTAACTCACATTAATTTTTTTTTTT | 47mer, [17, 80] start, [9, 110] end |
| TTTTTTTTTTAACATCACTTGCCACTCAACGAGCAGTTTTTTTTTT | 46mer, [16, 110] start, [37, 110] end |
| ACGCTGGCGTAAGTCGGGAAACCTGTCGTGCCAGCTGTTTTTTTTTT | 47mer, [18, 79] start, [7, 110] end |
| TTTTTTTTTTCATGGAAATACCTCGTATTGCTAAACTTTTTTTTTT | 46mer, [18, 110] start, [35, 110] end |
| GTCTGAAAGCCAACGCTGAGACGGGCAACAGCTGATTTTTTTTTTTT | 47mer, [21, 80] start, [5, 110] end |
| TTTTTTTTTTTTGGCAGATTCACACTGGTGACCTGGTTTTTTTTTT | 46mer, [20, 110] start, [33, 110] end |
| CTATTGAGAACGTTGCAGCAAGCGGTCCACGCTGGTTTTTTTTTTTT | 47mer, [22, 79] start, [3, 110] end |
| TTTTTTTTTTACAGACAATATTTTAACTATCGACATTTTTTTTTTT | 46mer, [22, 110] start, [31, 110] end |
| TTAATGCGGAAAATCCCCGAGATAGGGTTGAGTGTTGTTTTTTTTTT | 47mer, [25, 80] start, [1, 110] end |
| TTTTTTTTTTCTAAAACATCGCCTTCGCATCAAAGGTTTTTTTTTT | 46mer, [24, 110] start, [29, 110] end |
| TTTTTTTTTTTTCGCGTCGTCTTCACAGCGATGCCAGAGTCTGTAGT | 47mer, [26, 110] start, [26, 64] end |
| TTTTTTTTTTAGAGTGAGATCGGTTCTGGTGCCGGATTTTTTTTTT | 46mer, [28, 110] start, [55, 110] end |
| TTTTTTTTTTCATTACGCATCGCGAATCCTTGAAAATTTTTTTTTT | 46mer, [30, 110] start, [53, 110] end |
| TTTTTTTTTTAAGAGTTTCTGCGGAGTGAATAACCTTTTTTTTTTT | 46mer, [32, 110] start, [51, 110] end |
| TTTTTTTTTTTGGAAAGCAACGAGATGAAACAAACATTTTTTTTTT | 46mer, [34, 110] start, [49, 110] end |
| TTTTTTTTTTCGTGAGTATTACGAAAATCGCGCAGATTTTTTTTTT | 46mer, [36, 110] start, [47, 110] end |
| TTTTTTTTTTATCAAACGCCGCGTTCAGGTTTAACGTTTTTTTTTT | 46mer, [38, 110] start, [45, 110] end |
| TTTTTTTTTTACCATCGATTGTGCCATATCAAAATTTTTTTTTTTT | 46mer, [40, 110] start, [43, 110] end |
| GACGCAGACCTTTTCCATGAATTGGTAACTTTTTTTTTT | 39mer, [41, 72] start, [41, 110] end |
| TTTTTTTTTTATTTGCACGTAAAGCCTGAGAGTCTGTTTTTTTTTT | 46mer, [42, 110] start, [67, 110] end |
| TTTTTTTTTTTCAGATGAATATACAAAAACAGGAAGTTTTTTTTTT | 46mer, [44, 110] start, [65, 110] end |
| TTTTTTTTTTGGCGAATTATTCATTGTTAAAATTCGTTTTTTTTTT | 46mer, [46, 110] start, [63, 110] end |
| TTTTTTTTTTTCAAGAAAACAAAAGCTTTCATCAACTTTTTTTTTT | 46mer, [48, 110] start, [61, 110] end |
| TTTTTTTTTTTGCTTCTGTAAATGGGAACAAACGGCTTTTTTTTTT | 46mer, [50, 110] start, [59, 110] end |
| TTTTTTTTTTCATAGCGATAGCTAGTATCGGCCTCATTTTTTTTTT | 46mer, [52, 110] start, [57, 110] end |
| TTTTTTTTTTAACCAGGCAAAGCGCCATTCGCCATTCAGGCTGCGCA | 47mer, [54, 110] start, [54, 64] end |
| TTTTTTTTTTGGAAGATCGCACTCCTTTTGATAAGATTTTTTTTTT | 46mer, [56, 110] start, [83, 110] end |
| TTTTTTTTTTGGATTGACCGTAATAACGAGCGTCTTTTTTTTTTTT | 46mer, [58, 110] start, [81, 110] end |
| TTTTTTTTTTATTAAATGTGAGCTTAGTTGCTATTTTTTTTTTTTT | 46mer, [60, 110] start, [79, 110] end |
| TTTTTTTTTTCATTAAATTTTTGAAATCAGATATAGTTTTTTTTTT | 46mer, [62, 110] start, [77, 110] end |
| TTTTTTTTTTATTGTATAAGCAACAAGCCGTTTTTATTTTTTTTTT | 46mer, [64, 110] start, [75, 110] end |
| TTTTTTTTTTGAGCAAACAAGAGTTTACGAGCATGTTTTTTTTTTT | 46mer, [66, 110] start, [73, 110] end |
| TTTTTTTTTTAGCTATTTTTGAGGTTTATCAACAATTTTTTTTTTT | 46mer, [68, 110] start, [71, 110] end |
| CTAGCTGATAAATTAATGCCGGAGAGGGTTTTTTTTTTT | 39mer, [69, 72] start, [69, 110] end |
| TTTTTTTTTTAGATAAGTCCTGAAGGAATACCACATTTTTTTTTTT | 46mer, [70, 110] start, [95, 110] end |
| TTTTTTTTTTAGAAACCAATCAAACGATAAAAACCATTTTTTTTTT | 46mer, [72, 110] start, [93, 110] end |
| TTTTTTTTTTTTTTCATCGTAGGGGGGTAATAGTAATTTTTTTTTT | 46mer, [74, 110] start, [91, 110] end |
| TTTTTTTTTTAAGGCTTATCCGGGAAAACGAGAATGTTTTTTTTTT | 46mer, [76, 110] start, [89, 110] end |
| TTTTTTTTTTTGCACCCAGCTACAGTCAGAAGCAAATTTTTTTTTT | 46mer, [78, 110] start, [87, 110] end |
| TTTTTTTTTTTCCAGAGCCTAATCGGAAGCAAACTCTTTTTTTTTT | 46mer, [80, 110] start, [85, 110] end |
| TTTTTTTTTTGGTCATTTTTGCGGATGGCTTAGAGCTTA | 39mer, [82, 110] start, [82, 72] end |
| TTTTTTTTTTCAACAGGTCAGGAACTTTTTCATGAGTTTTTTTTTT | 46mer, [84, 110] start, [111, 110] end |
| TTTTTTTTTTGCGGATTGCATCAGCCTCCCTCAGAGTTTTTTTTTT | 46mer, [86, 110] start, [109, 110] end |
| TTTTTTTTTTACCATAAATCAAATTGCCATCTTTTCTTTTTTTTTT | 46mer, [88, 110] start, [107, 110] end |
| TTTTTTTTTTAATGTTTAGACTGCACCGTAATCAGTTTTTTTTTTT | 46mer, [90, 110] start, [105, 110] end |
| TTTTTTTTTTAAATAGCGAGAGGCCATTACCATTAGTTTTTTTTTT | 46mer, [92, 110] start, [103, 110] end |
| TTTTTTTTTTTCAACTAATGCAGGGTAAATATTGACTTTTTTTTTT | 46mer, [94, 110] start, [101, 110] end |
| TTTTTTTTTTAACATTATTACAGTTTACCAGCGCCATTTTTTTTTT | 46mer, [96, 110] start, [99, 110] end |
| TCTACGTTAATAAAACGAACTAACGGAACTTTTTTTTTT | 39mer, [97, 72] start, [97, 110] end |
| TTTTTTTTTTAAGACAAAAGGGCGCATTCCACAGACTTTTTTTTTT | 46mer, [98, 110] start, [123, 110] end |
| TTTTTTTTTTGGAAATTATTCATAAACAACTTTCAATTTTTTTTTT | 46mer, [100, 110] start, [121, 110] end |
| TTTTTTTTTTCAAGGCCGGAAACGAATTGCGAATAATTTTTTTTTT | 46mer, [102, 110] start, [119, 110] end |
| TTTTTTTTTTAGCGACAGAATCAGTGAATTTCTTAATTTTTTTTTT | 46mer, [104, 110] start, [117, 110] end |
| TTTTTTTTTTATAATCAAAATCACCATCGCCCACGCTTTTTTTTTT | 46mer, [106, 110] start, [115, 110] end |
| TTTTTTTTTTCCGCCACCCTCAGGCATCGGAACGAGTTTTTTTTTT | 46mer, [108, 110] start, [113, 110] end |
| TTTTTTTTTTGAAGTTTCCATTAAACGGGTAAAATACGT | 39mer, [110, 110] start, [110, 72] end |
| TTTTTTTTTTGGTAGCAACGGCTACAGAGGCTTTCCGCC | 39mer, [112, 110] start, [112, 72] end |
| TTTTTTTTTTATAACCGATATATTCGGTCGCGAAAGACA | 39mer, [114, 110] start, [113, 87] end |
| TTTTTTTTTTACAGCTTGATACCGATAGTTGCGCCGTACGCCCCCTT | 47mer, [116, 110] start, [107, 79] end |
| TTTTTTTTTTTAATTTTTTCACGTTGAAAATCTTTCGAG | 39mer, [118, 110] start, [117, 87] end |
| TTTTTTTTTTCAGTTTCAGCGGAGTGAGAATAGAAAATGAAAATCAC | 47mer, [120, 110] start, [103, 79] end |
| TTTTTTTTTTAGCCCTCATAGTTAGCGTAACATTTTGCT | 39mer, [122, 110] start, [121, 87] end |
| TTTTTTTTTTCACTGAGTTTCGTCACCAGTACAAACCCGTAGAAAAT | 47mer, [124, 110] start, [99, 79] end |
| CAAGCCCAATAGGAACCCATGTACCGTAATTTTTTTTTT | 39mer, [125, 72] start, [125, 110] end |

**Table S2. DNA oligonucleotide staple sequences for SQB design corresponding to figure S2.** (Left: purple strands, middle: green strands, right: orange strands)

| **CpG-containing staples for various CpG spacing patterns** | Start | End |
| --- | --- | --- |
| **2.5nm spacing-CpG1-18 strands** |  |  |
| TTTTTTTTTTCATGGAAATACCTCGTATTGCTAAACt*c*c*a*t*g*a*c*g*t*t*c*c*t*g*a*c*g*t*t* | 18[110] | 35[120] |
| TTTTTTTTTTTTGGCAGATTCACACTGGTGACCTGGt*c*c*a*t*g*a*c*g*t*t*c*c*t*g*a*c*g*t*t* | 20[110] | 33[120] |
| t*c*c*a*t*g*a*c*g*t*t*c*c*t*g*a*c*g*t*t*TGGAAAGCAACGAGATGAAACAAACAt*c*c*a*t*g*a*c*g*t*t*c*c*t*g*a*c*g*t*t* | 34[120] | 49[120] |
| t*c*c*a*t*g*a*c*g*t*t*c*c*t*g*a*c*g*t*t*CGTGAGTATTACGAAAATCGCGCAGATTTTTTTTTT | 36[120] | 47[110] |
| t*c*c*a*t*g*a*c*g*t*t*c*c*t*g*a*c*g*t*t*GGCGAATTATTCATTGTTAAAATTCGt*c*c*a*t*g*a*c*g*t*t*c*c*t*g*a*c*g*t*t* | 46[120] | 63[120] |
| t*c*c*a*t*g*a*c*g*t*t*c*c*t*g*a*c*g*t*t*TCAAGAAAACAAAAGCTTTCATCAACt*c*c*a*t*g*a*c*g*t*t*c*c*t*g*a*c*g*t*t* | 48[120] | 61[120] |
| t*c*c*a*t*g*a*c*g*t*t*c*c*t*g*a*c*g*t*t*CATTAAATTTTTGAAATCAGATATAGt*c*c*a*t*g*a*c*g*t*t*c*c*t*g*a*c*g*t*t* | 62[120] | 77[120] |
| t*c*c*a*t*g*a*c*g*t*t*c*c*t*g*a*c*g*t*t*ATTGTATAAGCAACAAGCCGTTTTTATTTTTTTTTT | 64[120] | 75[110] |
| t*c*c*a*t*g*a*c*g*t*t*c*c*t*g*a*c*g*t*t*TTTTCATCGTAGGGGGGTAATAGTAAt*c*c*a*t*g*a*c*g*t*t*c*c*t*g*a*c*g*t*t* | 74[120] | 91[120] |
| t*c*c*a*t*g*a*c*g*t*t*c*c*t*g*a*c*g*t*t*AAGGCTTATCCGGGAAAACGAGAATGt*c*c*a*t*g*a*c*g*t*t*c*c*t*g*a*c*g*t*t* | 76[120] | 89[120] |
| t*c*c*a*t*g*a*c*g*t*t*c*c*t*g*a*c*g*t*t*AATGTTTAGACTGCACCGTAATCAGTTTTTTTTTTT | 90[120] | 105[110] |
| t*c*c*a*t*g*a*c*g*t*t*c*c*t*g*a*c*g*t*t*AAATAGCGAGAGGCCATTACCATTAGTTTTTTTTTT | 92[120] | 103[110] |
| **2.5nm spacing-CpG1-18 strands 3´-3´ linkage** |  |  |
| TTTTTTTTTTCATGGAAATACCTCGTATTGCTAAAC  t*c*c*a*t*g*a*c*g*t*t*c*c*t*g*a*c*g*t*t* | 18[110] | 35[120] |
| TTTTTTTTTTTTGGCAGATTCACACTGGTGACCTGG  t*c*c*a*t*g*a*c*g*t*t*c*c*t*g*a*c*g*t*t* | 20[110] | 33[120] |
| t*c*c*a*t*g*a*c*g*t*t*c*c*t*g*a*c*g*t*t*TGGAAAGCAACGAGATGAAACAAACA  t*c*c*a*t*g*a*c*g*t*t*c*c*t*g*a*c*g*t*t* | 34[120] | 49[120] |
| t*c*c*a*t*g*a*c*g*t*t*c*c*t*g*a*c*g*t*t*CGTGAGTATTACGAAAATCGCGCAGATTTTTTTTTT | 36[120] | 47[110] |
| t*c*c*a*t*g*a*c*g*t*t*c*c*t*g*a*c*g*t*t*GGCGAATTATTCATTGTTAAAATTCG  t*c*c*a*t*g*a*c*g*t*t*c*c*t*g*a*c*g*t*t* | 46[120] | 63[120] |
| t*c*c*a*t*g*a*c*g*t*t*c*c*t*g*a*c*g*t*t*TCAAGAAAACAAAAGCTTTCATCAAC  t*c*c*a*t*g*a*c*g*t*t*c*c*t*g*a*c*g*t*t* | 48[120] | 61[120] |
| t*c*c*a*t*g*a*c*g*t*t*c*c*t*g*a*c*g*t*t*CATTAAATTTTTGAAATCAGATATAG  t*c*c*a*t*g*a*c*g*t*t*c*c*t*g*a*c*g*t*t* | 62[120] | 77[120] |
| t*c*c*a*t*g*a*c*g*t*t*c*c*t*g*a*c*g*t*t*ATTGTATAAGCAACAAGCCGTTTTTATTTTTTTTTT | 64[120] | 75[110] |
| t*c*c*a*t*g*a*c*g*t*t*c*c*t*g*a*c*g*t*t*TTTTCATCGTAGGGGGGTAATAGTAA  t*c*c*a*t*g*a*c*g*t*t*c*c*t*g*a*c*g*t*t* | 74[120] | 91[120] |
| t*c*c*a*t*g*a*c*g*t*t*c*c*t*g*a*c*g*t*t*AAGGCTTATCCGGGAAAACGAGAATG  t*c*c*a*t*g*a*c*g*t*t*c*c*t*g*a*c*g*t*t* | 76[120] | 89[120] |
| t*c*c*a*t*g*a*c*g*t*t*c*c*t*g*a*c*g*t*t*AATGTTTAGACTGCACCGTAATCAGTTTTTTTTTTT | 90[120] | 105[110] |
| t*c*c*a*t*g*a*c*g*t*t*c*c*t*g*a*c*g*t*t*AAATAGCGAGAGGCCATTACCATTAGTTTTTTTTTT | 92[120] | 103[110] |
| **3.5nm spacing-CpG2-18 strands** |  |  |
| t*c*c*a*t*g*a*c*g*t*t*c*c*t*g*a*c*g*t*t*AAGAGTTTCTGCGGAGTGAATAACCTTTTTTTTTTT | 32[120] | 51[110] |
| t*c*c*a*t*g*a*c*g*t*t*c*c*t*g*a*c*g*t*t*TGGAAAGCAACGAGATGAAACAAACATTTTTTTTTT | 34[120] | 49[110] |
| t*c*c*a*t*g*a*c*g*t*t*c*c*t*g*a*c*g*t*t*CGTGAGTATTACGAAAATCGCGCAGATTTTTTTTTT | 36[120] | 47[110] |
| t*c*c*a*t*g*a*c*g*t*t*c*c*t*g*a*c*g*t*t*ATCAAACGCCGCGTTCAGGTTTAACGTTTTTTTTTT | 38[120] | 45[110] |
| t*c*c*a*t*g*a*c*g*t*t*c*c*t*g*a*c*g*t*t*TCAGATGAATATACAAAAACAGGAAGTTTTTTTTTT | 44[120] | 65[110] |
| t*c*c*a*t*g*a*c*g*t*t*c*c*t*g*a*c*g*t*t*GGCGAATTATTCATTGTTAAAATTCGTTTTTTTTTT | 46[120] | 63[110] |
| t*c*c*a*t*g*a*c*g*t*t*c*c*t*g*a*c*g*t*t*TCAAGAAAACAAAAGCTTTCATCAACTTTTTTTTTT | 48[120] | 61[110] |
| t*c*c*a*t*g*a*c*g*t*t*c*c*t*g*a*c*g*t*t*ATTAAATGTGAGCTTAGTTGCTATTTTTTTTTTTTT | 60[120] | 79[110] |
| t*c*c*a*t*g*a*c*g*t*t*c*c*t*g*a*c*g*t*t*CATTAAATTTTTGAAATCAGATATAGTTTTTTTTTT | 62[120] | 77[110] |
| t*c*c*a*t*g*a*c*g*t*t*c*c*t*g*a*c*g*t*t*ATTGTATAAGCAACAAGCCGTTTTTATTTTTTTTTT | 64[120] | 75[110] |
| t*c*c*a*t*g*a*c*g*t*t*c*c*t*g*a*c*g*t*t*GAGCAAACAAGAGTTTACGAGCATGTTTTTTTTTTT | 66[120] | 73[110] |
| t*c*c*a*t*g*a*c*g*t*t*c*c*t*g*a*c*g*t*t*AGAAACCAATCAAACGATAAAAACCATTTTTTTTTT | 72[120] | 93[110] |
| t*c*c*a*t*g*a*c*g*t*t*c*c*t*g*a*c*g*t*t*TTTTCATCGTAGGGGGGTAATAGTAATTTTTTTTTT | 74[120] | 91[110] |
| t*c*c*a*t*g*a*c*g*t*t*c*c*t*g*a*c*g*t*t*AAGGCTTATCCGGGAAAACGAGAATGTTTTTTTTTT | 76[120] | 89[110] |
| t*c*c*a*t*g*a*c*g*t*t*c*c*t*g*a*c*g*t*t*ACCATAAATCAAATTGCCATCTTTTCTTTTTTTTTT | 88[120] | 107[110] |
| t*c*c*a*t*g*a*c*g*t*t*c*c*t*g*a*c*g*t*t*AATGTTTAGACTGCACCGTAATCAGTTTTTTTTTTT | 90[120] | 105[110] |
| t*c*c*a*t*g*a*c*g*t*t*c*c*t*g*a*c*g*t*t*AAATAGCGAGAGGCCATTACCATTAGTTTTTTTTTT | 92[120] | 103[110] |
| t*c*c*a*t*g*a*c*g*t*t*c*c*t*g*a*c*g*t*t*TCAACTAATGCAGGGTAAATATTGACTTTTTTTTTT | 94[120] | 101[110] |
| **5nm spacing-CpG3-18 staples** |  |  |
| t*c*c*a*t*g*a*c*g*t*t*c*c*t*g*a*c*g*t*t*GCCCTTCACCGCCAAGAATACGTGGCTTTTTTTTTT | 4[120] | 23[110] |
| t*c*c*a*t*g*a*c*g*t*t*c*c*t*g*a*c*g*t*t*CATTAATGAATCGTGGATTATTTACATTTTTTTTTT | 6[120] | 21[110] |
| t*c*c*a*t*g*a*c*g*t*t*c*c*t*g*a*c*g*t*t*TGCGTTGCGCTCACAGGAAAAACGCTTTTTTTTTTT | 8[120] | 19[110] |
| t*c*c*a*t*g*a*c*g*t*t*c*c*t*g*a*c*g*t*t*AAATTGTTATCCGTTTGATTAGTAATTTTTTTTTTT | 10[120] | 17[110] |
| t*c*c*a*t*g*a*c*g*t*t*c*c*t*g*a*c*g*t*t*AAGAGTTTCTGCGGAGTGAATAACCTTTTTTTTTTT | 32[120] | 51[110] |
| t*c*c*a*t*g*a*c*g*t*t*c*c*t*g*a*c*g*t*t*TGGAAAGCAACGAGATGAAACAAACATTTTTTTTTT | 34[120] | 49[110] |
| t*c*c*a*t*g*a*c*g*t*t*c*c*t*g*a*c*g*t*t*CGTGAGTATTACGAAAATCGCGCAGATTTTTTTTTT | 36[120] | 47[110] |
| t*c*c*a*t*g*a*c*g*t*t*c*c*t*g*a*c*g*t*t*ATTAAATGTGAGCTTAGTTGCTATTTTTTTTTTTTT | 60[120] | 79[110] |
| t*c*c*a*t*g*a*c*g*t*t*c*c*t*g*a*c*g*t*t*CATTAAATTTTTGAAATCAGATATAGTTTTTTTTTT | 62[120] | 77[110] |
| t*c*c*a*t*g*a*c*g*t*t*c*c*t*g*a*c*g*t*t*ATTGTATAAGCAACAAGCCGTTTTTATTTTTTTTTT | 64[120] | 75[110] |
| t*c*c*a*t*g*a*c*g*t*t*c*c*t*g*a*c*g*t*t*GAGCAAACAAGAGTTTACGAGCATGTTTTTTTTTTT | 66[120] | 73[110] |
| t*c*c*a*t*g*a*c*g*t*t*c*c*t*g*a*c*g*t*t*ACCATAAATCAAATTGCCATCTTTTCTTTTTTTTTT | 88[120] | 107[110] |
| t*c*c*a*t*g*a*c*g*t*t*c*c*t*g*a*c*g*t*t*AATGTTTAGACTGCACCGTAATCAGTTTTTTTTTTT | 90[120] | 105[110] |
| t*c*c*a*t*g*a*c*g*t*t*c*c*t*g*a*c*g*t*t*AAATAGCGAGAGGCCATTACCATTAGTTTTTTTTTT | 92[120] | 103[110] |
| t*c*c*a*t*g*a*c*g*t*t*c*c*t*g*a*c*g*t*t*ACAGCTTGATACCGATAGTTGCGCCGTACGCCCCCTT | 116[120] | 107[79] |
| t*c*c*a*t*g*a*c*g*t*t*c*c*t*g*a*c*g*t*t*TAATTTTTTCACGTTGAAAATCTTTCGAG | 118[120] | 117[87] |
| t*c*c*a*t*g*a*c*g*t*t*c*c*t*g*a*c*g*t*t*CAGTTTCAGCGGAGTGAGAATAGAAAATGAAAATCAC | 120[120] | 103[79] |
| t*c*c*a*t*g*a*c*g*t*t*c*c*t*g*a*c*g*t*t*AGCCCTCATAGTTAGCGTAACATTTTGCT | 122[120] | 121[87] |
| **7.5nm spacing-CpG4-18 staples** |  |  |
| t*c*c*a*t*g*a*c*g*t*t*c*c*t*g*a*c*g*t*t*TTCCAGTTTGGAAGAAAATGCTGAATTTTTTTTTTT | 0[110] | 27[110] |
| t*c*c*a*t*g*a*c*g*t*t*c*c*t*g*a*c*g*t*t*GCCCTTCACCGCCAAGAATACGTGGCTTTTTTTTTT | 4[110] | 23[110] |
| t*c*c*a*t*g*a*c*g*t*t*c*c*t*g*a*c*g*t*t*TGCGTTGCGCTCACAGGAAAAACGCTTTTTTTTTTT | 8[110] | 19[110] |
| t*c*c*a*t*g*a*c*g*t*t*c*c*t*g*a*c*g*t*t*CCCGGGTACCGAGCCGAGTAAAAGAGTTTTTTTTTT | 12[110] | 15[110] |
| t*c*c*a*t*g*a*c*g*t*t*c*c*t*g*a*c*g*t*t*CATTACGCATCGCGAATCCTTGAAAATTTTTTTTTT | 30[110] | 53[110] |
| t*c*c*a*t*g*a*c*g*t*t*c*c*t*g*a*c*g*t*t*TGGAAAGCAACGAGATGAAACAAACATTTTTTTTTT | 34[110] | 49[110] |
| t*c*c*a*t*g*a*c*g*t*t*c*c*t*g*a*c*g*t*t*ATCAAACGCCGCGTTCAGGTTTAACGTTTTTTTTTT | 38[110] | 45[110] |
| t*c*c*a*t*g*a*c*g*t*t*c*c*t*g*a*c*g*t*t*GGAAGATCGCACTCCTTTTGATAAGATTTTTTTTTT | 56[110] | 83[110] |
| t*c*c*a*t*g*a*c*g*t*t*c*c*t*g*a*c*g*t*t*ATTAAATGTGAGCTTAGTTGCTATTTTTTTTTTTTT | 60[110] | 79[110] |
| t*c*c*a*t*g*a*c*g*t*t*c*c*t*g*a*c*g*t*t*ATTGTATAAGCAACAAGCCGTTTTTATTTTTTTTTT | 64[110] | 75[110] |
| t*c*c*a*t*g*a*c*g*t*t*c*c*t*g*a*c*g*t*t*AGCTATTTTTGAGGTTTATCAACAATTTTTTTTTTT | 68[110] | 71[110] |
| t*c*c*a*t*g*a*c*g*t*t*c*c*t*g*a*c*g*t*t*GCGGATTGCATCAGCCTCCCTCAGAGTTTTTTTTTT | 86[110] | 109[110] |
| t*c*c*a*t*g*a*c*g*t*t*c*c*t*g*a*c*g*t*t*AATGTTTAGACTGCACCGTAATCAGTTTTTTTTTTT | 90[110] | 105[110] |
| t*c*c*a*t*g*a*c*g*t*t*c*c*t*g*a*c*g*t*t*TCAACTAATGCAGGGTAAATATTGACTTTTTTTTTT | 94[110] | 101[110] |
| t*c*c*a*t*g*a*c*g*t*t*c*c*t*g*a*c*g*t*t*GGTAGCAACGGCTACAGAGGCTTTCCGCC | 112[110] | 112[72] |
| t*c*c*a*t*g*a*c*g*t*t*c*c*t*g*a*c*g*t*t*ACAGCTTGATACCGATAGTTGCGCCGTACGCCCCCTT | 116[110] | 107[79] |
| t*c*c*a*t*g*a*c*g*t*t*c*c*t*g*a*c*g*t*t*CAGTTTCAGCGGAGTGAGAATAGAAAATGAAAATCAC | 120[110] | 103[79] |
| t*c*c*a*t*g*a*c*g*t*t*c*c*t*g*a*c*g*t*t*CACTGAGTTTCGTCACCAGTACAAACCCGTAGAAAAT | 124[110] | 99[79] |
| **CpG2-dimer 1** |  |  |
| *t*c*c*a*t*g*a*c*g*t*t*c*c*t*g*a*c*g*t*tTCTGTCCATCACGAAACCAGCAATACTTTTTTTTTT | 14[110] | 39[110] |
| *t*c*c*a*t*g*a*c*g*t*t*c*c*t*g*a*c*g*t*tCATGGAAATACCTCGTATTGCTAAACTTTTTTTTTT | 18[110] | 35[110] |
| *t*c*c*a*t*g*a*c*g*t*t*c*c*t*g*a*c*g*t*tACAGACAATATTTTAACTATCGACATTTTTTTTTTT | 22[110] | 31[110] |
| *t*c*c*a*t*g*a*c*g*t*t*c*c*t*g*a*c*g*t*tAAGAGTTTCTGCGGAGTGAATAACCTTTTTTTTTTT | 32[110] | 51[110] |
| *t*c*c*a*t*g*a*c*g*t*t*c*c*t*g*a*c*g*t*tCGTGAGTATTACGAAAATCGCGCAGATTTTTTTTTT | 36[110] | 47[110] |
| *t*c*c*a*t*g*a*c*g*t*t*c*c*t*g*a*c*g*t*tACCATCGATTGTGCCATATCAAAATTTTTTTTTTTT | 40[110] | 43[110] |
| *t*c*c*a*t*g*a*c*g*t*t*c*c*t*g*a*c*g*t*tGGATTGACCGTAATAACGAGCGTCTTTTTTTTTTTT | 58[110] | 81[110] |
| *t*c*c*a*t*g*a*c*g*t*t*c*c*t*g*a*c*g*t*tCATTAAATTTTTGAAATCAGATATAGTTTTTTTTTT | 62[110] | 77[110] |
| *t*c*c*a*t*g*a*c*g*t*t*c*c*t*g*a*c*g*t*tGAGCAAACAAGAGTTTACGAGCATGTTTTTTTTTTT | 66[110] | 73[110] |
| *t*c*c*a*t*g*a*c*g*t*t*c*c*t*g*a*c*g*t*tAGATAAGTCCTGAAGGAATACCACATTTTTTTTTTT | 70[110] | 95[110] |
| *t*c*c*a*t*g*a*c*g*t*t*c*c*t*g*a*c*g*t*tTTTTCATCGTAGGGGGGTAATAGTAATTTTTTTTTT | 74[110] | 91[110] |
| *t*c*c*a*t*g*a*c*g*t*t*c*c*t*g*a*c*g*t*tTGCACCCAGCTACAGTCAGAAGCAAATTTTTTTTTT | 78[110] | 87[110] |
| *t*c*c*a*t*g*a*c*g*t*t*c*c*t*g*a*c*g*t*tGGAAATTATTCATAAACAACTTTCAATTTTTTTTTT | 100[110] | 121[110] |
| *t*c*c*a*t*g*a*c*g*t*t*c*c*t*g*a*c*g*t*tAGCGACAGAATCAGTGAATTTCTTAATTTTTTTTTT | 104[110] | 117[110] |
| *t*c*c*a*t*g*a*c*g*t*t*c*c*t*g*a*c*g*t*tCCGCCACCCTCAGGCATCGGAACGAGTTTTTTTTTT | 108[110] | 113[110] |
| *t*c*c*a*t*g*a*c*g*t*t*c*c*t*g*a*c*g*t*tATAACCGATATATTCGGTCGCGAAAGACA | 114[110] | 113[87] |
| *t*c*c*a*t*g*a*c*g*t*t*c*c*t*g*a*c*g*t*tTAATTTTTTCACGTTGAAAATCTTTCGAG | 118[110] | 117[87] |
| *t*c*c*a*t*g*a*c*g*t*t*c*c*t*g*a*c*g*t*tAGCCCTCATAGTTAGCGTAACATTTTGCT | 122[110] | 121[87] |
| *t*c*c*a*t*g*a*c*g*t*t*c*c*t*g*a*c*g*t*tTCTGTCCATCACGAAACCAGCAATACTTTTTTTTTT | 14[110] | 39[110] |
| **CpG2-dimer 2** |  |  |
| *t*c*c*a*t*g*a*c*g*t*t*c*c*t*g*a*c*g*t*tTTCCAGTTTGGAAGAAAATGCTGAATTTTTTTTTTT | 0[110] | 27[110] |
| *t*c*c*a*t*g*a*c*g*t*t*c*c*t*g*a*c*g*t*tCATTAATGAATCGTGGATTATTTACATTTTTTTTTT | 6[110] | 21[110] |
| *t*c*c*a*t*g*a*c*g*t*t*c*c*t*g*a*c*g*t*tCCCGGGTACCGAGCCGAGTAAAAGAGTTTTTTTTTT | 12[110] | 15[110] |
| *t*c*c*a*t*g*a*c*g*t*t*c*c*t*g*a*c*g*t*tTCTGTCCATCACGAAACCAGCAATACTTTTTTTTTT | 14[110] | 39[110] |
| *t*c*c*a*t*g*a*c*g*t*t*c*c*t*g*a*c*g*t*tTTGGCAGATTCACACTGGTGACCTGGTTTTTTTTTT | 20[110] | 33[110] |
| *t*c*c*a*t*g*a*c*g*t*t*c*c*t*g*a*c*g*t*tTTCGCGTCGTCTTCACAGCGATGCCAGAGTCTGTAGT | 26[110] | 26[64] |
| *t*c*c*a*t*g*a*c*g*t*t*c*c*t*g*a*c*g*t*tATTTGCACGTAAAGCCTGAGAGTCTGTTTTTTTTTT | 42[110] | 67[110] |
| *t*c*c*a*t*g*a*c*g*t*t*c*c*t*g*a*c*g*t*tGGCGAATTATTCATTGTTAAAATTCGTTTTTTTTTT | 46[110] | 63[110] |
| *t*c*c*a*t*g*a*c*g*t*t*c*c*t*g*a*c*g*t*tCATAGCGATAGCTAGTATCGGCCTCATTTTTTTTTT | 52[110] | 57[110] |
| *t*c*c*a*t*g*a*c*g*t*t*c*c*t*g*a*c*g*t*tGGAAGATCGCACTCCTTTTGATAAGATTTTTTTTTT | 56[110] | 83[110] |
| *t*c*c*a*t*g*a*c*g*t*t*c*c*t*g*a*c*g*t*tCATTAAATTTTTGAAATCAGATATAGTTTTTTTTTT | 62[110] | 77[110] |
| *t*c*c*a*t*g*a*c*g*t*t*c*c*t*g*a*c*g*t*tAGCTATTTTTGAGGTTTATCAACAATTTTTTTTTTT | 68[110] | 71[110] |
| *t*c*c*a*t*g*a*c*g*t*t*c*c*t*g*a*c*g*t*tAAGACAAAAGGGCGCATTCCACAGACTTTTTTTTTT | 98[110] | 123[110] |
| *t*c*c*a*t*g*a*c*g*t*t*c*c*t*g*a*c*g*t*tAGCGACAGAATCAGTGAATTTCTTAATTTTTTTTTT | 104[110] | 117[110] |
| *t*c*c*a*t*g*a*c*g*t*t*c*c*t*g*a*c*g*t*tGAAGTTTCCATTAAACGGGTAAAATACGT | 110[110] | 110[72] |
| *t*c*c*a*t*g*a*c*g*t*t*c*c*t*g*a*c*g*t*tGGTAGCAACGGCTACAGAGGCTTTCCGCC | 112[110] | 112[72] |
| *t*c*c*a*t*g*a*c*g*t*t*c*c*t*g*a*c*g*t*tTAATTTTTTCACGTTGAAAATCTTTCGAG | 118[110] | 117[87] |
| *t*c*c*a*t*g*a*c*g*t*t*c*c*t*g*a*c*g*t*tCACTGAGTTTCGTCACCAGTACAAACCCGTAGAAAAT | 124[110] | 99[79] |
| **3.5nm spacing-12 CpG strands** |  |  |
| *t*c*c*a*t*g*a*c*g*t*t*c*c*t*g*a*c*g*t*tTGGAAAGCAACGAGATGAAACAAACATTTTTTTTTT | 34[110] | 49[110] |
| *t*c*c*a*t*g*a*c*g*t*t*c*c*t*g*a*c*g*t*tCGTGAGTATTACGAAAATCGCGCAGATTTTTTTTTT | 36[110] | 47[110] |
| *t*c*c*a*t*g*a*c*g*t*t*c*c*t*g*a*c*g*t*tTCAGATGAATATACAAAAACAGGAAGTTTTTTTTTT | 44[110] | 65[110] |
| *t*c*c*a*t*g*a*c*g*t*t*c*c*t*g*a*c*g*t*tGGCGAATTATTCATTGTTAAAATTCGTTTTTTTTTT | 46[110] | 63[110] |
| *t*c*c*a*t*g*a*c*g*t*t*c*c*t*g*a*c*g*t*tTCAAGAAAACAAAAGCTTTCATCAACTTTTTTTTTT | 48[110] | 61[110] |
| *t*c*c*a*t*g*a*c*g*t*t*c*c*t*g*a*c*g*t*tCATTAAATTTTTGAAATCAGATATAGTTTTTTTTTT | 62[110] | 77[110] |
| *t*c*c*a*t*g*a*c*g*t*t*c*c*t*g*a*c*g*t*tATTGTATAAGCAACAAGCCGTTTTTATTTTTTTTTT | 64[110] | 75[110] |
| *t*c*c*a*t*g*a*c*g*t*t*c*c*t*g*a*c*g*t*tAGAAACCAATCAAACGATAAAAACCATTTTTTTTTT | 72[110] | 93[110] |
| *t*c*c*a*t*g*a*c*g*t*t*c*c*t*g*a*c*g*t*tTTTTCATCGTAGGGGGGTAATAGTAATTTTTTTTTT | 74[110] | 91[110] |
| *t*c*c*a*t*g*a*c*g*t*t*c*c*t*g*a*c*g*t*tAAGGCTTATCCGGGAAAACGAGAATGTTTTTTTTTT | 76[110] | 89[110] |
| *t*c*c*a*t*g*a*c*g*t*t*c*c*t*g*a*c*g*t*tAATGTTTAGACTGCACCGTAATCAGTTTTTTTTTTT | 90[110] | 105[110] |
| *t*c*c*a*t*g*a*c*g*t*t*c*c*t*g*a*c*g*t*tAAATAGCGAGAGGCCATTACCATTAGTTTTTTTTTT | 92[110] | 103[110] |
| **3.5nm spacing-35 CpG strands** |  |  |
| t*c*c*a*t*g*a*c*g*t*t*c*c*t*g*a*c*g*t*t*TCTGTCCATCACGAAACCAGCAATACTTTTTTTTTT | 14[110] | 39[110] |
| t*c*c*a*t*g*a*c*g*t*t*c*c*t*g*a*c*g*t*t*AACATCACTTGCCACTCAACGAGCAGTTTTTTTTTT | 16[110] | 37[110] |
| t*c*c*a*t*g*a*c*g*t*t*c*c*t*g*a*c*g*t*t*CATGGAAATACCTCGTATTGCTAAACTTTTTTTTTT | 18[110] | 35[110] |
| t*c*c*a*t*g*a*c*g*t*t*c*c*t*g*a*c*g*t*t*TTGGCAGATTCACACTGGTGACCTGGTTTTTTTTTT | 20[110] | 33[110] |
| t*c*c*a*t*g*a*c*g*t*t*c*c*t*g*a*c*g*t*t*ACAGACAATATTTTAACTATCGACATTTTTTTTTTT | 22[110] | 31[110] |
| t*c*c*a*t*g*a*c*g*t*t*c*c*t*g*a*c*g*t*t*CATTACGCATCGCGAATCCTTGAAAATTTTTTTTTT | 30[110] | 53[110] |
| t*c*c*a*t*g*a*c*g*t*t*c*c*t*g*a*c*g*t*t*AAGAGTTTCTGCGGAGTGAATAACCTTTTTTTTTTT | 32[110] | 51[110] |
| t*c*c*a*t*g*a*c*g*t*t*c*c*t*g*a*c*g*t*t*TGGAAAGCAACGAGATGAAACAAACATTTTTTTTTT | 34[110] | 49[110] |
| t*c*c*a*t*g*a*c*g*t*t*c*c*t*g*a*c*g*t*t*CGTGAGTATTACGAAAATCGCGCAGATTTTTTTTTT | 36[110] | 47[110] |
| t*c*c*a*t*g*a*c*g*t*t*c*c*t*g*a*c*g*t*t*ATCAAACGCCGCGTTCAGGTTTAACGTTTTTTTTTT | 38[110] | 45[110] |
| t*c*c*a*t*g*a*c*g*t*t*c*c*t*g*a*c*g*t*t*ATTTGCACGTAAAGCCTGAGAGTCTGTTTTTTTTTT | 42[110] | 67[110] |
| t*c*c*a*t*g*a*c*g*t*t*c*c*t*g*a*c*g*t*t*TCAGATGAATATACAAAAACAGGAAGTTTTTTTTTT | 44[110] | 65[110] |
| t*c*c*a*t*g*a*c*g*t*t*c*c*t*g*a*c*g*t*t*GGCGAATTATTCATTGTTAAAATTCGTTTTTTTTTT | 46[110] | 63[110] |
| t*c*c*a*t*g*a*c*g*t*t*c*c*t*g*a*c*g*t*t*TCAAGAAAACAAAAGCTTTCATCAACTTTTTTTTTT | 48[110] | 61[110] |
| t*c*c*a*t*g*a*c*g*t*t*c*c*t*g*a*c*g*t*t*TGCTTCTGTAAATGGGAACAAACGGCTTTTTTTTTT | 50[110] | 59[110] |
| t*c*c*a*t*g*a*c*g*t*t*c*c*t*g*a*c*g*t*t*GGATTGACCGTAATAACGAGCGTCTTTTTTTTTTTT | 58[110] | 81[110] |
| t*c*c*a*t*g*a*c*g*t*t*c*c*t*g*a*c*g*t*t*ATTAAATGTGAGCTTAGTTGCTATTTTTTTTTTTTT | 60[110] | 79[110] |
| t*c*c*a*t*g*a*c*g*t*t*c*c*t*g*a*c*g*t*t*CATTAAATTTTTGAAATCAGATATAGTTTTTTTTTT | 62[110] | 77[110] |
| t*c*c*a*t*g*a*c*g*t*t*c*c*t*g*a*c*g*t*t*ATTGTATAAGCAACAAGCCGTTTTTATTTTTTTTTT | 64[110] | 75[110] |
| t*c*c*a*t*g*a*c*g*t*t*c*c*t*g*a*c*g*t*t*GAGCAAACAAGAGTTTACGAGCATGTTTTTTTTTTT | 66[110] | 73[110] |
| t*c*c*a*t*g*a*c*g*t*t*c*c*t*g*a*c*g*t*t*AGATAAGTCCTGAAGGAATACCACATTTTTTTTTTT | 70[110] | 95[110] |
| t*c*c*a*t*g*a*c*g*t*t*c*c*t*g*a*c*g*t*t*AGAAACCAATCAAACGATAAAAACCATTTTTTTTTT | 72[110] | 93[110] |
| t*c*c*a*t*g*a*c*g*t*t*c*c*t*g*a*c*g*t*t*TTTTCATCGTAGGGGGGTAATAGTAATTTTTTTTTT | 74[110] | 91[110] |
| t*c*c*a*t*g*a*c*g*t*t*c*c*t*g*a*c*g*t*t*AAGGCTTATCCGGGAAAACGAGAATGTTTTTTTTTT | 76[110] | 89[110] |
| t*c*c*a*t*g*a*c*g*t*t*c*c*t*g*a*c*g*t*t*TGCACCCAGCTACAGTCAGAAGCAAATTTTTTTTTT | 78[110] | 87[110] |
| t*c*c*a*t*g*a*c*g*t*t*c*c*t*g*a*c*g*t*t*GCGGATTGCATCAGCCTCCCTCAGAGTTTTTTTTTT | 86[110] | 109[110] |
| t*c*c*a*t*g*a*c*g*t*t*c*c*t*g*a*c*g*t*t*ACCATAAATCAAATTGCCATCTTTTCTTTTTTTTTT | 88[110] | 107[110] |
| t*c*c*a*t*g*a*c*g*t*t*c*c*t*g*a*c*g*t*t*AATGTTTAGACTGCACCGTAATCAGTTTTTTTTTTT | 90[110] | 105[110] |
| t*c*c*a*t*g*a*c*g*t*t*c*c*t*g*a*c*g*t*t*AAATAGCGAGAGGCCATTACCATTAGTTTTTTTTTT | 92[110] | 103[110] |
| t*c*c*a*t*g*a*c*g*t*t*c*c*t*g*a*c*g*t*t*TCAACTAATGCAGGGTAAATATTGACTTTTTTTTTT | 94[110] | 101[110] |
| t*c*c*a*t*g*a*c*g*t*t*c*c*t*g*a*c*g*t*t*AAGACAAAAGGGCGCATTCCACAGACTTTTTTTTTT | 98[110] | 123[110] |
| t*c*c*a*t*g*a*c*g*t*t*c*c*t*g*a*c*g*t*t*GGAAATTATTCATAAACAACTTTCAATTTTTTTTTT | 100[110] | 121[110] |
| t*c*c*a*t*g*a*c*g*t*t*c*c*t*g*a*c*g*t*t*CAAGGCCGGAAACGAATTGCGAATAATTTTTTTTTT | 102[110] | 119[110] |
| t*c*c*a*t*g*a*c*g*t*t*c*c*t*g*a*c*g*t*t*AGCGACAGAATCAGTGAATTTCTTAATTTTTTTTTT | 104[110] | 117[110] |
| t*c*c*a*t*g*a*c*g*t*t*c*c*t*g*a*c*g*t*t*ATAATCAAAATCACCATCGCCCACGCTTTTTTTTTT | 106[110] | 115[110] |
| **3.5nm spacing-63 CpG strands** |  |  |
| *t*c*c*a*t*g*a*c*g*t*t*c*c*t*g*a*c*g*t*tTTCCAGTTTGGAAGAAAATGCTGAATTTTTTTTTTT | 0[110] | 27[110] |
| *t*c*c*a*t*g*a*c*g*t*t*c*c*t*g*a*c*g*t*tTGCCCCAGCAGGCCGAACTGATAGCCTTTTTTTTTT | 2[110] | 25[110] |
| *t*c*c*a*t*g*a*c*g*t*t*c*c*t*g*a*c*g*t*tGCCCTTCACCGCCAAGAATACGTGGCTTTTTTTTTT | 4[110] | 23[110] |
| *t*c*c*a*t*g*a*c*g*t*t*c*c*t*g*a*c*g*t*tCATTAATGAATCGTGGATTATTTACATTTTTTTTTT | 6[110] | 21[110] |
| *t*c*c*a*t*g*a*c*g*t*t*c*c*t*g*a*c*g*t*tTGCGTTGCGCTCACAGGAAAAACGCTTTTTTTTTTT | 8[110] | 19[110] |
| *t*c*c*a*t*g*a*c*g*t*t*c*c*t*g*a*c*g*t*tAAATTGTTATCCGTTTGATTAGTAATTTTTTTTTTT | 10[110] | 17[110] |
| *t*c*c*a*t*g*a*c*g*t*t*c*c*t*g*a*c*g*t*tCCCGGGTACCGAGCCGAGTAAAAGAGTTTTTTTTTT | 12[110] | 15[110] |
| *t*c*c*a*t*g*a*c*g*t*t*c*c*t*g*a*c*g*t*tTCTGTCCATCACGAAACCAGCAATACTTTTTTTTTT | 14[110] | 39[110] |
| *t*c*c*a*t*g*a*c*g*t*t*c*c*t*g*a*c*g*t*tAACATCACTTGCCACTCAACGAGCAGTTTTTTTTTT | 16[110] | 37[110] |
| *t*c*c*a*t*g*a*c*g*t*t*c*c*t*g*a*c*g*t*tCATGGAAATACCTCGTATTGCTAAACTTTTTTTTTT | 18[110] | 35[110] |
| *t*c*c*a*t*g*a*c*g*t*t*c*c*t*g*a*c*g*t*tTTGGCAGATTCACACTGGTGACCTGGTTTTTTTTTT | 20[110] | 33[110] |
| *t*c*c*a*t*g*a*c*g*t*t*c*c*t*g*a*c*g*t*tACAGACAATATTTTAACTATCGACATTTTTTTTTTT | 22[110] | 31[110] |
| *t*c*c*a*t*g*a*c*g*t*t*c*c*t*g*a*c*g*t*tCTAAAACATCGCCTTCGCATCAAAGGTTTTTTTTTT | 24[110] | 29[110] |
| *t*c*c*a*t*g*a*c*g*t*t*c*c*t*g*a*c*g*t*tTTCGCGTCGTCTTCACAGCGATGCCAGAGTCTGTAGT | 26[110] | 26[64] |
| *t*c*c*a*t*g*a*c*g*t*t*c*c*t*g*a*c*g*t*tAGAGTGAGATCGGTTCTGGTGCCGGATTTTTTTTTT | 28[110] | 55[110] |
| *t*c*c*a*t*g*a*c*g*t*t*c*c*t*g*a*c*g*t*tCATTACGCATCGCGAATCCTTGAAAATTTTTTTTTT | 30[110] | 53[110] |
| *t*c*c*a*t*g*a*c*g*t*t*c*c*t*g*a*c*g*t*tAAGAGTTTCTGCGGAGTGAATAACCTTTTTTTTTTT | 32[110] | 51[110] |
| *t*c*c*a*t*g*a*c*g*t*t*c*c*t*g*a*c*g*t*tTGGAAAGCAACGAGATGAAACAAACATTTTTTTTTT | 34[110] | 49[110] |
| *t*c*c*a*t*g*a*c*g*t*t*c*c*t*g*a*c*g*t*tCGTGAGTATTACGAAAATCGCGCAGATTTTTTTTTT | 36[110] | 47[110] |
| *t*c*c*a*t*g*a*c*g*t*t*c*c*t*g*a*c*g*t*tATCAAACGCCGCGTTCAGGTTTAACGTTTTTTTTTT | 38[110] | 45[110] |
| *t*c*c*a*t*g*a*c*g*t*t*c*c*t*g*a*c*g*t*tACCATCGATTGTGCCATATCAAAATTTTTTTTTTTT | 40[110] | 43[110] |
| *t*c*c*a*t*g*a*c*g*t*t*c*c*t*g*a*c*g*t*tATTTGCACGTAAAGCCTGAGAGTCTGTTTTTTTTTT | 42[110] | 67[110] |
| *t*c*c*a*t*g*a*c*g*t*t*c*c*t*g*a*c*g*t*tTCAGATGAATATACAAAAACAGGAAGTTTTTTTTTT | 44[110] | 65[110] |
| *t*c*c*a*t*g*a*c*g*t*t*c*c*t*g*a*c*g*t*tGGCGAATTATTCATTGTTAAAATTCGTTTTTTTTTT | 46[110] | 63[110] |
| *t*c*c*a*t*g*a*c*g*t*t*c*c*t*g*a*c*g*t*tTCAAGAAAACAAAAGCTTTCATCAACTTTTTTTTTT | 48[110] | 61[110] |
| *t*c*c*a*t*g*a*c*g*t*t*c*c*t*g*a*c*g*t*tTGCTTCTGTAAATGGGAACAAACGGCTTTTTTTTTT | 50[110] | 59[110] |
| *t*c*c*a*t*g*a*c*g*t*t*c*c*t*g*a*c*g*t*tCATAGCGATAGCTAGTATCGGCCTCATTTTTTTTTT | 52[110] | 57[110] |
| *t*c*c*a*t*g*a*c*g*t*t*c*c*t*g*a*c*g*t*tAACCAGGCAAAGCGCCATTCGCCATTCAGGCTGCGCA | 54[110] | 54[64] |
| *t*c*c*a*t*g*a*c*g*t*t*c*c*t*g*a*c*g*t*tGGAAGATCGCACTCCTTTTGATAAGATTTTTTTTTT | 56[110] | 83[110] |
| *t*c*c*a*t*g*a*c*g*t*t*c*c*t*g*a*c*g*t*tGGATTGACCGTAATAACGAGCGTCTTTTTTTTTTTT | 58[110] | 81[110] |
| *t*c*c*a*t*g*a*c*g*t*t*c*c*t*g*a*c*g*t*tATTAAATGTGAGCTTAGTTGCTATTTTTTTTTTTTT | 60[110] | 79[110] |
| *t*c*c*a*t*g*a*c*g*t*t*c*c*t*g*a*c*g*t*tCATTAAATTTTTGAAATCAGATATAGTTTTTTTTTT | 62[110] | 77[110] |
| *t*c*c*a*t*g*a*c*g*t*t*c*c*t*g*a*c*g*t*tATTGTATAAGCAACAAGCCGTTTTTATTTTTTTTTT | 64[110] | 75[110] |
| *t*c*c*a*t*g*a*c*g*t*t*c*c*t*g*a*c*g*t*tGAGCAAACAAGAGTTTACGAGCATGTTTTTTTTTTT | 66[110] | 73[110] |
| *t*c*c*a*t*g*a*c*g*t*t*c*c*t*g*a*c*g*t*tAGCTATTTTTGAGGTTTATCAACAATTTTTTTTTTT | 68[110] | 71[110] |
| *t*c*c*a*t*g*a*c*g*t*t*c*c*t*g*a*c*g*t*tAGATAAGTCCTGAAGGAATACCACATTTTTTTTTTT | 70[110] | 95[110] |
| *t*c*c*a*t*g*a*c*g*t*t*c*c*t*g*a*c*g*t*tAGAAACCAATCAAACGATAAAAACCATTTTTTTTTT | 72[110] | 93[110] |
| *t*c*c*a*t*g*a*c*g*t*t*c*c*t*g*a*c*g*t*tTTTTCATCGTAGGGGGGTAATAGTAATTTTTTTTTT | 74[110] | 91[110] |
| *t*c*c*a*t*g*a*c*g*t*t*c*c*t*g*a*c*g*t*tAAGGCTTATCCGGGAAAACGAGAATGTTTTTTTTTT | 76[110] | 89[110] |
| *t*c*c*a*t*g*a*c*g*t*t*c*c*t*g*a*c*g*t*tTGCACCCAGCTACAGTCAGAAGCAAATTTTTTTTTT | 78[110] | 87[110] |
| *t*c*c*a*t*g*a*c*g*t*t*c*c*t*g*a*c*g*t*tTCCAGAGCCTAATCGGAAGCAAACTCTTTTTTTTTT | 80[110] | 85[110] |
| *t*c*c*a*t*g*a*c*g*t*t*c*c*t*g*a*c*g*t*tGGTCATTTTTGCGGATGGCTTAGAGCTTA | 82[110] | 82[72] |
| *t*c*c*a*t*g*a*c*g*t*t*c*c*t*g*a*c*g*t*tCAACAGGTCAGGAACTTTTTCATGAGTTTTTTTTTT | 84[110] | 111[110] |
| *t*c*c*a*t*g*a*c*g*t*t*c*c*t*g*a*c*g*t*tGCGGATTGCATCAGCCTCCCTCAGAGTTTTTTTTTT | 86[110] | 109[110] |
| *t*c*c*a*t*g*a*c*g*t*t*c*c*t*g*a*c*g*t*tACCATAAATCAAATTGCCATCTTTTCTTTTTTTTTT | 88[110] | 107[110] |
| *t*c*c*a*t*g*a*c*g*t*t*c*c*t*g*a*c*g*t*tAATGTTTAGACTGCACCGTAATCAGTTTTTTTTTTT | 90[110] | 105[110] |
| *t*c*c*a*t*g*a*c*g*t*t*c*c*t*g*a*c*g*t*tAAATAGCGAGAGGCCATTACCATTAGTTTTTTTTTT | 92[110] | 103[110] |
| *t*c*c*a*t*g*a*c*g*t*t*c*c*t*g*a*c*g*t*tTCAACTAATGCAGGGTAAATATTGACTTTTTTTTTT | 94[110] | 101[110] |
| *t*c*c*a*t*g*a*c*g*t*t*c*c*t*g*a*c*g*t*tAACATTATTACAGTTTACCAGCGCCATTTTTTTTTT | 96[110] | 99[110] |
| *t*c*c*a*t*g*a*c*g*t*t*c*c*t*g*a*c*g*t*tAAGACAAAAGGGCGCATTCCACAGACTTTTTTTTTT | 98[110] | 123[110] |
| *t*c*c*a*t*g*a*c*g*t*t*c*c*t*g*a*c*g*t*tGGAAATTATTCATAAACAACTTTCAATTTTTTTTTT | 100[110] | 121[110] |
| *t*c*c*a*t*g*a*c*g*t*t*c*c*t*g*a*c*g*t*tCAAGGCCGGAAACGAATTGCGAATAATTTTTTTTTT | 102[110] | 119[110] |
| *t*c*c*a*t*g*a*c*g*t*t*c*c*t*g*a*c*g*t*tAGCGACAGAATCAGTGAATTTCTTAATTTTTTTTTT | 104[110] | 117[110] |
| *t*c*c*a*t*g*a*c*g*t*t*c*c*t*g*a*c*g*t*tATAATCAAAATCACCATCGCCCACGCTTTTTTTTTT | 106[110] | 115[110] |
| *t*c*c*a*t*g*a*c*g*t*t*c*c*t*g*a*c*g*t*tCCGCCACCCTCAGGCATCGGAACGAGTTTTTTTTTT | 108[110] | 113[110] |
| *t*c*c*a*t*g*a*c*g*t*t*c*c*t*g*a*c*g*t*tGAAGTTTCCATTAAACGGGTAAAATACGT | 110[110] | 110[72] |
| *t*c*c*a*t*g*a*c*g*t*t*c*c*t*g*a*c*g*t*tGGTAGCAACGGCTACAGAGGCTTTCCGCC | 112[110] | 112[72] |
| *t*c*c*a*t*g*a*c*g*t*t*c*c*t*g*a*c*g*t*tATAACCGATATATTCGGTCGCGAAAGACA | 114[110] | 113[87] |
| *t*c*c*a*t*g*a*c*g*t*t*c*c*t*g*a*c*g*t*tACAGCTTGATACCGATAGTTGCGCCGTACGCCCCCTT | 116[110] | 107[79] |
| *t*c*c*a*t*g*a*c*g*t*t*c*c*t*g*a*c*g*t*tTAATTTTTTCACGTTGAAAATCTTTCGAG | 118[110] | 117[87] |
| *t*c*c*a*t*g*a*c*g*t*t*c*c*t*g*a*c*g*t*tCAGTTTCAGCGGAGTGAGAATAGAAAATGAAAATCAC | 120[110] | 103[79] |
| *t*c*c*a*t*g*a*c*g*t*t*c*c*t*g*a*c*g*t*tAGCCCTCATAGTTAGCGTAACATTTTGCT | 122[110] | 121[87] |
| *t*c*c*a*t*g*a*c*g*t*t*c*c*t*g*a*c*g*t*tCACTGAGTTTCGTCACCAGTACAAACCCGTAGAAAAT | 124[110] | 99[79] |
| **irregular spacing-CpGi-18 staples** |  |  |
| t*c*c*a*t*g*a*c*g*t*t*c*c*t*g*a*c*g*t*t*TTCCAGTTTGGAAGAAAATGCTGAATTTTTTTTTTT | 0[110] | 27[110] |
| t*c*c*a*t*g*a*c*g*t*t*c*c*t*g*a*c*g*t*t*TGCCCCAGCAGGCCGAACTGATAGCCTTTTTTTTTT | 2[110] | 25[110] |
| t*c*c*a*t*g*a*c*g*t*t*c*c*t*g*a*c*g*t*t*TGCGTTGCGCTCACAGGAAAAACGCTTTTTTTTTTT | 8[110] | 19[110] |
| t*c*c*a*t*g*a*c*g*t*t*c*c*t*g*a*c*g*t*t*TCTGTCCATCACGAAACCAGCAATACTTTTTTTTTT | 14[110] | 39[110] |
| t*c*c*a*t*g*a*c*g*t*t*c*c*t*g*a*c*g*t*t*TTGGCAGATTCACACTGGTGACCTGGTTTTTTTTTT | 20[110] | 33[110] |
| t*c*c*a*t*g*a*c*g*t*t*c*c*t*g*a*c*g*t*t*AGAGTGAGATCGGTTCTGGTGCCGGATTTTTTTTTT | 28[110] | 55[110] |
| t*c*c*a*t*g*a*c*g*t*t*c*c*t*g*a*c*g*t*t*CGTGAGTATTACGAAAATCGCGCAGATTTTTTTTTT | 36[110] | 47[110] |
| t*c*c*a*t*g*a*c*g*t*t*c*c*t*g*a*c*g*t*t*ATTTGCACGTAAAGCCTGAGAGTCTGTTTTTTTTTT | 42[110] | 67[110] |
| t*c*c*a*t*g*a*c*g*t*t*c*c*t*g*a*c*g*t*t*GGCGAATTATTCATTGTTAAAATTCGTTTTTTTTTT | 46[110] | 63[110] |
| t*c*c*a*t*g*a*c*g*t*t*c*c*t*g*a*c*g*t*t*TCAAGAAAACAAAAGCTTTCATCAACTTTTTTTTTT | 48[110] | 61[110] |
| t*c*c*a*t*g*a*c*g*t*t*c*c*t*g*a*c*g*t*t*GGATTGACCGTAATAACGAGCGTCTTTTTTTTTTTT | 58[110] | 81[110] |
| t*c*c*a*t*g*a*c*g*t*t*c*c*t*g*a*c*g*t*t*AGCTATTTTTGAGGTTTATCAACAATTTTTTTTTTT | 68[110] | 71[110] |
| t*c*c*a*t*g*a*c*g*t*t*c*c*t*g*a*c*g*t*t*AGAAACCAATCAAACGATAAAAACCATTTTTTTTTT | 72[110] | 93[110] |
| t*c*c*a*t*g*a*c*g*t*t*c*c*t*g*a*c*g*t*t*CAACAGGTCAGGAACTTTTTCATGAGTTTTTTTTTT | 84[110] | 111[110] |
| t*c*c*a*t*g*a*c*g*t*t*c*c*t*g*a*c*g*t*t*AATGTTTAGACTGCACCGTAATCAGTTTTTTTTTTT | 90[110] | 105[110] |
| t*c*c*a*t*g*a*c*g*t*t*c*c*t*g*a*c*g*t*t*AACATTATTACAGTTTACCAGCGCCATTTTTTTTTT | 96[110] | 99[110] |
| t*c*c*a*t*g*a*c*g*t*t*c*c*t*g*a*c*g*t*t*AAGACAAAAGGGCGCATTCCACAGACTTTTTTTTTT | 98[110] | 123[110] |
| t*c*c*a*t*g*a*c*g*t*t*c*c*t*g*a*c*g*t*t*CCGCCACCCTCAGGCATCGGAACGAGTTTTTTTTTT | 108[110] | 113[110] |

**Table S3.** **CpG-containing staple sequences** **for various CpG spacing patterns.** The CpG strands were self-ligated or fabricated by IDT company. The CpG strands replace original poly-T strands (orange) at desired helix positions as shown in the second column. For all the CpG-containing staple strands, a sequence of poly T oligonucleotide at the 5´ end or 3´ end (or both) was substituted by the CpG sequence. The majority of the staple stands have the CpG sequence on the 5´end to maintain equivalent orientation of CpG across all designs. CpG1 has 3´-3´ linkage to achieve same CpG orientation. The positions in the second column correspond to the labeled helices in Fig. S4 (helices are numbered between 0 and 125). * refers to the PS modification of CpG oligonucleotides. In the second and third columns, the number before the square brackets refers to the helix position; the number inside the square brackets refers to the nucleotide base position corresponding to the CaDNAno file.

| Antibodies | Fluorophores | Catalog No.  (Mostly from Biolegend) |  | Antibodies | Fluorophores | Catalog No.  (Mostly from Biolegend) |
| --- | --- | --- | --- | --- | --- | --- |
| CD11c | BV570 | 117331 |  | CD3 | BV570 | 100225 |
| CD40 | PE | 124610 |  | CD4 | PE | 100408 |
| CD80 | APC/fire750 | 561134 |  | CD8 | APC/CY7 | 100714 |
| CD86 | PE/CY7 | 105014 |  | CD69 | PE-CY7 | 104512 |
| MHC-II | BV421 | 107632 |  | CD25 | BV711 | 102024 |
| PD-L1 | BV711 | 124319 |  | CD62L | PerCP | 104430 |
| DEC205 | PerCP/Cy5.5 | 138208 |  | CD127 | APC | 135012 |
| Viability | Zombie UV | 423108 |  | CD44 | AF700 | 103026 |
| CD11b | BV605 | 101257 |  | FoxP3 | BV421 | 126419 |
| Gr-1 | FITC | 108406 |  | IL-2 | PE-cy5 | 503824 |
| F4/80 | PE594 | 123146 |  | IFNγ | BV510 | 505841 |
| CD206 | BV650 | 141723 |  | IL-4 | PE594 | 504132 |
| NK1.1 | PE | 156504 |  | TNFα | PE594 | 506346 |
| TLR9 | FITC | ab58864 |  | CD8 Tetramer | BV421 | [MBL-TB-5001-4](https://www.mblintl.com/products/tb-5001-4) |
| MyD88 | / | ab135693 |  | Goat Anti-Rabbit IgG H&L | AF594 | ab150080 |

**Table S4**. Antibodies used for flow cytometry. These antibodies were grouped into different panels depending on the experimental goals.

**Supplemental methods**

**Agarose gel electrophoresis**

Folded SQB samples or purified SQB samples (CpG and/or OVA conjugated) were subjected to 2% native agarose gel electrophoresis at 70 V for 2.5 hours (gel prepared in 0.5 × TBE buffer supplemented with 11 mM MgCl_2_ and/or 0.005% (v/v) EtBr or SYBR safe). The gel was imaged using a Typhoon imager.

**TEM analysis**

The structural integrity of the SQB was verified using negative stain transmission electron microscopy (TEM). Prior to adding the samples, grids were cleaned using plasma discharge for 30 seconds. 3.0 µL of 4-10 nM SQB solution was then deposited on a carbon coated Formvar grid (Electron Microscopy Sciences). After 3 minutes, the sample was wicked from the grid by gently blotting the grid with the edge of the filter paper. A drop of uranyl formate solution (2% w/v in H_2_O) was then deposited onto the grid for 30 seconds, and the excess solution was wicked using filter paper. Studies were conducted using a JEOL JEM-1400 transmission electron microscope in brightfield mode at 80 kV.

**Cell uptake assay**

Cell uptake were performed using HeLa tumor cell line and Human Embryonic Kidney cells (HEK293) maintained in high-glucose Dulbecco’s modified Eagle medium (DMEM) (Gibco, Gaithersburg) and 10% fetal bovine serum (FBS) (Lonza, Wakersville) with penicillin–streptomycin. For flow cytometry, HEK293 cells were seeded at a density of 250,000 cells/mL into tissue culture treated 48 well plates (BD Life Sciences) and allowed to grow for 24 hours in 200 µL of media. CpG-SQBs with various spacings were added to a final concentration of 0.1–1 nM and incubated with cells. All samples were performed in triplicate.

**Western Blot**

Cells were plated and stimulated with different versions of vaccines. Following stimulation, cells were lysed using Pierce RIPA buffer (Thermo Scientific). Protein levels were detected using Pierce BCA protein assay (Thermo Fisher Scientific, 23227). Samples were then prepared combined with 4x LDS sample buffer and DTT before being sonicated for 15 minutes followed by a 10-minute incubation at 70°C. Proteins were separated using an SDS Page gel before transfer. Thermo Fisher iBlot 2 Dry Blotting System was used to transfer proteins to the membrane. After transfer, the membrane was blocked for 30 minutes in 2% Dry Milk Powder (RPI, M17200) in PBS-T (1% Tween 20 in Dulbecco’s Phosphate Buffered Saline) and incubated with a primary antibody then a second antibody. Primary antibodies used included anti-mouse TLR9 (Abcam, ab58864), anti-mouse TLR13 (Novus, NBP2-24539SS), and anti-GAPDH (Biovision, A1814). Secondary antibodies used included Anti-Rabbit IgG (H+L), Mouse/Human ads-HRP (Southern Biotech, 4050-05) and Mouse IgG HRP-conjugated Antibody (R&D Systems, HAF007).

**Liposome Preparation**

First, 6.2mg of POPC (Anatrace, P516) was measured into a glass tube and dissolved in sufficient amount of chloroform (VWR, BDH83627.400). The tube was then covered with parafilm, and a constant stream of Nitrogen gas was blown into the tube until all chloroform evaporated. Then, the tube was dried in a vacuum overnight. Lipids were then dissolved in 1 × TE (5 mM Tris and 1 mM EDTA) with 1% sodium cholate (VWR, 101183-986) to a final concentration of 6.25mM by sonicating for 1 hour. Dissolved lipids were incubated at 4°C for 1 hour with appropriate amounts of CpG and OVA-AF488 (Ovalbumin, Alexa Fluor™ 488 Conjugate, ThermoFisher, O34781). Incorporation frequencies were determined previously using a standard curve made by measuring fluorescent signals of liposomes after incorporation of various concentrations of OVA-AF488 or Cy5-tagged oligonucleotide. This protocol was optimized to ensure that the CpG and OVA were incorporated at an equivalent dose to that delivered by the SQB. Bio-Beads SM-2 Adsorbents (Bio-Rad, 1523920) were prepared via 3 washes with 0.5mL 1 × TE followed by spinning at 3 minutes at 16000 rcf and removing the floating, inactivated beads after each step. Activated Bio-beads were then added to each sample are a ratio of 300mg of Bio-beads to 250 µL of sample and gently rotated at 4°C overnight. Liposomes were concentrated using a 30 kDa Amicon Ultra-0.5 Centrifugal Filter Unit (Millipore Sigma, UFC5030) and then diluted with 1 × TE to the desired final concentration.

**Neoantigen peptide antigen conjugation with ‘handle’ oligonucleotide**

A ‘anti-juno handle’ oligonucleotide, which corresponds to 24 sites on the extruding face of the SQB, were ordered from IDT with an amine modification on the 5’ terminal (NH2-TTCTAGGGTTAAAAGGGGACG). These handle strands will replace some of the poly-T oligonucleotides on the extruding face of the SQB. The neoantigen peptides were ordered from GenScript with an azide-modified lysine on the N terminal. The ‘handle’ oligonucleotide was diluted to a final concentration of 1000 µM with phosphate buffer (pH 8.0). The neoantigen peptides were dissolved in DMSO, according to the manufacturer’s instructions. The oligonucleotide was conjugated with dibenzocyclooctyne-N-hydroxysuccinimidyl ester (DBCO-NHS ester; Millipore, #761524). The DBCO-NHS ester (diluted in DMSO to 2000 µM concentration) was incubated with the oligonucleotide (diluted in phosphate buffer pH 8.0 to 200 µM concentration) in 1:1 volume ratio and incubated overnight at ambient temperature. The final concentration of DBCO-NHS ester exceeded the concentration of the oligonucleotide 10:1, to ensure that all oligonucleotides were successfully conjugated with DBCO-NHS ester. After incubation, the product was purified via Illustra NAP column (GE Healthcare Life Sciences, #17-0852-02) and eluted with sterile water. The flow-through was concentrated via 3K Amicron Ultra Centrifugal Filter Unit (Millipore; #UFC500324) at 14,000 rcf for 15 minutes at 4°C. DBCO incorporation was confirmed by observation of an OD310 peak and final concentration was determined via measurement of A280 using the NanoDrop.

The N-terminal azide modified neoantigen peptides (M27: REGVELCPGNKYEMRRHGTTHSLVIHD; M33: DSGSPFPAAVILRDALHMARGLKYLHQ, M30: PSKPSFQEFVDWENVSPELNSTDQPFL, M47: GRGHLLGRLAAIVGKQVLLGRKVVVVR ) were conjugated with the DBCO-conjugated oligonucleotide via a copper-free click chemistry reaction. The peptide was combined with the DBCO-conjugated oligonucleotide at a ratio of 1.5: 1. 10X PBS was added to the reaction mixture so that the final reaction included 1XPBS. The reaction was incubated overnight at room temperature. 15% denaturing PAGE (dPAGE) was used to confirm successful conjugation of the peptide to the DBCO-modified oligonucleotide. 8% dPAGE was used to purify the samples. The reaction mixture was combined with formamide loading buffer (FLB) at a volume ratio of 1:1. The resulting mixture was loaded into a dPAGE gel, and the gel run at 250V for 50 minutes. The peptide-conjugated oligonucleotide band was observed via UV shadowing on a thin layer chromatography (TLC) plate and cut out with a razor blade. The gel was crushed with a pestle in a 15 mL Falcon tube. 1 × TE buffer was added in a 1:1 volume ratio and the solution was incubated overnight at 25°C, shaking at high speed. The gel solution was filtered through Freeze ‘N Squeeze DNA Gel Extraction Spin Columns (Bio-Rad; #7326165) according to the manufacturer’s protocol and then purified via ethanol precipitation. The peptide-oligonucleotide pellet was resuspended in 1 × PBS, and the concentration was determined via NanoDrop. Successful purification was confirmed via dPAGE analysis.

**Denaturing PAGE (dPAGE) verification of peptide-conjugated oligonucleotide**

Denaturing PAGE gels were cast using SequaGel-UreaGel System Kit (Fisher Scientific; #EC-833). 5 picomoles of the samples were mixed with formamide loading buffer (FLB) in 1:1 volume ratio. The mixture was denatured at 95°C for 2 minutes and then loaded into the wells. The gel was run in 0.5 × TBE buffer for 45 minutes at 250V. The gel was stained with SYBR Gold for 10 minutes and then imaged with the Typhoon Gel Scanner.

**Ethanol precipitation**

The peptide-oligonucleotide conjugates were purified via ethanol precipitation. After filtering, 100% ethanol and 3M NaOAc were added to the sample in a 50mL Falcon tube, so that a final concentration of 80% ethanol and 0.3M NaOAc was achieved. The solution was placed in the -80°C freezer for a minimum of one hour. The sample was then spun down at high speed in a pre-cooled RC6+ Superspeed Centrifuge (Thermo Scientific). The sample was spun for 16,000g for 1 hour at -20°C. A white pellet was observed after spinning. The pellet was washed twice with 5 mL of 75% ethanol. The pellet was dried at ambient temperature and resuspended in 1 × PBS.

**Neoantigen peptide-conjugated oligonucleotide hybridization with SQB**

The purified peptide-conjugated oligonucleotides were hybridized to the PEG purified SQB. The peptide-conjugated oligonucleotides were added in 2 × excess to the SQB, maintaining a final concentration of 10mM MgCl_2_ and 1 × TE via addition of stock solutions of 10 × TE and 100mM MgCl_2_. The SQBs were added last to ensure that they are not subjected to variable concentrations of TE and MgCl_2_, which can disrupt the structural integrity of the DNA origami_._ The peptide-SQB solution was incubated for 1-2 hours at 37°C while shaking. The purity and integrity of the sample was confirmed via agarose gel electrophoresis and TEM. The peptide-SQBs were purified via PEG purification and their concentration was determined via NanoDrop. The number of peptides conjugated to each SQB was determined via a DNA I degradation assay (below).

**DNAse I degradation and silver stain analysis**

The peptide loaded SQBs were subjected to DNase I degradation. 1 picomole of SQBs was incubated with 1.0 U/µl DNase I (NEB) with 10 × DNase I buffer diluted in water (Gibco). Samples were incubated in the thermocycler for 30 minutes at 37°C. The sample was combined with 4 × NuPAGE LDS sample buffer (ThermoFisher; #NP0008), incubated at 70°C for 10 minutes in the thermocycler, and loaded in 4-12% NuPAGE Bis-Tris gels (ThermoFisher; #NP0322). The gel was run in 1 × MES SDS running buffer (ThermoFisher; #NP0002) at 200V for 32 minutes. The gel was analyzed via silver stain (Pierce, #24612), following the manufacturer’s protocol. The gel was imaged under Silver Stain setting on Image Lab 6 on a Gel Doc EZ Imager (Bio-Rad). ImageJ was used to quantify the band intensity and determine the conjugation efficiency of the peptides to the SQBs.

**Processing LNs**

The axillary and superficial cervical LNs on the left side of the mouse (the same side as the vaccine injection) were collected for flow. The LNs were meshed through a 70 µm cell strainer using a syringe plunger to obtain a single cell suspension. The suspension was separated and stained with various panels of multiple antibodies. The cells were then washed and analyzed using flow cytometry (LSRFortessa, BD).

**Processing blood cells**

Blood cells were collected in heparin-coated tubes to prevent clotting. Plasma was collected by spinning at 3000 rpm for 5 minutes to separate the plasma cells from other blood cells. Plasma was saved at -80 °C before analyzing. The blood cells were treated with red blood cell lysis buffer 3 times. The remaining peripheral blood mononuclear cells (PBMCs) were analyzed using flow cytometry or via ELIspot (LSRFortessa, BD).

**Tumor cell processing into single cell suspension**

Mice were sacrificed on day 15 after tumor inoculation. The melanoma tumors ranged from 2–15 mm in diameter at the time of sacrifice for tumor-cell processing. The tumors were dissociated from the skin and muscle. The tumor tissues were then sliced into 1–2 mm pieces and dissociated in DMEM supplemented with 80 U/mL collagenase type IV (Gibco # 17104019), 20 U/mL DNase I in HBSS (Sigma #11284932001), and 2 mg/mL bovine serum albumin (Sigma, #05470) for 1.5 hours at 37°C, shaking at a slow speed to ensure proper mixing. The resultant cell suspension was filtered through a 70 μm cell strainer to obtain a single cell suspension. The cells were then stained with antibodies in various panels for flow cytometry analysis.

**Anti-dsDNA antibody and anti-PEG antibody test**

The plasma samples collected from various animal tumor model studies were applied to evaluate the anti-dsDNA antibody and anti-PEG antibody in the mouse serum after being treated by different conditions. The anti-mouse dsDNA IgG (#5120), anti-mouse PEG IgG (#PEG-030), and anti-mouse dsDNA IgA/G/M (#5110) ELISA kits were purchased from Alpha Diagnostic. For anti-dsDNA antibody tests, the plasma samples were diluted 100 times. For anti-PEG antibody tests, the plasma samples were diluted 10 times. The experiments were performed following the detailed instruction provided by the manufacturer.

**Extended statistical analyses**

The linear mixed effect model for tumor volume trajectory in figure 5B was estimated using a random intercept for each mouse, and fixed effect estimates for time, treatment modalities and their interaction. The intercept is defined as the control group starting at day 0. Day is treated as continuous and is highly significant. It modifies the effect of the control across time. All other arm variables represent the intercept for the growth rates of various modalities of treatment. The intercepts are not significantly different from the controls. The interaction variables, between Day and Arm measure the trajectories of that modalities rate. Here the trajectory growth rate for 3.5nm (CpG2) is significantly lower than the control, and show the greatest reduction in tumor volume across time when compared to the control. Irregular Spacing (CpGi) & Free CpG2+Free OVA are both statistically different from the control, and show a greater reduction in tumor volume across time than the control. The statistical difference between CpG2-SQB+SQB-OVA & 7nm (CPG4) and the control is marginally significant, with both arms responding better than the control. If we change the reference arm from control to CpG2, the growth of every other treatment is larger than the CpG2 whose growth rate is now dictated by day and its intercept (Table S5). The trajectory growth rate for CpG2 is significantly slower than CpG4 and CpGi. For other tumor growth results with less groups involved, two-way ANOVA was applied with Tukey's multiple comparisons test.

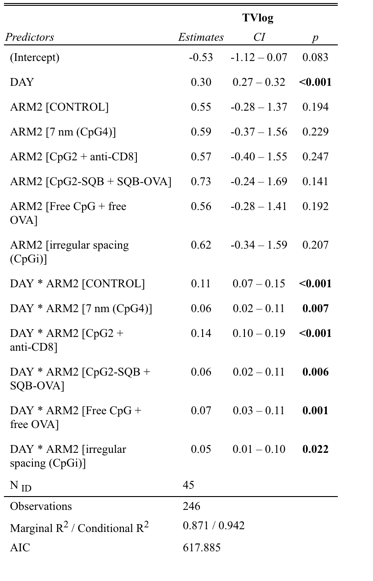

Table S5. Statistics results for Fig. 5B tumor trajectory growth when using CpG2 as the reference arm.
